# The landscape of allele-specific expression in human kidneys

**DOI:** 10.64898/2025.12.03.692205

**Authors:** Ana C. Onuchic-Whitford, Junmo Sung, Eric D. Sakkas, Michelle T. McNulty, Christopher L. O’Connor, Anya Greenberg, Jihoon G. Yoon, Sowmya Badina, NEPTUNE (Nephrotic Syndrome Study Network), Laura H. Mariani, Markus Bitzer, Matthew G. Sampson, Dongwon Lee

**Affiliations:** Division of Nephrology, Brigham and Women’s Hospital; Boston, 02115, USA; Division of Nephrology, Boston Children’s Hospital; Boston, 02115, USA; Harvard Medical School; Boston, 02115, USA; Broad Institute of MIT and Harvard; Cambridge, 02142, USA; Department of Medicine, University of Michigan; Ann Arbor, 48109, USA; Department of Laboratory Medicine, Yonsei University College of Medicine; Seoul, 03722, Republic of Korea; Department of Laboratory Medicine, Gangnam Severance Hospital; Seoul, 06273, Republic of Korea; Centre for Digital Health and Precision Medicine, The Apollo University; Chittoor, 517127, India

## Abstract

Allele-specific expression (ASE), the preferential expression of one gene copy, is a key mechanism of genomic regulation. However, its role in human kidney disease remains poorly understood. In this study, we generated a high-quality, genome-wide ASE map using paired whole-genome sequencing and RNA-seq from microdissected glomerular (GLOM) and tubulointerstitial (TUBE) compartments of patients with proteinuric kidney disease. We showed that the majority of common ASE events were deterministic and sequence-mediated. We also found that diseased kidneys exhibited significantly more ASE in GLOM than TUBE, compared to controls. Unexpectedly, higher ASE in GLOM than TUBE was significantly associated with improved kidney disease outcomes in the disease cohort. Differential gene expression analysis suggested this was the result of an active, protective transcriptional response, including ribosome and ATP synthesis upregulation, rather than pathogenic dysregulation. Our work reveals glomerular ASE as a marker of adaptive transcriptional activity in proteinuric kidney disease.

## Introduction

Humans are diploid organisms and our cells carry two copies of each autosomal gene, which can be differentiated by heterozygous loci^1^. The contribution of each gene copy, or allele, to a gene’s total expression is not always balanced, and predominant expression of one allele over the other constitutes the phenomenon of allele-specific expression (ASE), or allelic imbalance^2^. ASE results from multiple regulatory mechanisms, with a large heritable component, but is also impacted by non-genetic factors^3,4^. Thus, ASE analysis is a versatile method to assess genomic and epigenetic phenomena, including, non-coding cis-regulatory variants, genomic imprinting, and alternative splicing^5^.

As an intra-individual analysis, ASE minimizes batch effects and environmental confounders to uncover true cis-acting regulatory mechanisms and rare variant effects^6^. However, ASE analysis is subject to technical biases requiring rigorous quality control (QC), particularly regarding haplotype phasing, genotyping, and RNA read mapping^7^. Errors or biases in any of these steps can result in substantial inaccuracies difficult to correct at a cohort level, especially when analyzing rare variants^1,7–10^. Several tools exist for ASE estimation to address some of these issues, but accurately quantifying ASE remains technically challenging^1,11^.

Prior studies for ASE analysis have often relied solely on RNA sequencing (RNA-seq) or on single nucleotide polymorphism (SNP) arrays for genotyping^12,13^. Although RNA-seq can be used alone for both genotype calling and expression quantification, extreme ASE can result in heterozygous sites being miscalled as homozygous^1^. This is a well-described caveat of RNA-only ASE analysis and impedes identification of strong allelic imbalance, including monoallelic expression. SNP arrays, conversely, require imputation that can introduce genotype errors^7^.

Thus, whole-genome sequencing (WGS) is considered the gold-standard genotyping strategy in ASE analysis, as it enables more accurate phasing, higher confidence heterozygous variant calling, minimization of imputation, and assessment of rare variants^6,8^.

ASE has been extensively studied in diseases such as cancer, neurodevelopmental disorders and heart disease, with demonstrated instances of causality^5,14–19^. However, ASE evaluation in kidney has previously focused on non-diseased tissue either from tumor nephrectomies^12,13,20^ or post-mortem samples^21^. Furthermore, datasets such as GTEx^22^ do not account for the complexity of kidney structure, which comprises two distinct functional compartments: the glomeruli and tubulointerstitium^23^. As glomeruli comprise only a small fraction of the total cell count in the kidney cortex^24^, glomerular transcripts are much less abundant in bulk tissue than those of the tubulointerstitium. This makes it challenging to study ASE of glomerular-specific genes. Such compartmental resolution is particularly important when studying proteinuric diseases, characterized by loss of large amounts of protein in the urine due to defects in glomerular filtration^25^. The most severe form of proteinuric disease is the nephrotic syndrome, with loss of ≥3.5 grams of urinary protein daily, resulting in hypoalbuminemia and swelling of the face and extremities. Proteinuric disorders account for up to 90% of end-stage kidney disease (ESKD) worldwide^26^. Specifically, a common mechanism is injury to glomerular podocytes (specialized cells which comprise only ∼1% of kidney cortex), and most of the >70 monogenic causes of proteinuria involve genes expressed in this cell type^27^. However, many patients with glomerular disorders remain undiagnosed, and several known genetic etiologies lack a clear mechanistic pathway to disease^28,29^.

The goal of this study was to discover the genome-wide landscape of ASE in the human kidney and its role in the pathogenesis and progression of proteinuric kidney diseases (**Fig. 1A**). We use a unique dataset of WGS with paired RNA-seq from microdissected compartments of kidney biopsy from the Nephrotic Syndrome Study Network (NEPTUNE)^30^. We create a rigorous, generalizable framework for ASE analysis, integrating both available and novel computational tools. Our pipeline includes new steps to address major sources of bias in ASE, including overlapping genes and concealed genotyping errors. We generate a high-confidence ASE dataset and characterize distinct ASE patterns in glomeruli (GLOM) and tubulointerstitium (TUBE) of diseased kidneys, compared to controls. We then perform extensive investigation into the molecular mechanisms, cell-type specificity, clinical correlations, and differential gene signatures linked to ASE in the human kidney.

**Fig. 1.**
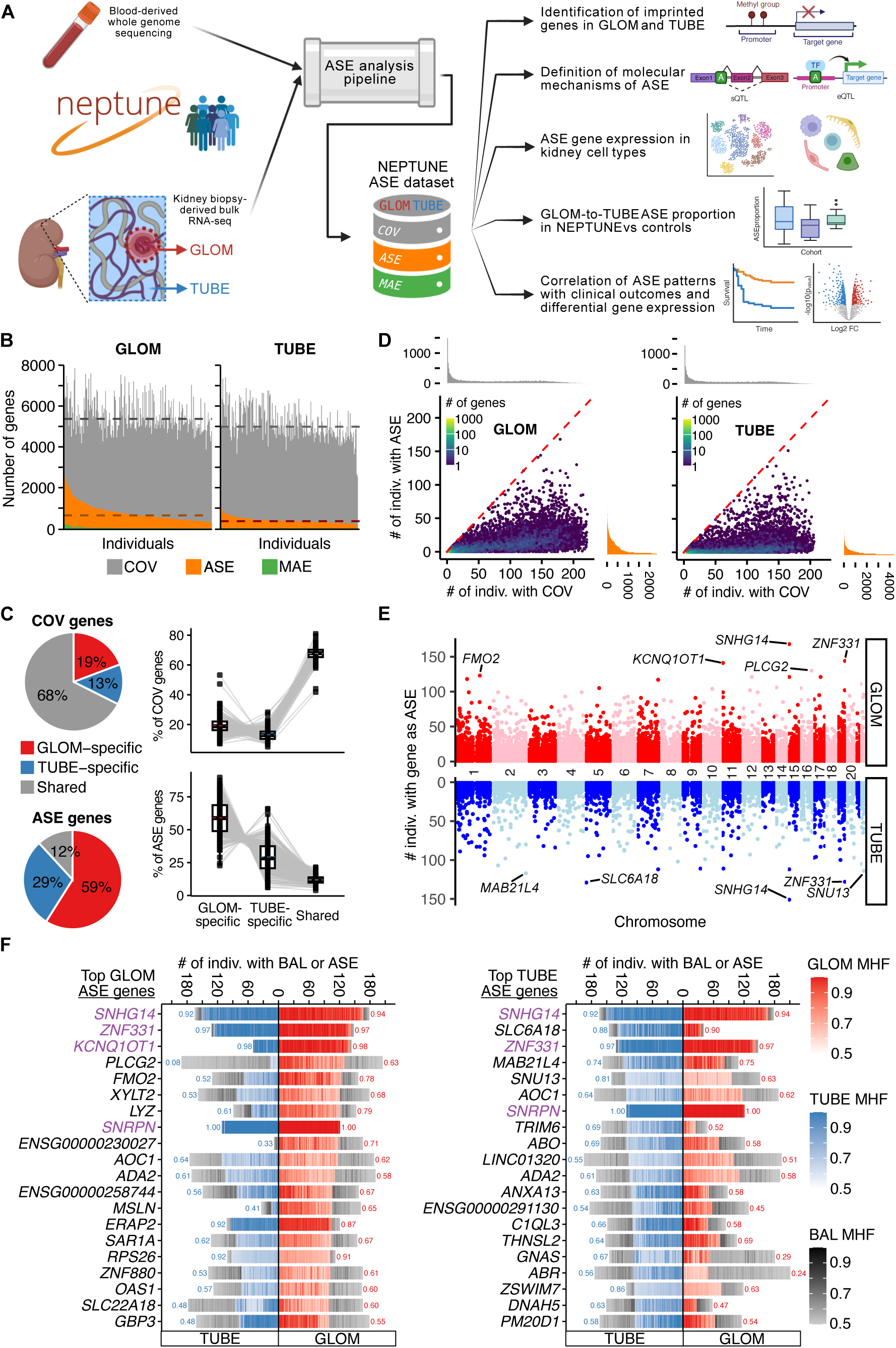
Study design and overview of ASE in kidney glomerular and tubulointerstitial compartments in NEPTUNE. **(A)** Schematic overview of our study. (**B**) Individual-level data: Each bar represents one individual, ordered by number of ASE genes; stacked bars are cumulative. Dashed lines indicate the median of COV (grey) and ASE (orange) genes per sample. (**C**) Overlap of COV (top) and ASE (bottom) genes between GLOM and TUBE in paired samples (n=206). Pie charts show the mean proportions of shared and compartment-specific genes across all individuals. Boxplots show the distribution of these proportions. (**D**) Gene-level data: each dot represents a gene, with the number of individuals of this gene as COV (x-axis), compared to that of ASE (y-axis). The heatmap represents point density to indicate overlapping data points. Marginal histograms are shown on both axes. (**E**) The number of individuals sharing an ASE gene (y-axis) is plotted (GLOM: top, TUBE: bottom) against its genomic coordinate (x-axis). The five most common ASE genes are labeled. (**F**) The top 20 ASE genes in GLOM (left) and TUBE (right), ordered by number of ASE individuals of this gene. The horizontal axis shows the number of ASE and BAL individuals in GLOM or TUBE. Individual stacked bars have color grading determined by major haplotypic fraction (MHF) and sorted by the binomial test *P* value. Imprinted genes (from our imprinting analysis, Fig. 2A) are indicated in purple. The proportion of individuals with ASE among those with COV is shown as the number adjacent to each bar. Indiv, individual.

## Results

### A new workflow for generation of high-confidence ASE data

We assembled a rigorous pipeline for high-confidence ASE calling in kidney tissue from the NEPTUNE proteinuric kidney disease cohort (**Methods**). Biopsies from these individuals were microdissected into GLOM and TUBE compartments, followed by bulk RNA sequencing. We identified 240 NEPTUNE individuals with available paired WGS and GLOM RNA-seq, of which 218 also had TUBE RNA-seq.

Our ASE analysis workflow is summarized in **Fig. S1** and described in detail in **Methods**. Here, we highlight three key features introduced in this work: 1) novel QC steps to blacklist variants prone to genotyping errors; 2) bias reduction from misassigned RNA reads in loci containing overlapping genes; and 3) ASE definition using multiple criteria to minimize false positive calls.

Briefly, we first performed WGS-based genotyping using the GATK best practices pipeline, followed by genotype phasing using WhatsHap^31^ and SHAPEIT4^32^. Both tools employ read-backed phasing, an advantage of WGS, and SHAPEIT4 allowed us to incorporate haplotype data from ∼500,000 participants of the UK Biobank^33^ to leverage population-level phasing information. As detailed below, our estimated gene-level phasing error (flip) rate was low (∼3%), enabling accurate ASE quantification even for genes whose haplotypes were not fully connected by RNA-seq reads.

To ensure high-confidence genotype calling, we also implemented multiple WGS QC measures, excluding variants with: 1) inconsistent genotype before and after phasing; 2) biased DNA allelic fraction; or 3) abnormal DNA read depth. These variants showed a significant deviation of DNA (or RNA) allelic fraction from the expected 0.5, suggesting poor genotyping or structural variation (**Fig. S2**). In addition, we blacklisted variants with bi-allelic RNA expression at homozygous WGS genotypes, suggesting systematic poor read alignment in RNA-seq (**Fig. S3**). Altogether, we removed an average of 543 variants per sample. These blacklisted variants showed a substantially higher proportion of variant-level ASE than the QC-passed variants, consistent with false-positive ASE from erroneous genotyping (**Fig. S4**). Per sample, an average of 302 genes harbored these problematic variants, and thus would appear to have significant ASE without this filter. Our QC strategy also identified and excluded 18 problematic samples with an excess number of such problematic variants (n ≥ 1,000) (**Fig. S2D**). Our findings were further confirmed by verifyBamID analysis^34^, which flagged 13 of the 18 same samples as having DNA-RNA mismatches. This resulted in a final number of 222 GLOM and 206 paired TUBE samples (**Table S1**).

Next, to correct for allelic mapping bias in ASE analyses, we performed RNA-seq read alignment using STAR (Spliced Transcripts Alignment to a Reference) with WASP (Workflow for Allele-Specific Processing)^9,35,36^ and confirmed the same benefits in our dataset (**Fig. S5**). WASP-passing reads, taken only from transcripts overlapping heterozygous loci, represented∼9% of total reads per sample. We then performed ASE analysis using phASER (phasing and Allele Specific Expression from RNA-seq)^6^, incorporating the individualized blacklists described above. phASER employs read-backed phasing from RNA-seq to optimize phasing of rare variants, and accurately phases variants over longer distances due to RNA splicing. In addition, phASER reports gene-level ASE as the number of reads mapping to gene haplotypes, rather than the sum of a gene’s variant-level reads. For downstream analyses, we included genes with sufficient coverage (total haplotypic count ≥20, based on prior literature standards^12,37^), and designated these as “COV” genes. Consistently, at lower coverage cutoffs, our data showed inflated ASE and monoallelic expression (MAE, defined as expression of only one haplotype) (**Fig. S6**).

A major limitation in standard ASE pipelines is that RNA reads aligned to heterozygous variants are assigned to all genes spanning that genomic position, affecting ∼48% of expressed genes in our dataset. To address this, we developed TOGA (Transcript reassignment for Overlapping Genes in ASE), a novel pipeline that resolves these overlapping genes (**Fig. S7; Methods**). Briefly, we first estimated the proportion of allelic counts originating from the target gene relative to overlapping ones; if interference were negligible, the target gene was designated as “resolved.” If not, we recalculated haplotypic counts using reads from non-overlapping regions only. If sufficient coverage remained after this filtration, the gene was also resolved; otherwise, it was “excluded” or remained “unresolved” with its original haplotypic count, depending on its coverage (**Fig. S8**). This strategy successfully resolved 75% of expressed genes per individual (with only 4% unresolved, and 21% excluded), substantially reducing false-positive ASE and elucidating the gene responsible for ASE in overlapping regions (**Fig. S9; Table S2**). Going forward, we kept only “resolved” and “unresolved” genes in downstream analyses—opting to include the latter to avoid missing potential biological signals. However, “unresolved” genes rarely exhibited ASE due to diluted signals from multiple genes.

To quantify the magnitude of ASE, we defined the term “Major Haplotypic Fraction” (MHF): the fraction of reads from the higher-expressed haplotype, with values ranging from 0.5 (balanced) to 1.0 (monoallelic). We defined a gene as having significant ASE as follows: (1) sufficient RNA coverage for high-confidence ASE analysis, i.e., post-TOGA COV genes; (2) binomial test with *P_adj_* < 0.05; and (3) MHF ≥ 0.6 (i.e., one haplotype expressed ≥50% more than the other). This MHF threshold was implemented to address known overdispersion in ASE data, which is not managed by the binomial test alone; a similar approach and cutoff have been described previously^3,7^. **Fig. S10** shows an illustrative ASE analysis plot in one NEPTUNE individual.

### The landscape of ASE in human kidneys with proteinuric disease

In our ASE analysis, we achieved extensive coverage of heterozygous variants (**Table S3**), resulting in a median of 5,370 genes in GLOM and 4,989 in TUBE amenable to ASE analysis (COV genes) per individual (**Fig 1B, Table S4**). Cumulatively across NEPTUNE GLOM and TUBE, we observed 18,304 COV genes in ≥1 individual, of which 14,619 (∼80%) were protein-coding—thus, most genes in the human genome were covered. While the number of COV genes was similar between compartments, the ASE patterns were distinct. We identified a median of 643 GLOM ASE genes per individual, nearly double the median of 361 TUBE ASE genes (**Fig 1B, Table S4**). Moreover, while COV genes were largely shared (mean 68%) between GLOM and TUBE compartments in an individual, only a mean of 12% of ASE genes were similarly shared. Thus, ASE was predominantly compartment-specific, with more than double the proportion of GLOM-specific compared to TUBE-specific ASE (59% vs. 29%) (**Fig. 1C**).

Monoallelic expression (MAE) genes were rare and similarly distinct, with a median of 14 GLOM and 8 TUBE MAEs per individual. Collectively, we identified 3,575 GLOM and 968 TUBE MAEs in ≥1 individual (**Table S4**). All ASE genes are reported in **Table S5**.

COV and ASE distribution per gene is shown in **Fig. 1D**. Most COV genes (∼66% in GLOM, and ∼67% in TUBE) were shared by ≥10% of individuals. However, concordant with prior studies^19^, ASE was largely private. Shared ASE genes followed an exponentially decreasing distribution (**Fig. 1D**, **Fig. S11**), with the majority having ASE in few individuals (≤8 in GLOM, and ≤3 in TUBE). In fact, only 13.7% (GLOM) and 6.7% (TUBE) of ASE genes had ASE in ≥10% of individuals, and we later find that these genes share certain molecular mechanisms of ASE, such as imprinting and strong eQTL effects. Overall, this indicates kidney ASE patterns are predominantly individual-specific.

ASE events were identified throughout the genome in both GLOM and TUBE, without statistical enrichment in any specific genomic region, except for depletion of the excluded chromosome 6 HLA regions (**Fig. 1E**). The 20 most common ASE genes in GLOM and TUBE were largely distinct, as only 5 were shared (**Fig. 1F**). Known imprinted genes were among the most frequent ASE genes and tended to have very high MHF, as imprinting involves epigenetic silencing of one allele. When ranking ASE genes by the proportion of ASE among COV individuals, even more imprinted genes were found among the top 20, indicating that ASE will often be detected for imprinted genes with sufficient coverage (**Fig. S12**).

### Identification of imprinted genes in human proteinuric kidneys

Given the enrichment of imprinted genes among the top ASE genes (**Fig. 1F; Fig. S12**) and their known tissue specificity^37^, we sought to systematically identify imprinted genes within our dataset. To do this, we adapted an established pipeline^37^ that assigns an imprinting likelihood to every ASE gene. This is based on characteristics distinct from other regulatory mechanisms: monoallelic expression in all individuals where the expressed haplotype varies randomly across individuals due to parent-of-origin effects rather than genotype **(Fig. S13A**). With this pipeline, we initially identified 26 candidate imprinted genes in GLOM (nine not previously identified in any tissue) and 18 candidates in TUBE (five novel) (**Figs. S14-17**).

This method has a potential limitation: the risk of genes with eQTLs of very large effect size (leading to near mono-allelic expression) being classified as imprinted. This can occur if the causal non-coding regulatory variant is heterozygous in most individuals and in low linkage disequilibrium with the coding ASE variants. While the prior study concluded that genes meeting these conditions are extremely unlikely^37^, we found that our novel candidate imprinted genes frequently overlapped with strong eQTLs^38^. Specifically, 7 out of 9 novel candidates in GLOM had significant eQTLs (*P* < 10^-6^), compared to only 1 out of 17 of known imprinted genes. A similar pattern was also observed in TUBE. Therefore, we sought to distinguish true imprinting from strong eQTL effects by implementing an additional step that incorporates genotype data and eQTL effect direction to test which mechanism better explained the observed monoallelic expression (**Methods**). We found that most novel candidates were indeed better explained by strong eQTL effects, including 8 GLOM genes (**Fig. S14**) and 4 TUBE genes (**Fig. S16**). In TUBE, one known imprinted gene, *MEST*, was also better explained by an eQTL (**Fig. S17**).

Adding this step reduced the number of identified imprinted genes to 18 in GLOM and 13 in TUBE (**Fig 2A**). Thus, we ultimately discovered two novel candidate imprinted genes: *IGLV6-57* (immunoglobulin lambda variable 6-57) in GLOM and *GLYATL2* (glycine-N-acyltransferase like 2) in TUBE. We found many known imprinted genes to be uniquely imprinted in each compartment (7 in GLOM and 2 in TUBE), but the absence of imprinting in the other compartment was largely due to insufficient ASE data (i.e., low expression), rather than observed balanced expression.

**Fig. 2.**
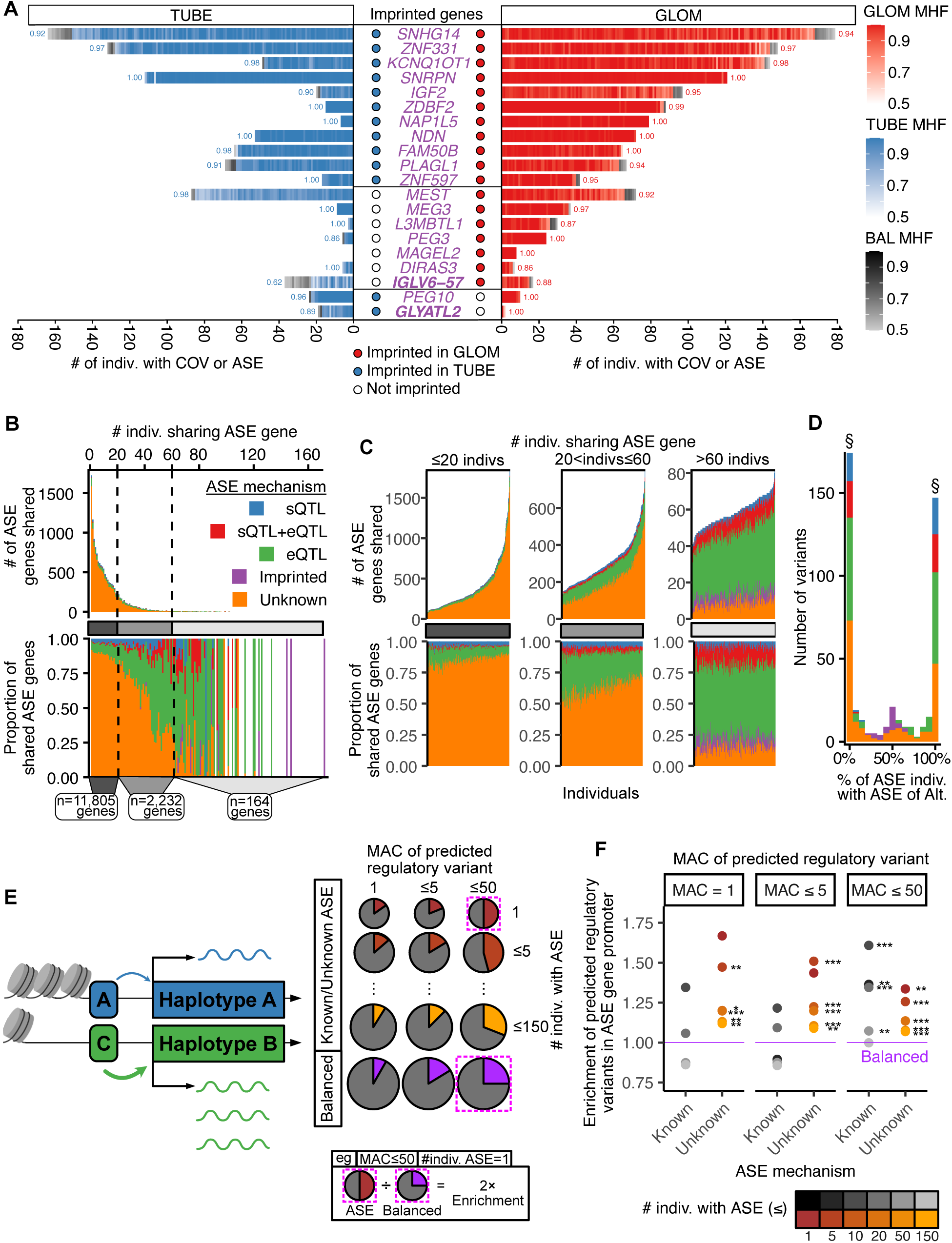
Molecular mechanisms of ASE in NEPTUNE. **(A)** The 20 imprinted genes in GLOM and/or TUBE, identified by our imprinting analysis. Similar to Fig. 1F, bars show the number of individuals with BAL or ASE in GLOM or TUBE, with individual stacked bars color graded by MHF. Circles indicate predicted imprinted status of each gene in the respective compartment, with novel genes shown in bold. **(B)** The top panel is a distribution of the ASE gene frequency across the GLOM samples, stratified and colored by their molecular mechanisms, while the bottom panel shows their proportions. Dashed lines split ASE genes into three groups by number of individuals sharing the ASE gene. The total gene count per group is shown below. **(C)** Each bar represents one individual, stratified by the three ASE gene groups defined in (B). The top panels show numbers of ASE genes, colored by each molecular mechanism, while the bottom panels show their proportions. **(D)** A distribution of the alternative allele preference of ASE variants in GLOM, stratified and color coded by their molecular mechanisms. § indicates perfect allelic preference (PAP) of Ref. (0%) or Alt. (100%). **(E)** Schematic of the promoter variant analysis. **(F)** Enrichment of predicted promoter regulatory variants of ASE gene-individual pairs, comparing known and unknown ASE mechanisms to balanced gene-individual pairs in GLOM. Each dot represents a group of ASE gene-individual pairs, defined by their mechanism (known/unknown), ASE frequency, and the minor allele count (MAC) of the predicted regulatory variants. The y-axis shows enrichment relative to balanced genes (purple line = 1.0). The significance was determined by a one-sided binomial test of the observed count against the expected probability, defined by the proportion of balanced gene-individual pairs with predicted regulatory variants at the corresponding MAC. (\**P* < 0.05, \*\**P* < 0.01, \*\*\**P* < 0.001 ***).

To test tissue specificity of imprinted GLOM and TUBE genes, we compared them to a list of 299 known human imprinted genes (**Table S6; Fig. S18**), which we curated based on literature review^37,39^. Over 85% of these 299 genes did not have ASE data in GLOM or TUBE due to lack of heterozygous variants, low expression, and/or removal by TOGA. However, most of the known imprinted genes amenable to our ASE-based imprinting analysis still did not demonstrate an imprinting pattern (21 of 38 in GLOM, and 30 of 43 in TUBE). Instead, these genes showed biallelic expression (**Fig. S19**), suggesting loss of imprinting in a tissue-specific manner—consistent with prior studies^37^.

### ASE is primarily sequence-mediated

Alongside imprinting, ASE is often driven by sequence-mediated mechanisms, such as eQTLs and splicing QTLs (sQTLs)^1,3^. Thus, we sought to classify every ASE event in each individual by identifying heterozygous eQTL/sQTLs associated with the corresponding gene (**Methods**)^38,40^. In the absence of imprinting, eQTL, or sQTL effects, we assigned its mechanism as “unknown.” We discovered two major patterns. First, the causal mechanism for rare ASE genes (e.g., ASE in ≤20 GLOM) was overwhelmingly unknown, while sequence-mediated or imprinted mechanism drove most of the common ASE genes (e.g., ASE in >60 GLOM) (**Fig. 2B**, **Fig. S20A**). Second, all individuals showed remarkably consistent proportions of their ASE explained by each mechanism (**Fig. 2C, Fig. S20B**). Specifically, the vast majority (mean 86.2%) of the commonly shared ASE genes—which have sufficient power for QTL detection—could be classified into the four categories: sQTLs (4.7%), eQTLs (58.4%), sQTLs+eQTLs (14.5%), and imprinting (8.6%) in GLOM. As a representative example, *PLCG2*—one of the most common GLOM ASE genes (**Fig. 1F**)—demonstrated GLOM-specific, sequence-mediated regulation. In GLOM, *PLCG2* exhibited near-monoallelic expression driven by a strong eQTL, rs4243211 (**Fig. S21**). Conversely, in TUBE, *PLCG2* had no significant eQTL and showed balanced expression, underscoring the compartment-specificity of certain regulatory mechanisms.

To further validate these findings, we evaluated our ASE data at the single-variant level and assessed the consistency of allelic preference across individuals. We first defined an ASE variant based on: (1) variant-level total RNA coverage ≥ 8, (2) allelic imbalance ≥ 50%, and (3) *P_adj_* < 0.05 (binomial test). We restricted our analysis to common ASE variants (i.e., heterozygous genotype in ≥10% of samples, of which ≥50% had ASE) to ensure sufficient power for comparison. For each variant, we calculated the proportion of heterozygous individuals preferring the alternative allele (**Fig. 2D, Fig. S20C)**. Remarkably, most ASE variants showed extreme allelic preference; the majority (e.g., 65.3% in GLOM) demonstrated “perfect allelic preference” (PAP), where the same specific allele (reference or alternative) was consistently preferred in every individual (extreme bars in histograms of **Fig. 2D** and **Fig. S20C**).

Conversely, ASE variants within imprinted genes showed random preference, consistent with its parent-of-origin mechanism. We next identified ASE variants that were also significant eQTL/sQTL SNPs for their respective ASE genes, categorizing these intersections as ASE+eQTL (green), ASE+sQTL (blue), or ASE+eQTL+sQTL (red). As expected, variants in these categories were significantly enriched for PAP, accounting for 72.1%, 92.7%, and 84.8% of variants in GLOM, respectively. Moreover, 61.6% of PAP variants fell into one of these three categories, while none were classified as imprinting (**Fig. 2D**). Overall, similar results were found in TUBE (**Fig. S20C**). Finally, we hypothesized that, if ASE is truly driven by eQTLs, most PAP ASE+eQTL variants should show strong concordance in allele preference between the ASE and eQTL analyses. Indeed, 96.2% of such variants showed allelic concordance in GLOM. Altogether, these results strongly suggest that ASE is primarily sequence-mediated.

We next hypothesized that genes with unknown mechanism may be driven by rare regulatory promoter variants that eQTL analyses are underpowered to identify. Using the ChromKid model^41^, which predicts variant effects on chromatin accessibility in kidney cell types, we analyzed heterozygous promotor variants of all COV genes, in each kidney compartment (**Methods**). We first evaluated promoter variants of balanced (non-ASE) genes, generating a baseline likelihood of identifying predicted regulatory variants in a typical promoter. We then compared ASE genes of known/unknown mechanisms with these balanced genes, calculating an enrichment score (**Fig. 2E**). As expected, genes with an unknown mechanism were significantly enriched for rare (minor allele count, MAC ≤5) predicted regulatory variants, compared to those of known mechanism (**Fig. 2F, Fig. S20D**). This enrichment difference disappeared when including more common (MAC ≤ 50) variants, indicating that rare variants specifically drive unknown ASE. Furthermore, in both mechanistically known and unknown ASE, rare ASE genes (i.e., observed in few individuals) were particularly enriched for these variants, with decreasing enrichment as we included genes with more common ASE (**Fig. 2F, Fig. S20D**). This suggests that while ASE is often driven by rare variants, common ASE genes are driven by a broader variety of mechanisms beyond them.

### Splicing events and phasing errors affect gene-level ASE consistency

Our prior analysis confirmed that sQTLs are a major mechanism driving ASE events. However, phASER’s algorithm represents a potential limitation in identifying sQTL-driven ASE, as it aggregates read counts across all variants of an entire gene haplotype. While this approach works well for eQTLs or imprinting (where the whole transcript is affected uniformly), it masks the complexity of splicing events, in which allelic bias may vary across variants within the same haplotype. This is best illustrated by the example of *OAS1* in **Fig. 3A**. rs10774671-A (alternative allele) disrupts a canonical splice acceptor site, producing an isoform (*OAS1.2*) which retains the full intron^42^. Consequently, this haplotype accumulates reads overlapping intronic variants, such as rs60623134. Conversely, the exonic alleles (such as rs2660) phased with the rs10774671-G (reference allele) will have more reads, as the canonical isoform *OAS1.1* is more stable than *OAS1.2*. Therefore, we hypothesize that a unifying characteristic of sQTL-driven ASE is intragenic discordance of allelic imbalance.

**Fig. 3.**
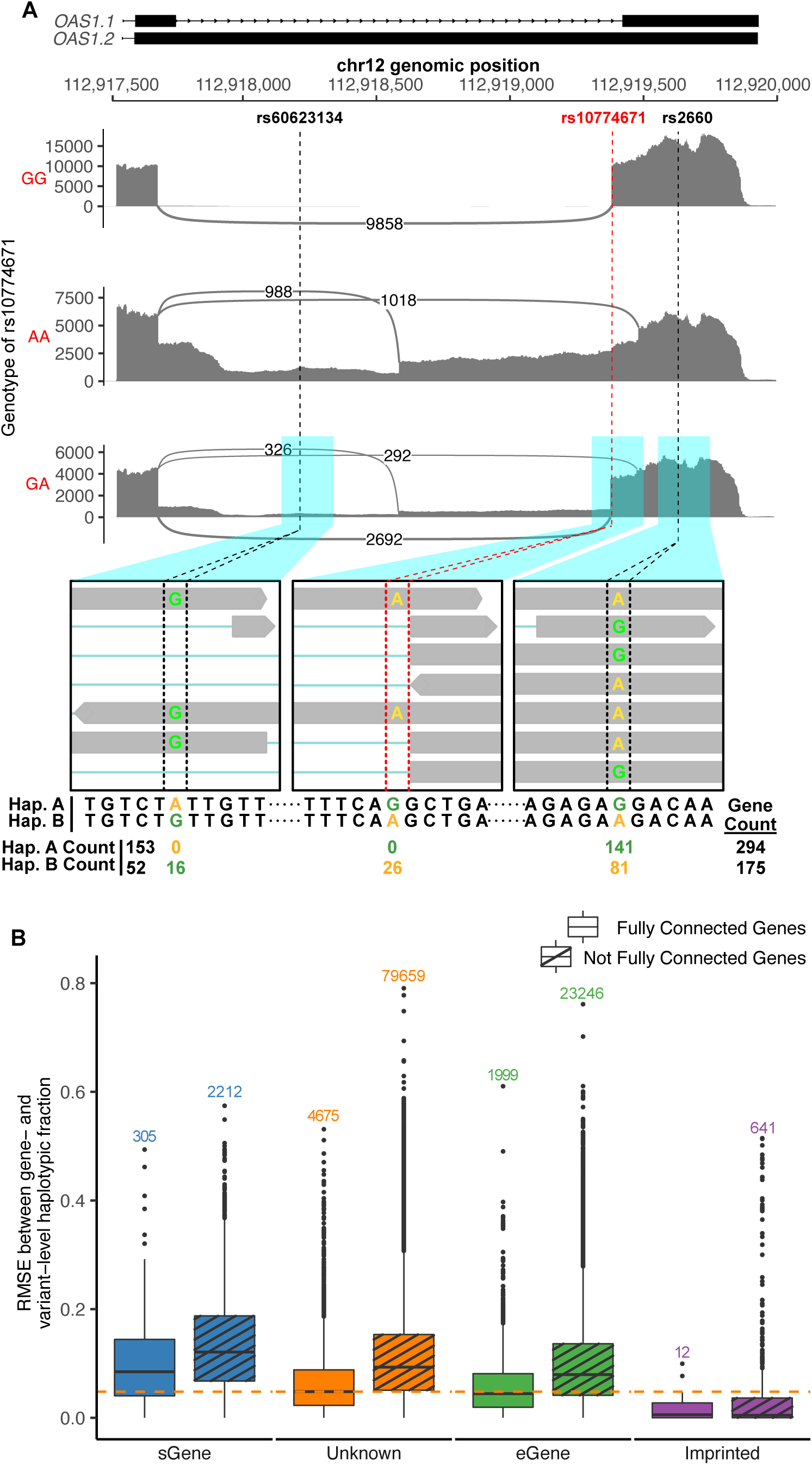
Alternative splicing and phasing errors confound gene-level ASE analysis. (**A**) Sashimi plots demonstrate alternative splicing of the 3’-end of *OAS1* based on sQTL rs10774671 genotype (red), as exemplified in 3 NEPTUNE individuals: one homozygous reference (GG), one homozygous alternative (AA), and one heterozygous (GA). Vertical dashed lines denote genomic locations of rs10774671 and two other SNPs (rs60623134 and rs2660), whose RNA-seq read alignments in the heterozygous individual are shown below the respective Sashimi plot. Variant-, haplotype-, and gene-level ASE counts are shown below the read alignments. (**B**) Quantification of discordance in variant-level and gene-level haplotypic fractions. Gene-level fraction of haplotype A (a randomly chosen haplotype by phASER) was used as the expected value, and the gene’s different variant-level haplotype A fractions were used as the observed values. Genes and respective variants were stratified by ASE mechanism. Gene-individual pairs were further separated into “fully connected genes,” where all the gene’s variants were phased by consecutive overlapping RNA reads by phASER, and those that were not, termed “not fully connected genes”. Numbers above boxplots indicate the number of gene-individual pairs in each category. sGenes whose variants separated by the splicing event were included only. The orange line indicates the median RMSE of the unknown mechanism, fully connected category. Pairwise Wilcoxon tests within each connected group showed that all categories had significant differences in mean RMSE (*P* < 0.001).

To investigate this, we calculated the deviation of variant-level haplotypic fraction from the corresponding gene-level haplotypic fraction, using root mean square error (RMSE) (**Methods**). We hypothesized that sGenes (genes with variant-driven alternative splicing, captured by sQTL analysis) would have higher RMSE values due to the allelic discordance within these genes. Indeed, sGenes had significantly higher RMSE values than genes with other ASE mechanisms (all comparisons Wilcoxon *P* < 0.001), while imprinted genes had the lowest (**Fig. 3B**). RMSEs for ASE genes with unknown mechanism fall between sGenes and eGenes, suggesting that some ASE events of unknown mechanism are driven by sQTLs not found in our dataset. This was largely true for TUBE samples as well (**Fig. S20E**).

Of note, we observed systematically higher RMSE values in genes whose haplotypes were not fully connected by phASER’s RNA read-backed phasing (**Fig. 3B)**. This is indicative of “haplotypic flips” arising from errors in population-based DNA phasing. Imprinted genes provide a ground truth to estimate this error rate: as they are expected to have strictly monoallelic expression, the observation of variants with monoallelic expression mapping to opposing haplotypes within the same imprinted gene confirms a phasing error. We used this principle to estimate the prevalence of haplotypic flips in our dataset. Using all our imprinted genes per individual (i.e., imprinted gene-individual pairs), we calculated the proportion in which at least two variants with read depth ≥ 8 demonstrated conflicting near-monoallelic expression (≥90% allelic bias). In GLOM, we tested 666 imprinted gene-individual pairs with a mean of 3.16 variants per gene, and identified a haplotypic flip rate of only 3%. Similarly, we tested 400 imprinted gene-individuals in TUBE (mean 3.75 variants) and identified a 2.5% rate. This is significantly lower than the previously reported gene-level phasing error rates (∼20%) ^8^, probably due to our enhanced pipeline, further confirming that our gene-level ASE is of high-quality.

### Common ASE genes exhibit cell-type specific expression

We next hypothesized that the ASE genes we discovered are transcriptionally coordinated within specific cell types. Using the Kidney Precision Medicine Project (KPMP) single cell RNA-seq dataset^24^, we classified 54 cell types by compartment and regrouped them into 12 major categories: two unique to GLOM, six unique to TUBE, and four shared between compartments (**Table S7**). We then performed pseudo-bulk aggregation of single-cell RNA-seq data, followed by normalization, to calculate the proportion of each ASE gene’s RNA expression attributable to each cell type within the compartment, termed the “cell-type proportion”. Genes with a cell-type proportion exceeding a defined threshold (i.e., 0.33 for GLOM and 0.2 for TUBE) in one cell type relative to others were classified as having “cell-type predominance” Genes whose expression was broadly distributed across all cell types, without predominance in any single type, were classified as “ubiquitous” (**Methods**).

We initially analyzed all ASE genes observed in ≥1 individual (**Fig. 4A**). In GLOM, 60.1% of ASE genes were ubiquitous; in TUBE, this proportion was 57.4%. However, when we restricted our analysis to more common ASE genes, the proportion of ubiquitous ASE genes decreased while cell-type predominant ASE genes increased. This increase in cell-type predominance was most pronounced for cell types unique to each compartment. For example, among ASE genes observed in at least 30% of individuals, we found increased cell-type predominance for podocytes (POD) and parietal epithelial cells (PEC) in GLOM, and for proximal tubule cells (PT) in TUBE (**Fig. 4A, B**). Compared to all ASE genes, POD-predominant ASE genes increased from 8.4% to 17.2%, and PEC-predominant genes increased from 13.2% to 21.4%, while ubiquitous genes decreased from 60.1% to 35.4%. In TUBE, PT-predominant ASE genes increased from 10.0% to 35.3%, and ubiquitous genes decreased from 57.4% to 33.6%. These findings demonstrate that while rare ASE genes are expressed across all cell types, common ASE genes tend to exhibit more compartment-and cell-type-specific expression patterns within GLOM and TUBE.

**Fig. 4.**
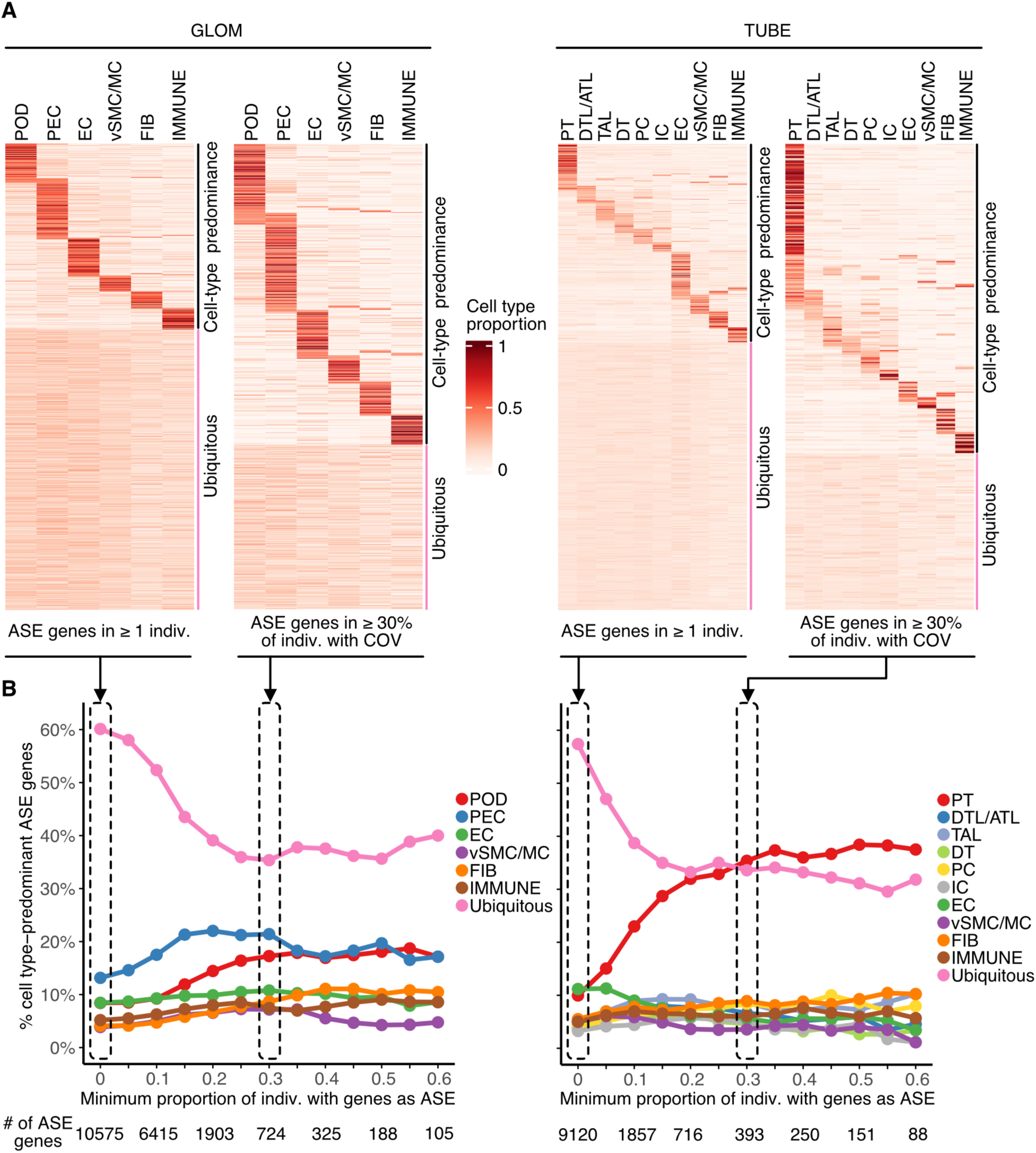
Cell-type specificity of ASE genes in NEPTUNE kidney compartments. (**A**) Heatmaps of cell-type proportions for all ASE genes (ASE observed in ≥1 individuals) and common ASE genes (ASE observed in ≥30% individuals with COV) in GLOM (left) and TUBE (right). Rows represent individual ASE genes; columns represent cell types; color scale indicates cell-type proportion. The right panel of each heatmap indicates whether ASE genes are “cell-type predominance” or “ubiquitous”. All analyses were restricted to genes with COV in ≥10% individuals. **(B)** Trends in cell-type predominance of ASE genes in GLOM (left) and TUBE (right) as a function of ASE frequency. The x-axis is the minimum proportion of individuals with ASE among individuals with COV, and the y-axis is the percentage of ASE genes dominantly expressed in a given cell type. Colors represent cell types in each compartment, and numbers below the x-axis denote the total number of ASE genes meeting the corresponding threshold. The dotted black box corresponds to the heatmap in (A). Unique GLOM cell types are podocytes (POD) and parietal epithelial cells (PEC). Unique TUBE cell types are proximal tubule cells (PT), intercalated cells (IC), distal convoluted tubule cells (DT), thick ascending limb cells (TAL), descending/ascending thin limb cells (DTL/ATL), and principal cells (PC). Common cell types include endothelial cells (EC), vascular smooth muscle/mesangial cells (vSMC/MC), fibroblasts (FIB), and immune cells (IMMUNE).

### Proteinuric kidneys have more genome-wide ASE in GLOM than TUBE

A notable observation in NEPTUNE was that GLOM samples had more ASE genes than TUBE (**Fig. 2**). We therefore calculated the genome-wide ASE proportion per sample (i.e., a sample’s number of ASE genes divided by their number of COV genes). The median ASE proportion in GLOM was indeed significantly higher than TUBE (11.3% vs 7.0%, *P* < 2.2 × 10^-16^), with 177 of 206 (85.9%) individuals showing a higher ASE proportion in GLOM (**Fig. 5A**).

**Fig. 5.**
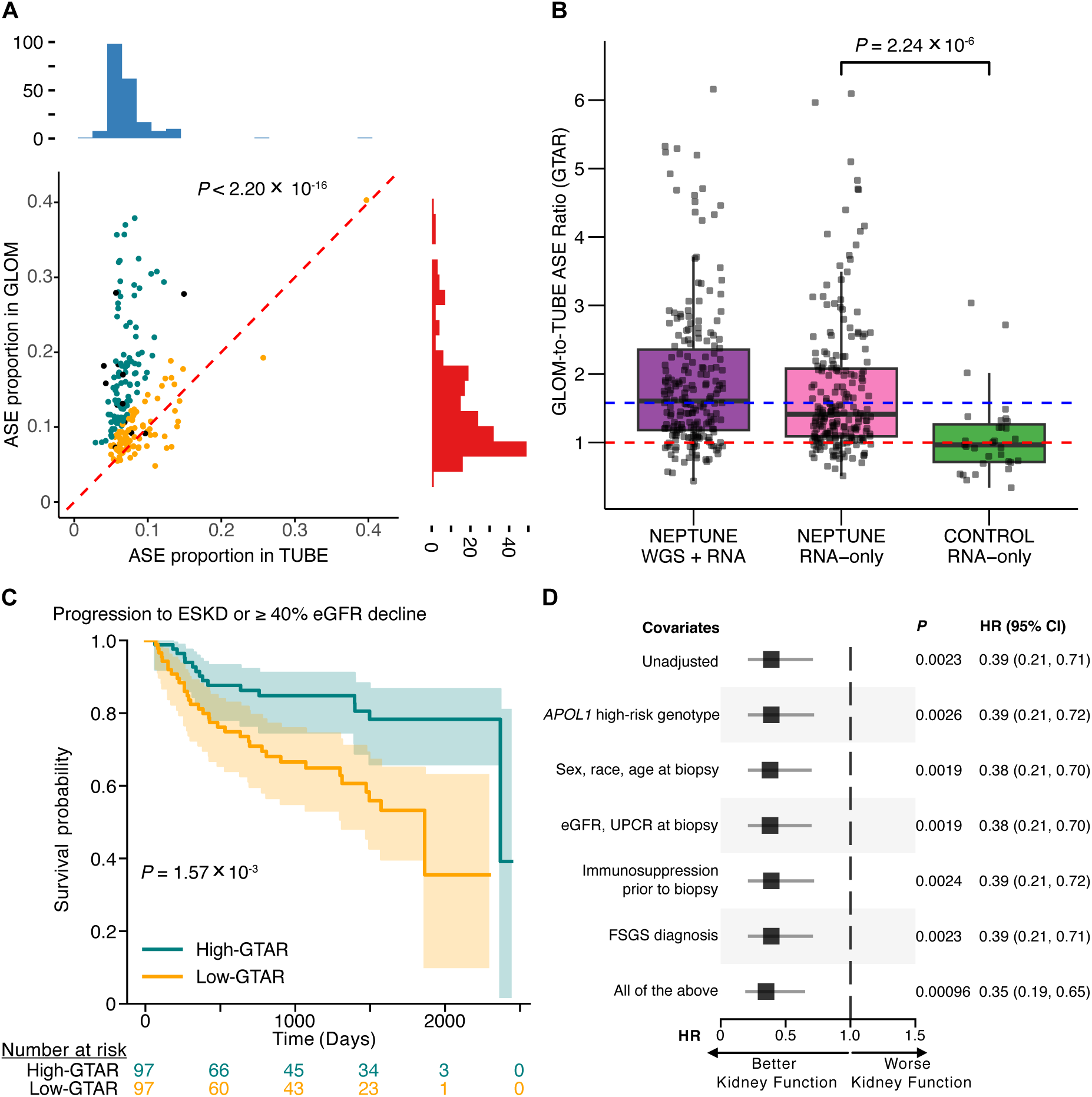
Genome-wide ASE proportion in GLOM/TUBE in NEPTUNE versus controls, and correlation of GTAR with clinical outcomes in NEPTUNE participants. (**A**) Scatterplot of genome-wide ASE proportion (ASE/COV per sample) in paired GLOM and TUBE samples. Each dot is a NEPTUNE individual with both GLOM and TUBE ASE data (n=206), plotted by ASE proportion in TUBE (x-axis) and in GLOM (y-axis). After dichotomizing into two groups for survival analysis, high-GTAR samples (n = 97) are indicated in teal and low-GTAR samples (n = 97) in yellow. Samples without endpoint data are in black (n = 12). Marginal histograms illustrate the distribution of genome-wide ASE proportion in each compartment. The Wilcoxon signed-rank test *P* is shown to compare GLOM and TUBE proportions. The red dashed line indicates equal GLOM and TUBE ASE proportions. (**B**) GLOM-to-TUBE ASE ratio (GTAR) per individual, stratified by cohort and ASE analysis pipeline. Each dot is an individual with paired GLOM and TUBE data (NEPTUNE n=206, control n=31). The blue dashed line represents the median GTAR for NEPTUNE (WGS + RNA pipeline), and the red dashed line indicates GTAR = 1. **(C)** Kaplan–Meier survival analysis of NEPTUNE individuals (n = 194), comparing the high-and low-GTAR groups. Endpoint is composite of ESKD or ≥ 40% eGFR decline. **(D)** Forest plot of Cox proportional hazards model, with hazard ratio (HR) in reference to high-GTAR group, and adjusted for multiple kidney disease-related covariates. ESKD, end-stage kidney disease; eGFR, estimated glomerular filtration rate; UPCR, urine protein-to-creatinine ratio; FSGS, focal segmental glomerulosclerosis.

To determine whether this discrepancy was specific to proteinuric disease or a normal phenomenon of kidney subcompartments, we additionally analyzed 31 microdissected “control” kidneys with preserved kidney function and without proteinuric disease (**Methods**). As these control samples had only bulk RNA-seq (without paired WGS), we developed an RNA-only ASE pipeline for direct comparison (**Methods; Fig. S22A**). We note that, in RNA-only ASE analysis, true (near)-monoallelic expression is missed as these sites are assumed to be homozygous, resulting in fewer identified ASE events and no monoallelic expression compared to WGS+RNA pipeline (**Table S4; Fig. S22B**). Nonetheless, results from the RNA-only analysis of NEPTUNE were highly consistent with the WGS+RNA pipeline (**Fig. S22C**), demonstrating significantly higher ASE proportion in GLOM vs TUBE across the cohort (median 7.67% vs.

5.27%, *P* < 2.2 × 10^-16^) **(Fig. S23A)**. In contrast, the control cohort showed no significant difference (median ASE proportion 6.48% in GLOM vs. 6.96% in TUBE, *P* = 0.68) (**Fig. S23B, C**). Due to baseline differences in the NEPTUNE and control cohorts’ biology and data (i.e., read depth, ancestry, age, and number of COV genes), we normalized the ASE proportions by calculating a GLOM-to-TUBE ASE proportion ratio (GTAR). In the RNA-only analysis, the median GTAR in NEPTUNE was significantly higher than the controls (1.40 vs 0.97; *P*=2.2 × 10^-6^) (**Fig. 5B**).

In light of this difference, we hypothesized that clinical outcomes in proteinuric kidney disease could vary as a function of GTAR. To test this, we dichotomized 194 NEPTUNE participants into a low-GTAR group (GTAR 0.44 – 1.578, similar to most controls) and a high-GTAR group (1.579 – 6.16), based on WGS+RNA analysis, after excluding 12 participants with missing outcome data (**Fig. 5A**). Kaplan-Meier survival analysis revealed that the high-GTAR group had slower kidney disease progression (composite endpoint of ESKD and/or ≥40% decline in estimated glomerular filtration rate—eGFR) compared to the low-GTAR group (*P* = 1.57 × 10^-3^) (**Fig. 5C**). Cox proportional hazards analysis confirmed this finding: high-GTAR patients were 65% less likely to reach the endpoint after adjusting for multiple kidney disease-related covariates (HR = 0.35, *P* = 9.63 × 10^-4^) (**Fig. 5D**). Of note, there was no difference in survival when stratifying solely based on either GLOM or TUBE ASE proportion (**Fig. S24**).

We next investigated GTAR’s correlation with additional clinical outcomes in NEPTUNE (**Methods**). The high-GTAR group achieved complete remission of proteinuria sooner (Cox adjusted HR = 1.58, *P* = 0.023) (**Fig. S25A**), demonstrating improved renal recovery compared to low-GTAR group. Similarly, high-GTAR individuals showed a trend towards later and less relapse of active nephrotic syndrome, although the difference was not statistically significant, likely due to limited sample size (**Fig. S25B**). In contrast, the low-GTAR group showed non-significantly higher kidney scarring, in the forms of global glomerulosclerosis (*P* = 0.083) and interstitial fibrosis (*P* = 0.051) (**Table S8**). Moreover, six of the seven participants who developed kidney failure were in the low-GTAR group. We found no significant associations with other demographic, clinical, or histologic parameters. Altogether, high GTAR was associated with better kidney outcomes.

### Differential gene expression analysis reveals downregulation of fibrosis genes and upregulation of ribosome and ATP pathways in the high-GTAR group

We next explored whether differences in gene expression patterns in each GTAR group could be driving variation in both ASE proportion and disease progression. To this end, we performed differential gene expression (DGE) analysis between high-and low-GTAR groups, using GLOM or TUBE RNA-seq and DESeq2^43^ (**Methods**). GLOM DGE analysis identified 181 significantly differentially expressed genes (DEGs), with *P_adj_* < 0.05 and absolute log_2_-fold change (log_2_FC) > 0.5. Of these, 115 genes were upregulated and 66 downregulated in the high-GTAR group (**Fig. 6A, Table S9**). In contrast, TUBE analysis demonstrated no significant DEGs. Thus, further analysis was focused on the GLOM DGE results.

**Fig. 6.**
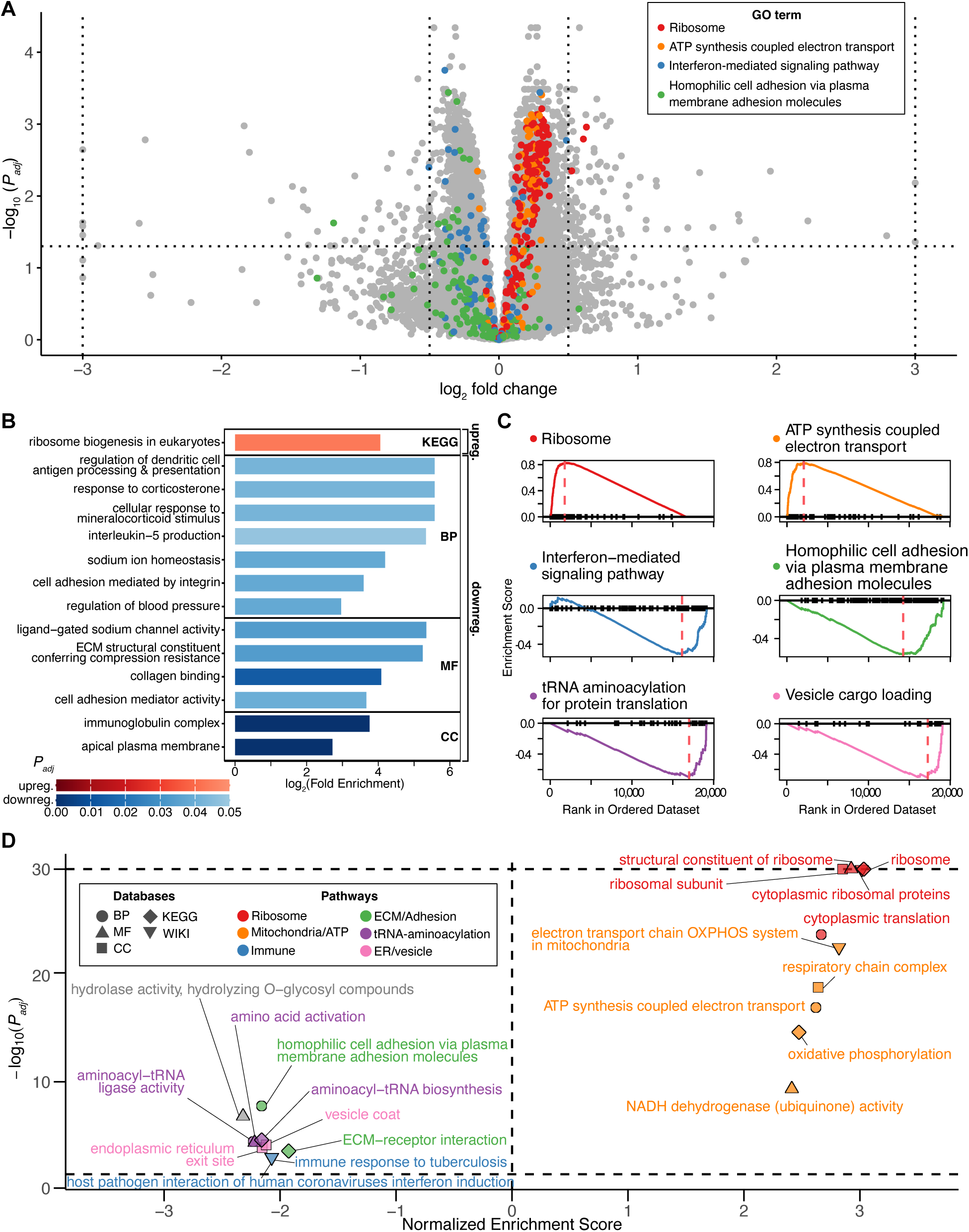
Differential glomerular gene expression between high-and low-GTAR groups and gene set enrichment analyses (GSEA). (**A**) Volcano plot of differentially expressed genes (DEGs) with the low GTAR group as reference. The horizontal dashed line indicates *P_adj_* threshold (0.05), and the inner and outer vertical dashed lines indicate log₂(fold change) cutoffs of ±0.5 and ±3, respectively. Genes exceeding ±3 are aligned with the outer vertical lines. Significantly upregulated GO terms related to ribosome and mitochondrial/ATP synthesis (red and orange) and downregulated immune-related and membrane-associated pathways (blue and green) are highlighted. The corresponding GO terms are shown in *(C)* with matching colors. (**B**) ORA of significantly upregulated (log₂FC > 0.5, *P_adj_* < 0.05) and downregulated (log₂FC < –0.5, *P_adj_* < 0.05) genes across multiple databases. Bars color scales reflect *P_adj_* from ORA (upregulated: red, downregulated: blue). (**C**) GSEA enrichment plots of significant GO terms, grouped into six functional categories: ribosome (red), mitochondria/ATP (orange), immune (blue), ECM/adhesion (green), tRNA-aminoacylation (purple), and ER/vesicle (pink). (**D**) GSEA summary of the top upregulated and downregulated GO terms ranked by normalized enrichment score across multiple databases. Terms with –log₁₀(*P_adj_*) > 30 are truncated and aligned at the top horizontal dashed line. The bottom dashed line indicates the threshold for statistical significance (*P_adj_* = 0.05). The vertical dashed line shows where there is no enrichment. Colors correspond to the six categories in *(C)* and grey color denotes GO terms not included in these categories. Shapes indicate the database source.

Characterization of GLOM DEGs revealed striking biological patterns. First, a high proportion of DEGs (62 of 181; 34.3%) were long non-coding RNAs (lncRNAs). The vast majority (58 of 62; 93.5%) were upregulated, including 20 antisense to protein-coding genes. Five upregulated lncRNAs were antisense to—thus expected to decrease expression of—genes known to promote kidney disease (e.g., *AQP5*, *LYZ*, *P2RX7*, *CERS6* and *NFAT3C*—with latter four specifically causing renal fibrosis)^44–54^. Second, the high-GTAR group had upregulation of several kidney-protective genes and downregulation of many genes which promote kidney injury—a notable finding given the better kidney clinical outcomes in the high-GTAR group.

Specifically, we found upregulation of *FCGR2B* (Fc gamma receptor IIb), an immune inhibitory receptor which suppresses autoimmunity and autoantibody-mediated glomerulonephritis^55–58^, and of *NGF* (nerve growth factor), secreted by podocytes and a protective factor against glomerular kidney injury^59–61^. Conversely, downregulated DEGs were strongly enriched for genes whose increased expression contributes to kidney dysfunction. Of the 66 downregulated DEGs, 10 (15.2%) promote kidney fibrosis (*FN1*, *CCL19*, *CCL21*, *ADAMTS4, ADAMTS16, CEMIP, ITGA11, NEU4* and *HK2*)^62–75^ or cardiac/pulmonary fibrosis (*ASPN*)^76,77^. Several other downregulated genes were associated with glomerular disease (e.g., *MRC2*, *ACER2, IL1RAP, NEFL*)^78–82^. Finally, 7 of 115 (6%) upregulated DEGs were ribosome-related, with no ribosomal genes among downregulated DEGs—a trend strongly supported by subsequent pathway analyses. Detailed characterization of all DEGs in the context of kidney disease biology is summarized in **Table S10**.

We next systematically assessed the 181 GLOM DEGs with over-representation analysis (ORA). For upregulated genes, “ribosome biogenesis in eukaryotes” (KEGG) was the single significant concept (*P_adj_* < 0.05). For downregulated genes, we identified 57 significant terms (many redundant), including “dendritic cell antigen processing and presentation”, “response to corticosterone”, “cell adhesion mediated by integrin”, “ligand-gated sodium channel activity”, “collagen binding” and “immunoglobulin complex” (**Fig. 6B, Table S11**).

To comprehensively identify molecular pathways, we also performed gene set enrichment analysis (GSEA) on the full GLOM transcriptome using clusterProfiler^83^, ranking all genes by signed(log_2_FC) ×-log_10_(*P*) (**Table S11**). Consistent with the ORA results, the top 2 upregulated terms (disregarding highly redundant terms) related to ribosome and oxidative phosphorylation in all 5 functional databases (**Fig. 6D**). The most strongly downregulated pathways involved cell adhesion and extra-cellular matrix interaction, interferon signaling, vesicle transport and endocytosis, and aminoacyl-tRNA biosynthesis. In the TUBE DGE analysis, GSEA did not identify any significantly enriched GO terms.

## Discussion

In this study, we present a comprehensive, genome-wide characterization of allele-specific expression in the human kidney, leveraging a unique dataset of paired WGS and microdissected bulk RNA-seq from the NEPTUNE cohort^30^. By developing and applying a rigorous computational pipeline, we addressed major technical hurdles in ASE analysis, such as overlapping genes and concealed genotyping errors. This allowed us to uncover an accurate landscape of ASE in proteinuric kidney disease. Our primary finding is that, unlike in control tissue, proteinuric kidneys exhibit a significantly higher proportion of genome-wide ASE in GLOM compared to TUBE. We quantified this discrepancy using a new metric, the GLOM-to-TUBE ASE ratio (GTAR), and demonstrated that a high GTAR was strongly associated with improved kidney survival and faster disease remission.

A critical implication of our work is the biological meaning behind the elevated GTAR. The association of high GTAR with better clinical outcomes suggests this imbalance is not a marker of dysregulation or injury. Instead, our differential gene expression analysis between high-and low-GTAR groups points to a more active, protective transcriptional state. We found that the high-GTAR (better outcome) group showed significant upregulation of pathways related to ribosome biogenesis and oxidative phosphorylation (ATP synthesis) in GLOM. This suggests that the observed increase in ASE is a byproduct of elevated transcriptional and metabolic activity, which may confer glomerular resilience to proteinuric injury. This is further supported by the specific DEGs identified; the high-GTAR group exhibited upregulation of known kidney-protective genes and concordant downregulation of a suite of pro-fibrotic and injury-related genes, suggesting an active, adaptive glomerular response.

Our study also provides insights into the molecular mechanisms driving ASE. We demonstrated that most ASE events are individual-specific. However, ASE genes shared across many individuals were predominantly driven by known molecular mechanisms (i.e., eQTLs, sQTLs, and genomic imprinting). More importantly, we provide an explanation for individual-specific ASE, previously speculated as stochastic or random effects. By integrating deep learning models^41^, we revealed that genes with an “unknown” mechanism of ASE were significantly enriched for rare, heterozygous predicted regulatory variants in their promoter regions. This finding strongly suggests that a substantial portion of rare, individual-specific ASE may not be stochastic, but rather a deterministic outcome of an individual’s unique gene regulatory architecture, which eQTL/sQTL studies are underpowered to detect.

The intersection of elevated, disease-specific glomerular ASE and the deterministic nature of its regulation raise a new set of interesting questions. For example, is it possible that the genetic architecture of certain individuals, perhaps through rare regulatory variants, prevents key genes from making an effective transcriptional response, predisposing them to the low-GTAR (poor outcome) state? This also raises the broader question of whether genetic effects on glomerular disease risk are context-specific, only becoming apparent or exerting their pathogenic effect under the physiological stress of glomerular injury. Consequently, it leads to the following important question: to what extent do studies relying on healthy reference datasets miss these critical, disease-specific regulatory mechanisms?

These biological insights were enabled by the development of a stringent and rigorous bioinformatic pipeline designed to generate high-confidence ASE data. We implemented multiple novel QC steps, including detection of discordant genotypes between pre-and post-phasing and systematic genotyping errors by evaluating homozygous samples. Furthermore, we developed TOGA, a new algorithm to resolve read misassignments from overlapping genes, a problem affecting nearly half of the analyzable genes. This rigorous framework, including the use of WASP ^9^ to correct mapping bias and phASER ^6^ for haplotype-level quantification, was critical for accurate ASE quantification.

This study has several limitations. First, the use of short-read RNA-seq can fail to resolve haplotypes across long distances, particularly for genes with few heterozygous variants.

However, we mitigated this by integrating population-level phasing from SHAPEIT4^32^ and RNA read-backed phasing from phASER, which leverages spliced reads to connect distant variants.

The high quality of our phasing is supported by our low estimation of gene-level haplotypic flips (2.5-3%) in known imprinted genes. Second, our primary analysis was performed on bulk RNA-seq from microdissected glomerular and tubulointerstitial compartments. While this provides crucial compartmental resolution, it lacks the granularity of single-cell analysis. We began to address this by integrating KPMP single-cell data^24^, which revealed that frequently shared ASE genes tend to be cell-type specific. Nonetheless, performing ASE analysis at a true single-cell resolution is needed to fully delineate the cell-specific drivers of ASE in kidney disease. Third, our comparison to the control cohort required using an RNA-only pipeline due to data availability, although the stark difference in GTAR was robust. Lastly, our identification of rare variants driving “unknown” ASE relies on computational predictions, and functional validation of these specific promoter variants is necessary.

In conclusion, we provide the first comprehensive landscape of allele-specific expression in kidney tissue in proteinuric disease, revealing widespread transcriptional changes in the glomerular compartments. We establish a new metric, GTAR, which links a higher proportion of glomerular ASE to a protective transcriptional state and improved kidney survival. This study demonstrates that rigorous ASE analysis is a powerful tool for uncovering disease biology and identifying clinically relevant biomarkers in human kidney disease.

## Acknowledgments

This study has benefited from useful comments from members of Sampson lab and Lee lab.

We would like to acknowledge the Boston Children’s Hospital High-Performance Computing Resources BCH HPC Clusters, Enkefalos 3 (E3), made available for conducting the research reported in this publication. Software used in the project was installed and configured by BioGrids. We would also like to acknowledge that Fig.1A was created with BioRender.com.

The Nephrotic Syndrome Study Network (NEPTUNE) is part of the Rare Diseases Clinical Research Network (RDCRN), which is funded by the National Institutes of Health (NIH) and led by the National Center for Advancing Translational Sciences (NCATS) through its Division of Rare Diseases Research Innovation (DRDRI). NEPTUNE is funded under grant number U54DK083912 as a collaboration between NCATS and the National Institute of Diabetes and Digestive and Kidney Diseases (NIDDK). Additional funding and/or programmatic support is provided by the University of Michigan, NephCure Kidney International and the Halpin Foundation. RDCRN consortia are supported by the RDCRN Data Management and Coordinating Center (DMCC), funded by NCATS and the National Institute of Neurological Disorders and Stroke (NINDS) under U2CTR002818.

The results here are in whole or part based upon data generated by the Kidney Precision Medicine Project (KPMP). Accessed June 26, 2025. https://www.kpmp.org. The KPMP is supported by the National Institute of Diabetes and Digestive and Kidney Diseases (NIDDK) through the following grants: U01DK133081, U01DK133091, U01DK133092, U01DK133093, U01DK133095, U01DK133097, U01DK114866, U01DK114908, U01DK133090, U01DK133113, U01DK133766, U01DK133768, U01DK114907, U01DK114920, U01DK114923, U01DK114933, U24DK114886, UH3DK114926, UH3DK114861, UH3DK114915, and UH3DK114937. We gratefully acknowledge the essential contributions of our patient participants and the support of the American public through their tax dollars.

The Genotype-Tissue Expression (GTEx) Project was supported by the Common Fund of the Office of the Director of the National Institutes of Health, and by NCI, NHGRI, NHLBI, NIDA, NIMH, and NINDS.

This research has been conducted using the UK Biobank Resource under Application Number 64945.

## Author contributions

Conceptualization: ACO, DL, MGS

Methodology: ACO, AG, CLO, DL, EDS, JS, JGY, LHM, MB, MTM

Investigation: ACO, CLO, DL, EDS, JS, MB, MTM, SB

Visualization: ACO, DL, EDS, JS, MTM

Funding acquisition: ACO, DL, MB, MGS

Project administration: ACO, DL

Supervision: ACO, DL, MGS

Writing – original draft: ACO, DL, EDS, JS

Writing – review & editing: ACO, AG, CLO, DL, EDS, JS, JGY, LHM, MB, MTM, MGS, SB

## Data and materials availability

Raw WGS and RNA-seq data from GLOM and TUBE are available through NEPTUNE (https://www.neptune-study.org/ancillary-studies). CONTROL GLOM and TUBE datasets are available through Dr. Markus Bitzer and NEPTUNE. Single cell RNA-seq data used to infer cell-type proportions of ASE genes can be accessed from KPMP repository (https://atlas.kpmp.org/repository). eQTL data are available through NephQTL2 (https://www.nephqtl2.org). The GTEx sQTL data used for the analyses described in this manuscript, GTEx_Analysis_v10_sQTL.tar, were obtained from: https://www.gtexportal.org/home/downloads/adult-gtex/qtl on 01/03/25. All code used for study analyses are publicly available in our Github repository (https://github.com/dongwonlee-lab/ASE_NEPTUNE).

## Supplementary Materials

Methods

Figs. S1 to S25

Tables S1 to S11

Members of the Nephrotic Syndrome Study Network

