## Supplemental Text and Figures for "The landscape of allele-specific expression in human kidneys"

**The file includes:**

Materials and Methods
Figs. S1 to S25
Tables S1 to S11
References
Members of the Nephrotic Syndrome Study Network

### Materials and Methods

Nephrotic Syndrome Study Network (NEPTUNE)

NEPTUNE[1] is a multicenter, longitudinal cohort of participants with proteinuric kidney disease, including nephrotic syndrome. Participants are enrolled at the time of their first clinically indicated kidney biopsy, during which an extra core is collected for research. This research biopsy core undergoes histopathologic characterization[2–4] and is also microdissected into glomerular (GLOM) and tubulointerstitial (TUBE) compartments for transcriptomic analysis. Whole genome sequencing (WGS) data derived from blood and RNA-seq data from GLOM and TUBE samples were obtained and analyzed. Race and ethnicity were self-reported or reported by parents of children. Reporting race and ethnicity in the NEPTUNE study was mandated by the US National Institutes of Health (NIH), consistent with the Inclusion of Women, Minorities, and Children policy. For analysis, the “Other” self-reported racial group comprises individuals with Unknown and Not Available (NA).

Processing of NEPTUNE whole-genome sequencing

CRAM files were created from raw WGS data from the NEPTUNE cohort, provided as FASTQ files, by following the Functional Equivalence pipeline[5]. Subsequent processing was performed using the WARP WholeGenomeGermlineSingleSample WDL pipeline (v2.3.3; https://github.com/broadinstitute/warp/), which was run on Cromwell v52. Individual-level gVCF files were produced using HaplotypeCaller from the GATK v4.2.0 suite, with default setting. Joint genotyping was then performed across the cohort using the JointGenotyping WDL pipeline from the Broad Institute with the default parameter set. Following joint calling, variant sites tagged as "ExcessHet" were filtered out. Finally, Variant Quality Score Recalibration (VQSR) filters were applied using the thresholds of VQSR Tranches SNP 99.7 and INDEL 99.0 to create the final VCF files.

Phasing of NEPTUNE WGS

Phasing is the process of separating an individual’s genotype data into two haplotypes, corresponding to the nucleotide sequences specific to each homologous, maternally- and paternally-inherited, chromosomes. When parental genotype data is not available, computational strategies are employed. These include read-backed phasing, which uses sequenced reads to connect variants, and population-based phasing, which uses haplotype frequencies from a reference panel. Here, we employed a hybrid approach combining both methods to maximize phasing accuracy, which is a critical preliminary step for the downstream calculation of haplotypic counts by phASER[6].

NEPTUNE WGS VCF files were initially processed using the WhatsHap tool, which performs read-backed phasing. This approach uses sequencing reads covering two or more heterozygous variants to assemble haplotype blocks, called “phase sets”, which are recorded in the VCF file with a unique PS tag attached to the genotype[7]. Quality control (QC) was then performed to filter out low-quality variants which could be incorrectly imputed during population phasing. We kept variants with GT missingness rate < 5% and HWE exact test $P > 1 \times 10^{-10}$. SHAPEIT4[8] was subsequently used for population-based phasing of the NEPTUNE WGS data in a 2-step approach. We initially ran SHAPEIT4 with the option --scaffold to generate haplotype scaffolds for NEPTUNE original VCFs, using the UKB haplotype panel as a reference. To create this reference panel, the BGEN format per-chromosome genotype files for ~ 500,000 UKB participants[9] were first converted to VCF format using the bgenix tool (enkre.net/cgi-bin/code/bgen/doc/trunk/doc/wiki/bgenix.md), with VCF header correction using bcftools. UKB VCF files were then lifted over from hg19 to hg38 using the GATK LiftoverVcf tool with the option --RECOVER_SWAPPED_REF_ALT. These scaffolds retain variants that are present in the original dataset and in the reference panel, representing a sparser set of variants that incorporate prior phase information of some heterozygous genotypes. We then ran SHAPEIT4 again using the NEPTUNE VCFs with read-backed phase sets as input and incorporating the previously generated individual scaffolds. With this approach, scaffold variants remain unchanged, and all additional variants from each NEPTUNE VCF are phased onto their respective scaffold, resulting in the final population-phased VCF.

Processing of NEPTUNE bulk RNA-seq

We analyzed paired-end RNA-seq data from 222 microdissected GLOM samples and 206 paired TUBE samples from the NEPTUNE cohort. QC of individual sequencing reads was performed using FastQC (http://www.bioinformatics.babraham.ac.uk/projects/fastqc/), assessing parameters such as adapter content, GC distribution, and per-base sequence quality. To detect potential contamination and off-target RNA, we applied FastQ Screen[10], which maps a random subset of reads from each sample to 16 different genomes and known artifacts. RNA-seq reads were subsequently aligned to the hg38 reference genome using STAR v2.7.11b[11]. The WASP algorithm[12] was applied during this process using the --waspOutputMode option, which flag reads exhibiting reference allele mapping bias with a vW tag in the SAM output. WASP (Workflow for Allele-Specific Processing) is a method that addresses read mapping bias by computationally flipping heterozygous sites, remapping the read, and discarding (or flagging) the read if it does not map to the same genomic location. This approach is widely accepted as it significantly improves reference bias and ASE accuracy while removing only a small proportion of heterozygous reads. Following alignment, duplicate reads were marked using samtools markdup v1.19.2 .This tool designates which of the duplicate reads to keep according to base quality scores rather than read mapping scores, as the latter may also contribute to reference bias[13]. Gene quantification for subsequent analyses was then performed using RSEM v1.3.1[14], a robust tool for estimating transcript abundances. As inputs, RSEM was provided with the hg38 reference genome, GENCODE v45 primary assembly for transcript and gene annotations[15], and BAM files generated using STAR's --quantMode TranscriptomeSAM option.

Sample identity and contamination verification of NEPTUNE WGS and RNA-seq

We used verifyBamID[16] to assess sample integrity, specifically checking for cross-sample contamination and potential sample swaps. This tool evaluates the concordance between sequencing reads and known genotypes to estimate contamination arising from sample mixtures. For the WGS data, verifyBamID was provided with a VCF file containing exonic variants filtered using GENCODE v45 annotations[15]. For the RNA-seq data, the STAR-aligned, duplicate-marked BAM files were used as input. This analysis was performed on all 240 GLOM and 218 TUBE samples. The results identified strong evidence of sample mismatches between WGS and RNA-seq data for 13 GLOM and 7 TUBE samples, indicating probable sample swaps. Due to the uncertainty of whether the mismatch originated from the WGS or RNA-seq identifier, we conservatively excluded these samples from both the NEPTUNE GLOM and TUBE cohorts for all downstream analyses.

Generation of individualized phASER blacklists for ASE analysis

High-confidence heterozygous (het) genotyping calls are necessary for valid allele-specific expression (ASE) analysis, as erroneous heterozygous calls can lead to false-positive monoallelic expression. We therefore implemented two new QC strategies to improve ASE calling by filtering suspicious variants beyond the default phASER blacklists[6].

First, we performed individual-level WGS QC to remove low-quality heterozygous variants. For all variants called heterozygous after population phasing, we calculated the fraction of sequencing reads supporting the alternative allele in both WGS and RNA-seq, denoted as DNA and RNA alt fraction. To calculate the RNA alt fraction, we ran phASER using only the default variant blacklists[6]. Variants were added to an individual-specific blacklist as follows. First, we identified variants with discordant genotype between pre- and post-population phasing. Second, we identified variants with extreme sequencing depths (WGS sequencing depth $< 10$ or $\geq 80$). Third, we identified variants with extreme allelic imbalance at the DNA level (DNA alt fraction $< 0.2$ or $> 0.8$). On average, this individual-level filter blacklisted 178 variants per GLOM sample, of which 5.52% had a mismatched genotype, 88.1% were alt fraction fails, and 44.7% were DNA depth fails. Similarly, an average of 184 per TUBE sample were blacklisted, of which 5.02% mismatched genotypes, 86.2% alt fraction fails, and 56.7% DNA depth fails. Note that some variants failed to pass multiple criteria. Furthermore, individuals with >1,000 such QC-failed variants (n=18) were removed from the analysis entirely.

Second, we implemented a cohort-level filter to identify and blacklist variants susceptible to systematic mapping errors, which cause RNA/DNA discordance. For each variant, individuals were grouped by their WGS-derived genotype: homozygous reference (HOM_REF), homozygous alternative (HOM_ALT), or heterozygous (HET). We then used samtools mpileup on the RNA-seq BAM files (prior to WASP-correction) to calculate the average RNA alt fraction for all individuals within each genotype group. Variants were added to a blacklist if the average RNA alt fraction were substantially discordant with the WGS genotype, that is, if the HOM_REF group had an average RNA alt fraction $> 0.1$ or the HOM_ALT group had an average RNA alt fraction $< 0.9$. This filter removed an additional average of 363 variants per GLOM sample and 361 per TUBE sample. After applying all blacklists (default and custom), the average individual had 62,395 high-confidence unique variants available for ASE analysis in GLOM and 65,614 in TUBE.

Quantification of gene-level ASE using phASER

For ASE analysis, we adopted the phASER framework[6]. This framework first outputs ASE data at the variant- and haplotypic-level, which is then aggregated by the phASER Gene AE tool to produce gene-level ASE quantification. We ran phASER using our population-phased genotypes derived from WGS and the WASP-corrected RNA-seq data for both GLOM (n = 222) and TUBE (n = 206) samples. When running phASER, several read-filtering criteria were applied as follows: only reads with a mapping quality $\geq$255 and a base quality $\geq$10 at the heterozygous variant were considered. Also, insertions and deletions (indels) were excluded. Additionally, duplicate reads and any reads that did not map as a proper pair were removed. Lastly, we ran phASER Gene AE to calculate gene-level ASE, using GENCODE v45 (comprehensive gene annotation with primary assembly regions) for gene annotations[15].

To determine which instances of unequal haplotypic expression consisted of significant ASE – i.e., deviated sufficiently from the expected random distribution of allelic counts, we took into account statistical and biological considerations. We only considered genes with total RNA coverage of 20 reads (COV), based on prior literature reporting high ASE accuracy[17,18]. To quantify the magnitude of ASE, we defined the term "Major Haplotypic Fraction" (MHF), calculated as [preferentially expressed (major) haplotypic read count / total haplotypic read count per gene], thus always ranging from 0.5 (balanced expression) to 1.0 (monoallelic expression), similar to Baran et al[18]. We performed binomial testing using actual counts to compare the MHF to the expected null distribution (0.5), followed by adjustment for multiple comparisons with the Benjamini-Hochberg method. Significance level of $P_{adj}$ was set at < 0.05. The binomial test has limitations in the setting of ASE, given known overdispersion of allele-specific data due to biological and technical factors[19]. Nonetheless, it remains the most widely used statistical test for ASE analysis[6,19,20], with alternate methods such as beta-binomial resulting in loss of true ASE signals[21]. To address the overdispersion of ASE, we instituted an allelic imbalance threshold of one haplotype being expressed at least 50% more than the other, corresponding to a MHF of ≥ 0.6. A similar approach and cutoff have been described in other ASE studies[21,22]. In summary, genes were determined to have ASE if: (1) sufficient gene expression (here defined as total RNA coverage ≥20, aggregated across all heterozygous loci, after TOGA), (2) binomial test with padj < 0.05, and (3) MHF ≥ 0.6.

TOGA: Transcript reassignment for Overlapping Genes in ASE

TOGA assigns reads overlapping multiple genes to their most likely gene of origin by integrating total RNA-seq expression data and gene exon/intron annotations. The detailed pipeline is described in **Fig. S7.** Specifically**,** from the set of overlapping COV genes within an individual, we first excluded non-expressed genes (TPM = 0) based on RSEM[14]. We then selected only genes that shared variants within overlapping regions and grouped the genes based on the union of overlapping variants. That is, a gene was included in a group if at least one of its variants overlapped with variants from other genes within the same group. Genes sharing no variants were considered "resolved".

We first define $V$ as a set of variants, $\{v_1, v_2, \ldots, v_M\}$, and $G$ as a set of genes, $\{g_1, g_2, \ldots, g_N\}$, where $M$ is the number of union of overlapping variants and $N$ is the number of union of overlapping genes. We assumed that an allelic count of an overlapping variant is distributed proportionally based on the genes' expression level and the variants' gene annotation status (e.g., exonic or intronic). We assumed exonic variants contribute significantly more to the count than intronic ones. Thus, raw weights ($w$) were assigned as follows: 0.99 for exonic variants (as most RNA reads originate from exons), 0.01 for intronic variants, and 0 for variant-gene pairs that do not exist.

The final weight, $W(v_i, g_j)$, representing the probability that variant $v_i$ originates from gene $g_j$, was normalized by the total weight of all genes covering that variants as formulated in Equation (1):

$$W(v_i, g_j) = \frac{w(v_i, g_j)e(g_j)}{\sum_{k=1}^{N} w(v_i, g_k)e(g_j)} \quad (1)$$

where $v_i$ is the $i^{th}$ overlapping variant, $g_j$ is the $j^{th}$ overlapping gene, $e(g_j)$ is $g_j$'s TPM, and $w(v_i, g_j)$ is the raw weight function defined above. Next, we summed the adjusted allelic counts of all overlapping variants based on $W$, followed by normalization across all overlapping genes. We defined this metric as CPOV (Contribution Proportion of Overlapping Variants) for $g_j$ in Equation (2), which represents the proportion of total overlapping reads contributed to a given gene.

$$CPOV_j = \frac{\sum_{i=1}^{M} C(v_i)W(v_i,g_j)}{\sum_{j=1}^{N}\sum_{i=1}^{M} C(v_i)W(v_i,g_j)} \quad (2)$$

where $C(v_i)$ is an original allelic count of $v_i$. For example, if there were a total of 100 RNA reads in a genomic interval where two genes, denoted X and Y, overlapped, and our TPM- and exon/intron-based weighting algorithm estimated that 80 reads came from gene X and 20 from gene Y, then the CPOV would be 0.8 and 0.2, respectively.

To further estimate the likelihood of incorrect read assignment, we defined another metric, IRPG (Incorrectly assigned Read Proportion in Gene):

$$IRPG_j = \frac{(1-CPOV_j)C(v_o)}{C(v_o)+C(v_{no})} \quad (3)$$

where $C(v_o)$ and $C(v_{no})$ are the read counts for overlapping and non-overlapping variants in a gene, respectively. Following the previous example, if gene X had 60 non-overlapping reads, then its IRPG would be (1 - 0.8) ×100/(100+60) = 20/160 = 0.125, indicating that an estimated 12.5% of reads assigned to gene X likely originated from the other overlapping genes. As detailed in **Fig S7**, we implemented a stepwise classification algorithm based on these metrics. First, genes with IRPG < 0.1 were considered unaffected by read misassignment and classified as "resolved". For genes with IRPG ≥ 0.1, we computed the adjusted haplotypic total count ($Cov_{adj}$) by redistributing reads from overlapping variants based on CPOV as shown in Equation (4):

$$Cov_{adj} = CPOV \times C(v_o) + C(v_{no}) \quad (4)$$

Subsequently, genes with $Cov_{adj}$ < 20 were excluded. Among the remaining genes, those with CPOV < 0.5 were considered to have low contribution of overlapping reads, thus we opted to use only non-overlapping reads for their ASE analysis, contingent on non-overlapping coverage ≥20, and classified as "resolved by low CPOV." For the genes with CPOV ≥ 0.5, given the higher contribution of overlapping reads, we opted to use all originally assigned reads for ASE analysis, but classified these genes as "unresolved" given the difficulty in accurately ascertaining haplotype-specific overlapping read counts. A detailed example illustrating the entire TOGA workflow is provided in **Fig. S8.** In ASE analysis, we calculated adjusted binomial p-values using the "resolved", "resolved by CPOV", and "unresolved" genes only. This method was consistently applied to the NEPTUNE WGS+RNA analysis, the NEPTUNE RNA-only analysis, and the analysis of our control samples.

Identification of imprinted genes

A list of known imprinted genes was curated using the union of two sources: Tucci et al.[23], which identified 313 human imprinted genes through an extensive literature review, and Baran et al. [18], which used statistical modeling to identify 42 tissue-specific imprinted candidate genes in 48 human adult tissues from the GTEx dataset. The union of these two datasets resulted in 316 known imprinted genes. We then corrected discrepancies in gene names between the two datasets and GENCODE v45 annotation[15] and compiled a final list of 299 known imprinted genes (**Table S6**).

To identify imprinted genes in our NEPTUNE samples, we incorporated the Baran pipeline, which assigns an imprinting likelihood to a gene based on ASE data. It uses a statistical model to label a variant in an individual as monoallelic, imbalanced, or biallelic expression based on the allelic counts. These variants in a gene are then aggregated across all individuals to compute a generalized likelihood ratio for imprinting (impglr) for the gene. It uses a unique characteristic of imprinted genes, which is that monoallelic expression pattern of a variant is agnostic to allelic identity due to the parent-of-origin mechanism of imprinting. Thus, an imprinted gene is expected to contain variants with monoallelic expression of the reference allele in some individuals and monoallelic expression of the alternative allele in others.

As input to the Baran pipeline, we used variant-level allelic counts from phASER. We only analyzed variants with RNA coverage ≥ 8 (RNA COV8) in COV genes. In addition, we applied a HWE p-value ≥ 0.001 following the Baran pipeline. This resulted in an average of 10,007 ± 1,571 variants per individual in GLOM and 8,944 ± 1,415 variants per individual in TUBE. Genes containing such variants (n = 15,963 GLOM; n = 14,886 TUBE) were then required to pass two additional criteria to be labeled as imprinted, as outlined Baran et al. First, the gene required sufficient data to consider its imprinted status. This included having at least two distinct variants and at least 5 individuals with a heterozygous variant in the gene. Additionally, the gene was required to contain a variant with monoallelic reference expression in one individual and monoallelic alternative expression in another (a unique signature of parent-of-origin mechanism). We removed heterogeneously imprinted genes using the pipeline's hetlr metric, defined as the likelihood ratio of heterogeneity of imprinting across the population, compared to other imprinting status. Genes with hetlr ≥ 1 were defined as heterogeneously imprinted and removed. The remaining genes were amenable to imprinting analysis (n = 2,419 GLOM; n = 2,619 TUBE) and were tested against the second criterion: a sufficient likelihood of imprinting. A gene was labelled as imprinted if its generalized likelihood ratio for imprinting (impglr) > 40 and ≥ 80% of its variant-individuals within the gene showed monoallelic expression.

Following this initial analysis, we found that our "novel" imprinted candidates were significantly enriched for strong eQTLs (nominal $P < 1 \times 10^{-6}$) compared to "known" imprinted genes. This suggested that a strong eQTL effect could mimic an imprinting signal. Therefore, we developed a subsequent filter using three binomial tests to assess whether an eQTL provided a better explanation for the observed ASE. For this correction, we gathered each gene's significant eQTLs ($P < 1 \times 10^{-6}$) from the NEPTUNE eQTL data[24]; if no significant eQTL was present, we used the variant with the lowest eQTL *P*. For each eQTL in a gene, we stratified the gene's coding variant-individual pairs into three groups. The "homozygous eQTL group" held all coding variant-individuals where the individual was homozygous for the eQTL. The "heterozygous alternative group" encompassed all coding variant-individuals where the individual was heterozygous for the eQTL and the alternative allele of the coding variant was phased with the alternative allele of the eQTL (purple group in **Figs. S14-17**). Finally, the "heterozygous reference group" encompassed all coding variant-individuals where the individual was heterozygous for the eQTL and the reference allele of the coding variant was phased with the alternative allele of the eQTL (red group in **Figs. S14-17**). We designed the three binomial tests such that the expected result was imprinting, while the alternative hypothesis represented an eQTL-driven ASE mechanism. Any gene initially labelled as an imprinting candidate that showed a significant p-value (supporting the eQTL hypothesis) in ≥ 2 of these three tests was relabeled as an eQTL (**Figs. S14-17**).

1. Homozygous eQTL group: For an imprinted gene, expression should remain monoallelic. For an eQTL, expression should become balanced. Therefore, a "success" (imprinting) was defined as near-monoallelic expression (≥ 90% RNA reference or alternative allele ratio). We set an expected success rate of 90%. A $p_{hom} < 0.05$ supported an eQTL mechanism.

2. Heterozygous alternative and reference groups: For an imprinted gene, expression should be 50% monoallelic reference and 50% monoallelic alternative, regardless of phasing. A binomial "success" (imprinting) was defined as an RNA reference allele ratio ≥ 50%, with an expected success rate of 50%. The alternative hypothesis (eQTL) posited that allelic imbalance direction would be consistent with the eQTL allelic direction. Note that we have two tests here as we perform the test for each allele (reference and alternative), separately.

Characterization of gene- and variant-level ASE mechanisms

To assign a molecular mechanism to each ASE event at the gene-individual level, we first filtered for resolved or unresolved (as determined by TOGA) ASE gene-individual pairs with RNA read coverage ≥ 20 (COV genes). Each ASE gene-individual event was then assigned as follows. First, if a gene was identified as "imprinted" by the pipeline described above, that label took precedence over other mechanisms. If not imprinted, gene-individual pairs were labelled as "eQTL" if the individual was heterozygous for a variant with an eQTL nominal p-value $< 1 \times 10^{-6}$ associated with that gene within 1Mb of the gene, using NEPTUNE eQTL results from our prior study[24]. For sQTLs, we used GTEx v10 sQTL-sGene pairs from kidney cortex[25], as determined by LeafCutter[26]. If the individual was heterozygous for a significant sQTL (i.e., any variant in the sQTLs.signif_pairs.tsv file from GTEx), the associated gene-individual pair was labelled as "sQTL". ASE gene-individual pairs containing both a significant sQTL and eQTL were labelled "sQTL+eQTL".

We also evaluated ASE at the variant level to assess allelic preference across individuals. We focused on variants with RNA coverage ≥ 8 within COV genes. Similar to gene-level ASE, an "ASE variant" was defined as a variant where one allele's expression was 50% greater than the other, with an adjusted binomial p-value < 0.05 (expected=0.5). On average, 3.83 ± 2.85% of variants were ASE variants per individual in GLOM, and 1.76 ± 0.94% in TUBE.

At the variant level, we calculated a variant's allelic preference across the population as the proportion of individuals with ASE of the alternative allele, relative to the total number of individuals with ASE for that variant. We filtered for variants where >10% of individuals were heterozygous and ≥ 50% of those heterozygous individuals had ASE. This was applied because variants with ASE in only a few individuals are limited to discrete, non-granular preference values (e.g., 0%, 50%, or 100% for two individuals). Subsequently, we assessed the overlap between these ASE variants and both eQTLs and sQTLs to determine if these mechanisms were enriched among variants with high directional consistency. To label eQTLs and sQTLs, we applied the same criteria as gene-level ASE classifications as above, except that the ASE variant should meet those criteria. Among the 55,423 variants with ASE in ≥ 1 individuals in NEPTUNE GLOM, 2,485 (4.48%) were eQTLs and 2,786 (5.03%) were sQTLs.

Prediction of regulatory variants in promoters

To explore the effect of promoter regulatory variants on ASE, we compared the prevalence of such variants in the promoters of ASE genes against a baseline established from balanced (non-ASE) genes. We conducted the following analysis on all COV genes in all individuals: First identified the open chromatin region containing the gene's transcription start site (TSS) and gathered all heterozygous variants within that region. We then used the ChromKid model developed by Loeb et al. to score each heterozygous variant's likelihood of impacting chromatin accessibility[27]. To score variants in GLOM, we computed the average of the variant's absolute individual score in immune cells, T cells, stromal cells, podocyte cells, and endothelial cells. For TUBE, we computed the average of the variant's absolute score in immune cells, collecting duct, T cells, endothelial cells, distal tubule, proximal tubule, and loop of Henle. Variants that scored in the top 20% (≥ 28.08 in GLOM; ≥ 21.72 in TUBE) were labelled as predicted regulatory variants (PRVs).

We subsequently split the COV gene-individual pairs into three groups: balanced expression, ASE with a known mechanism, and ASE with an unknown mechanism. We computed the proportion of balanced gene-individual pairs with a predicted regulatory variant in their promoters, thus generating a baseline likelihood of identifying PRVs in a typical gene's promoter. We stratified these groups by the rarity (minor allele count, MAC) of the PRV (e.g., proportion where the PRV has an MAC=1, MAC≤5, etc.).

For known/unknown ASE, we first grouped the gene-individual pairs by the number of individuals with ASE for that gene, generating subsets of increasing ASE gene commonality (e.g., ASE observed in 1, ≤5, ≤10, …, ≤150 individuals). As with the balanced group, in each ASE commonality subset, we calculated the proportion of gene-individual pairs that had a promoter PRV, stratifying these groups by the PRV rarity. In summary, ASE gene-individual pairs were grouped by: ASE mechanism (known/unknown), ASE gene commonality, and MAC of the promoter PRV.

Lastly, we calculated the enrichment of predicted regulatory variants of a given MAC in each ASE group by normalizing the proportions to the corresponding balanced gene group. For instance, consider a subset defined by private ASE events (number of individuals with ASE = 1) of an unknown ASE mechanism. If 8% of these gene-individual pairs have a singleton PRV (MAC=1), this value was compared to the baseline proportion of balanced gene-individual pairs carrying a singleton PRV in their promotors (e.g., 4%). The resulting PRV enrichment in this group would be 2, indicating a two-fold enrichment of singleton PRVs in private ASE genes of unknown mechanism.

RMSE and haplotypic flip analysis

To measure overall discordance across heterozygous allele counts on the same haplotype, we calculated the root-mean-square error (RMSE) of the haplotype A fraction for each ASE gene. This was computed using the gene-level haplotype A fraction from phASER Gene AE as the expected value, summed over the variant-level haplotype A fractions for all variants within that gene as follows:

$$RMSE_{gene} = \sqrt{\sum_{i}^{n_{var}} \frac{(\mu_{var,i} - \mu_{gene})^2}{n_{var}}} \quad (5)$$

where $n_{var}$ is the number of variants in the gene, $\mu_{var,i}$ is haplotype A fraction of the $i^{th}$ variant, and $\mu_{gene}$ is haplotype A fraction of the gene. We included only variants with RNA coverage ≥ 8 (RNA COV8) in COV genes with ≥ 2 such variants. ASE genes were labelled by their mechanism as described previously.

Our hypothesis was that sGenes would exhibit higher RMSE values due to variable inclusion/exclusion of variants within a gene. To accurately measure this, no gene-level haplotypic flips should occur within the gene. A haplotypic flip is a technical error where variants within a gene are incorrectly phased, resulting in the assignment of variant allelic counts to the wrong haplotype. In phASER, this can occur when two adjacent haplotype blocks for a gene are not connected by an RNA read, or "edge." When this occurs, phASER falls back on WGS phasing to connect the blocks. While this WGS-based connection is often correct, it is less reliable and prone to error, particularly for rare variants. Therefore, for the RMSE analysis, we only used "connected" gene-individual pairs, defined as those where all haplotype blocks possessed an edge to adjacent blocks, making no gene-level haplotypic flips possible. While connected genes represented only 8.53% of all gene-individual pairs, this filtering was critical for an accurate RMSE analysis.

The importance of this filter was confirmed using imprinted genes. Before filtering, some imprinted gene-individual pairs displayed spuriously high RMSE values (**Fig. 3B, S20E**). Further investigation revealed these were caused by gene-level haplotypic flips. We determined the prevalence of these flips by identifying imprinted genes where two variants within the same individual showed monoallelic expression (≥ 90% allelic ratio) towards opposite haplotypes. Filtering for "connected" genes successfully removed these high-RMSE imprinted genes, resulting in the expected low RMSE values for this category (**Fig. 3B, S20E**). We note that the Baran pipeline[18] for labeling imprinted genes was not impacted by these flips, as it directly uses variant-level allelic counts. Lastly, our hypothesis of higher RMSE values in sGenes was specific to sGenes with heterozygous variants located both within and outside the alternatively spliced intron; we termed these "Mixed sGenes." Many sGenes had heterozygous variants located only within the alternatively spliced region or only outside of it, leading all measured variants to have a similar effect. We termed these "Uniform sGenes." As expected, the "Mixed sGene" category displayed substantially higher RMSE values than any other ASE mechanism, including

"Uniform sGenes". Therefore, the "sGene" category presented in **Fig. 3B** and **Fig. S20E** consists exclusively of these "Mixed sGenes."

Calculating cell-type proportion of ASE genes using single-cell RNA-seq

To calculate the cell-type proportion of ASE genes, we used single-cell RNA sequencing (scRNA-seq) data generated by the Kidney Precision Medicine Project (KPMP) consortium[28]. This dataset, generated using the 10x Genomics platform, consisted of 79 samples (14 AKI, 37 CKD, and 28 healthy), comprising 31,496 genes and 225,177 cells in total. We systematically classified the 54 distinct cell types from KPMP into GLOM and TUBE compartments and subsequently regrouped them for analysis (**Table S7**). The original cell types were consolidated into 12 major categories: 2 unique to GLOM, 6 unique to TUBE, and 4 shared between compartments. Unique cell types within the GLOM compartment included podocytes (POD) and parietal epithelial cells (PEC). The TUBE compartment uniquely comprised proximal tubule cells (PT), intercalated cells (IC), distal convoluted tubule cells (DT), thick ascending limb cells (TAL), descending/ascending thin limb cells (DTL/ATL), and principal cells (PC). Cell types present in both compartments included fibroblasts (FIB), immune cells (IMMUNE), endothelial cells (EC), and vascular smooth muscle cells/mesangial cells (vSMC/MC).

For each compartment, normalized expression of each gene was quantified using a pseudobulk approach. Specifically, raw counts from individual cells were summed within each grouped cell type per gene. These counts were then normalized by the total expression of that cell type and scaled by a factor of 10,000. The cell-type proportion for a given gene was subsequently calculated as the proportion of its normalized expression within a specific cell type, divided by the total normalized expression for that gene across all grouped cell types in the corresponding compartment.

To define cell-type predominance of ASE genes, we considered an ASE gene to be enriched in a particular cell type if its cell-type fraction exceeded twice the expected value under uniform expression. For example, in GLOM, with 6 grouped cell types, a gene was considered cell-type enriched if its fraction was $> (1/6) \times 2 = 0.33$. For TUBE, with 10 grouped cell types, the threshold was $> (1/10) \times 2 = 0.2$.

The cohort of "control" kidneys

The control kidney cohort comprised 31 samples: 23 samples of non-affected kidney tissue from tumor nephrectomies in the PRECISE cohort[29] and 8 living kidney donor biopsies from the University of Michigan (**Table S1**). All control samples were microdissected into GLOM and TUBE following the same protocol as NEPTUNE; only paired-end bulk RNA-seq data were available for these samples. Participants in the PRECISE cohort had clinically preserved kidney function (estimated glomerular filtration rate [eGFR] ≥ 60 ml/min) without proteinuria (urine dipstick with negative or trace protein – within normal range). PRECISE samples were surgical sections instead of needle biopsy. Living donors met standard screening criteria for kidney donation, including age >18 years, preserved GFR, and absence of hypertension, diabetes, or proteinuria (specific demographic and clinical details were not available for this subgroup). Ancestry and sex were predicted by Peddy[30] based on RNA-seq derived genotypes. These study cohorts have been approved by UMich IRB (PRECISE: HUM00165536, HUM00052918; Living donors: HUM0002468).

ASE analysis pipeline using only RNA-seq data

To analyze our control cohort where only RNA-seq data are available, we developed a parallel optimized ASE analysis pipeline, which we termed "RNA-only" ASE pipeline (**Fig. S22A**). This workflow starts with RNA BAM files aligned by STAR (v2.7.11b) and duplicate-marked by samtools (v1.19.2). With the BAM file as input, genotype calling was then performed using the GATK variant calling workflow (v4.5.0.0; https://gatk.broadinstitute.org/hc/en-us/articles/360035531192-RNAseq-short-variant-discovery-SNPs-Indels). We applied SplitNCigarReads to segment paired-end reads with Ns in their CIGAR string. BaseRecalibrator was then used to correct base quality scores with inputs of known polymorphic sites (SNPs and indels) from the GATK Resource Bundle and reference genome from GENCODE v45 primary assembly[15]. After applying quality recalibration with ApplyBQSR, variants were called using HaplotypeCaller using a minimum phred-scaled confidence threshold of 20. Finally, hard filtering was performed with VariantFiltration, retaining variants with Fisher Strand (FS) $\leq 30$ and Quality by Depth (QD) $\geq 2$. Only autosomal heterozygous variants were retained in the final VCF files.

An initial ASE analysis using this RNA-derived VCF and WASP-filtered RNA BAMs revealed a median of six monoallelic genes per individual, a strong indicative of false-positive heterozygous calls, as the variants supporting these monoallelic genes should have been called homozygous. We identified two primary causes: (1) the use of non-WASP-filtered BAMs in the variant calling step, which allowed reference-biased reads to be miscalled as heterozygous, and (2) low-quality reads contributing to erroneous calls. To address this, we refined the pipeline by re-running the entire GATK variant calling workflow using the WASP-filtered BAMs and removed all variants that exhibited monoallelic expression. To further increase confidence, we excluded variants not present in the 1000 Genomes (1KG) project[31] and variants listed in the standard phASER blacklist. The subsequent steps for ASE calling were identical to those used in the WGS+RNA pipeline, including running phASER, applying TOGA, and calling ASE genes.

Survival analysis

To assess the association between disease progression and an elevated GLOM ASE proportion, we performed survival analysis. For the primary analysis, the event variable was defined as the occurrence of end-stage kidney disease (ESKD) or a $\geq 40\%$ decline in eGFR from the time of biopsy (eGFR40). The time variable was the number of days from biopsy to the ESKD or eGFR40 event. Right-censoring was applied to individuals without events, and samples without eGFR information were excluded, resulting in a total of 194 samples for this analysis. We defined kidney failure as two consecutive eGFR < 15 ml/min or ESKD requiring kidney replacement therapy.

We also analyzed time to complete remission among patients with active disease at biopsy. This analysis was restricted to patients with a urine protein-to-creatinine ratio (UPCR) > 0.3 g/g at biopsy, indicating active disease. Time was calculated from the biopsy date to either complete remission or the last follow-up. The primary endpoint was the achievement of complete remission, defined as UPCR $\leq 0.3$ g/g. Patients were censored at their last follow-up if remission had not been achieved. Using the same groups as our primary analysis, we excluded samples without remission data, resulting in a final cohort of 172 samples (84 in the high-GTAR group and 88 in the low-GTAR group).

Lastly, we assessed time to first relapse among patients who were in remission at the time of biopsy. Time was calculated from the biopsy date to either the first relapse or the last follow-up. The endpoint was disease relapse, defined as UPCR > 3.5 g/g. Patients were censored at their last follow-up if they remained relapse-free. This analysis included 21 patients (12 in the high-GTAR group and 9 in the low-GTAR group).

Differential gene expression and gene set enrichment analysis

We performed differential gene expression (DGE) analysis to identify genes differentially expressed between the high- and low-GTAR groups, as defined in the survival analysis. 3 low GTAR samples without complete information for sex, age, race, or RNA-seq batch were excluded, resulting in 97 samples in the high-GTAR group and 94 samples in the low-GTAR group. DGE analysis was performed using DESeq2[32] with RNA counts quantified by RSEM. The model adjusted for sex, age, race, and RNA-seq batch as covariates. Only autosomal genes with at least 10 counts in a minimum of three samples were retained for the analysis to exclude low-count, high-variability genes.

Gene set enrichment analysis (GSEA) was subsequently performed using clusterProfiler[33], with genes ranked by the product of signed($\log_2$FC) × $-\log_{10}$(P) from the DGE analysis. Genes with null adjusted p-values from DESeq2 were excluded. The gene groups evaluated were from the Gene Ontology (GO) databases, including Biological Process (BP), Molecular Function (MF), and Cellular Component (CC) terms, and pathway databases (KEGG and WikiPathways). P-values were adjusted using the Benjamini-Hochberg correction within each database, with significant terms defined as those with an adjusted $P < 0.05$. As a complementary approach, over-representation analysis (ORA) was also performed on the sets of significant upregulated ($\log_2FC > 0.5$, $P_{adj} < 0.05$) and downregulated ($\log_2FC < -0.5$, $P_{adj} < 0.05$) genes identified by DESeq2.

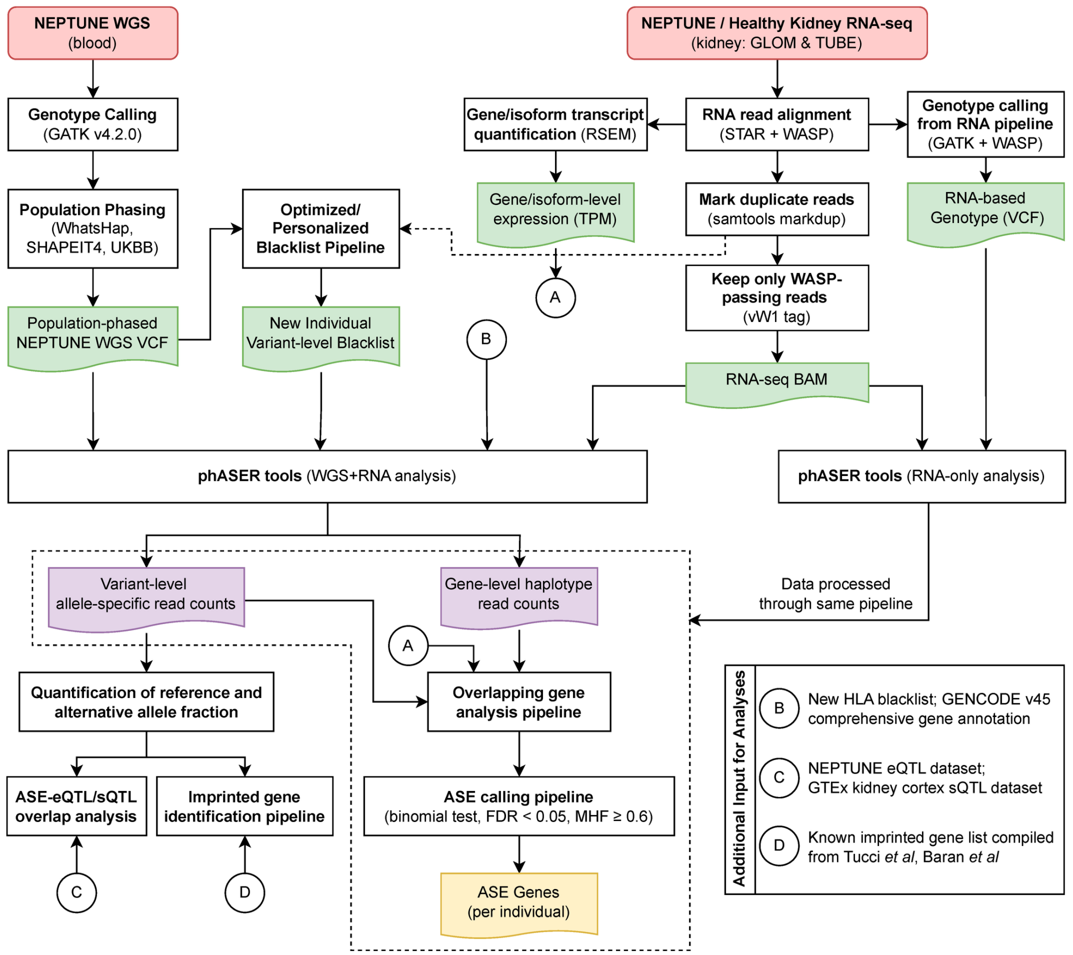


**Fig. S1. Flowchart of complete ASE analysis pipeline.** Red boxes indicated input datasets; white rectangles indicate bioinformatical steps; green document symbols indicate generated datasets/files pre-phASER; purple document symbols indicated generated datasets/files post-phASER; yellow document symbol indicates generated dataset/file after ASE calling. Additional input for specific analyses/steps are indicated by letters in circles, referenced in the legend.

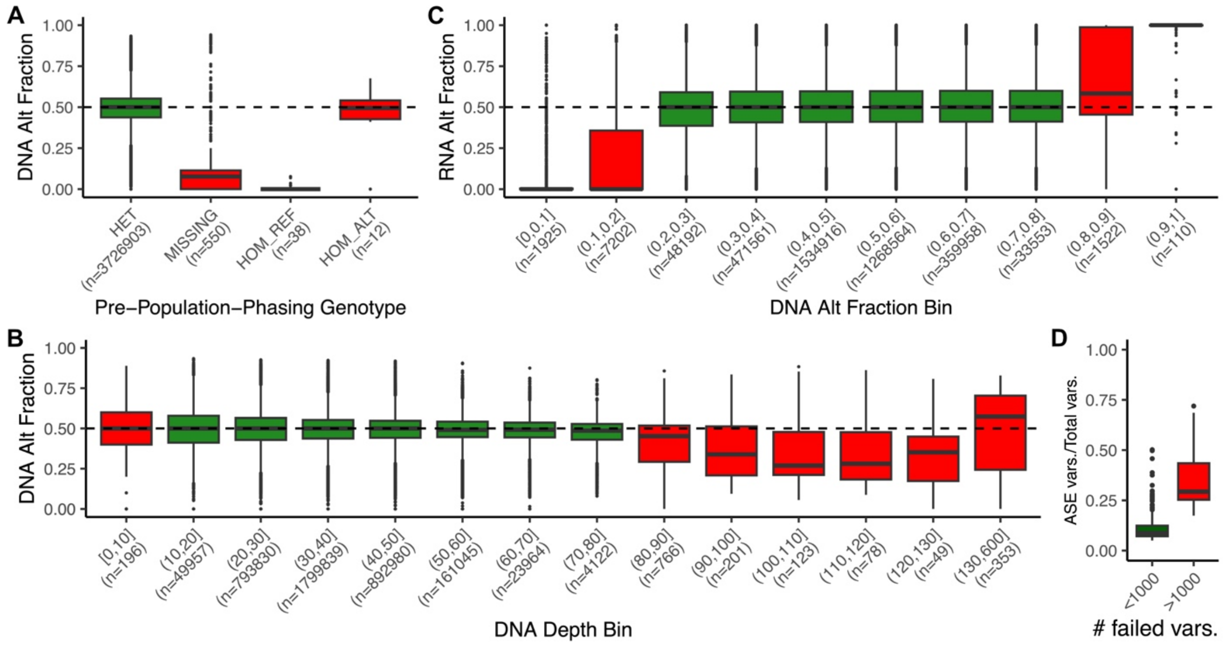


**Fig. S2. DNA/RNA alternative (alt) fraction stratified by key metrics in the WGS data. (A)** Distributions of DNA alt fractions of post-phasing heterozygous variants stratified by their pre-phasing genotypes. Variants called as heterozygous prior to phasing are shown in green, while variants with mismatched genotypes between pre- and post-phasing are shown in red. True heterozygous variants are expected to have a DNA alt fraction near 0.5 (dashed line). **(B)** Distributions of DNA alt fraction in post-phasing heterozygous variants stratified by their DNA sequencing depth. Variants with extreme sequencing depths (depth ≤ 10 or depth ≥ 80) were filtered out, and colored red in the figure. **(C)** Distributions of RNA alt fraction in post-phasing heterozygous variants stratified by their DNA alt fraction bins. DNA alt fraction correlated with RNA alt fraction. Post-phasing heterozygous variants with DNA alt fraction ≥ 0.8 or DNA alt fraction ≤ 0.2 (red) showed significant deviation from the expected RNA alt fraction of 0.5 (dashed line), and were therefore filtered out. **(D)** Proportion of total variants with ASE in a sample, stratified by the number of variants that failed the WGS filtering parameters. Samples with >1000 failed variants were filtered out.

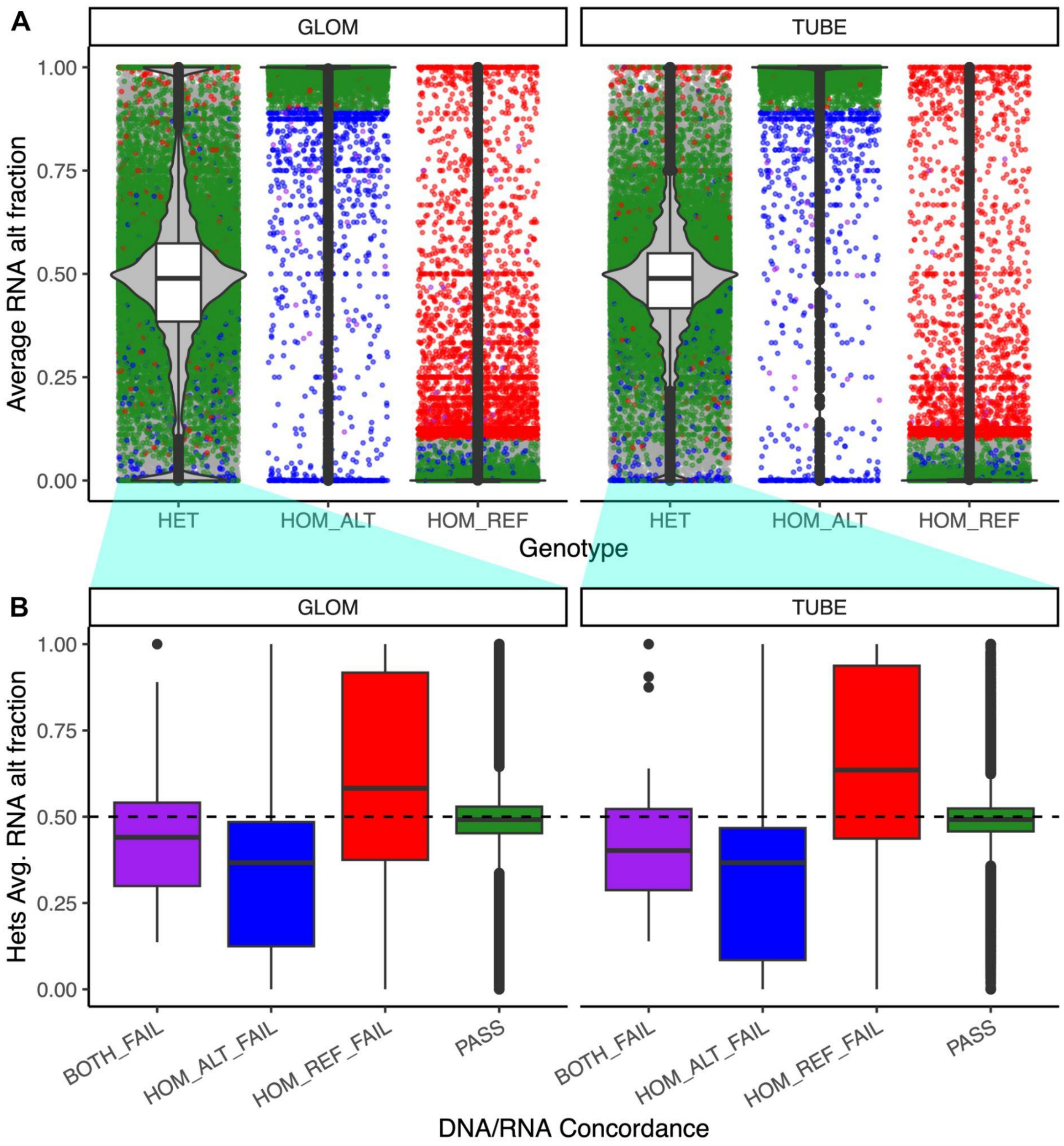


**Fig. S3. Concordance of VCF genotype with allelic expression in RNA-seq. (A)** Average RNA alt fraction in variants stratified by genotype groups. Variants whose HOM_ALT genotype was discordant with their expression (avg. RNA alt < 0.9) were colored blue and labeled HOM_ALT_FAIL in (B). HOM_REF_FAIL (red) represents any variant whose HOM_REF group had average RNA alt fraction > 0.1. Variants that failed in both the HOM_REF and HOM_ALT group were colored purple. Grey dots represent variants that pass in one homozygous group while there were no individuals in the other homozygous group. **(B)** Expansion of the HET variants in GLOM and TUBE to observe the Avg. RNA alt fraction distribution of het variants whose HOM_ALT/HOM_REF groups failed. Data is not WASP-corrected, and there is a slight bias for the reference allele.

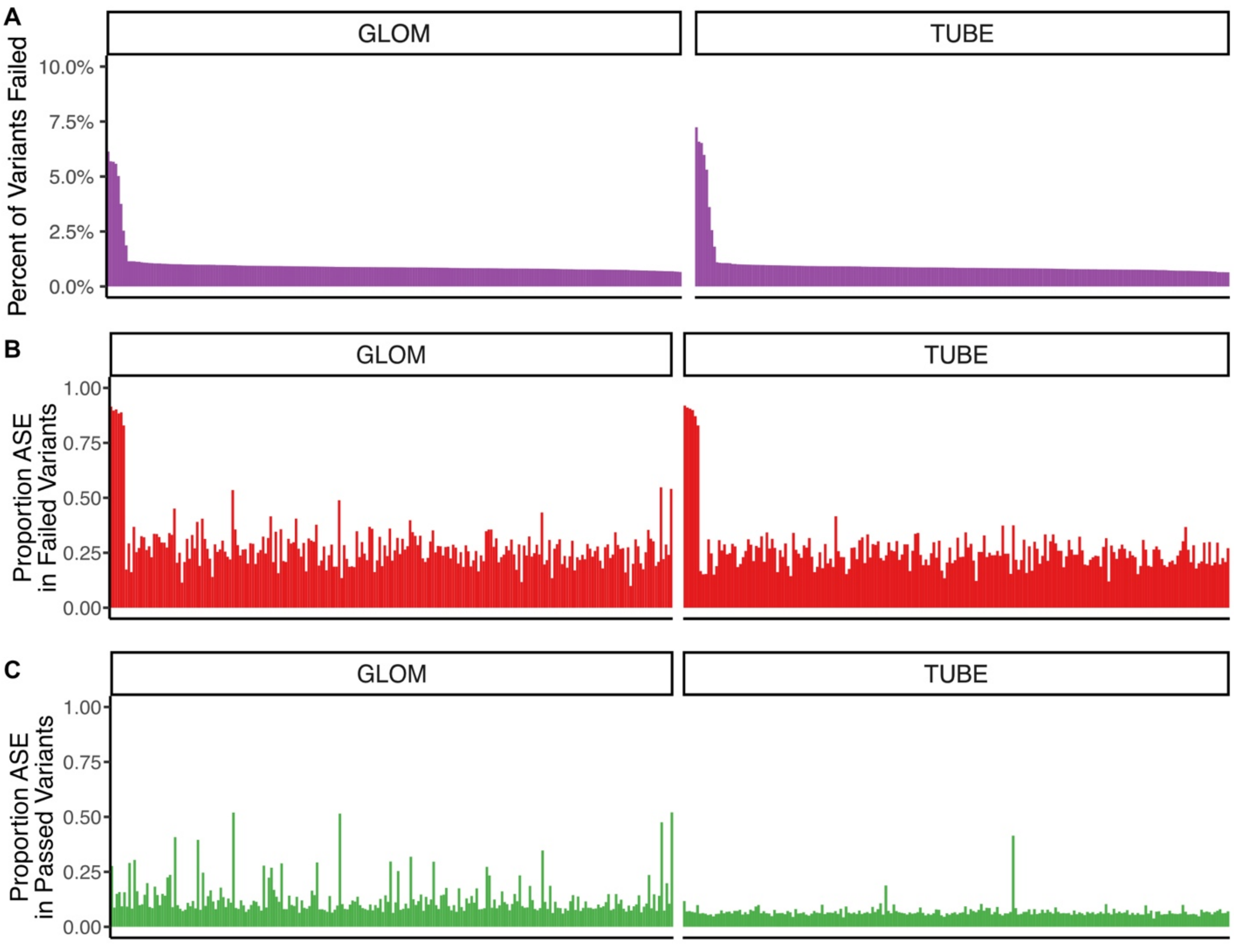


**Fig. S4. Assessment of variant blacklist filtering and impact on ASE. (A)** Percent of total variants that were excluded by the blacklist. Each bar is a single individual (n = 222 in GLOM, n = 206 in TUBE) and individuals are ordered high-to-low by percent of total variants blacklisted. This ordering is maintained in (B) and (C). **(B)** Proportion of variants that exhibited ASE among the variants that failed our blacklist. Each bar is an individual, and individuals are ordered as in (A). **(C)** Proportion of variants that exhibited ASE among the variants that passed our blacklist. Each bar is an individual, and individuals are ordered as in (A).

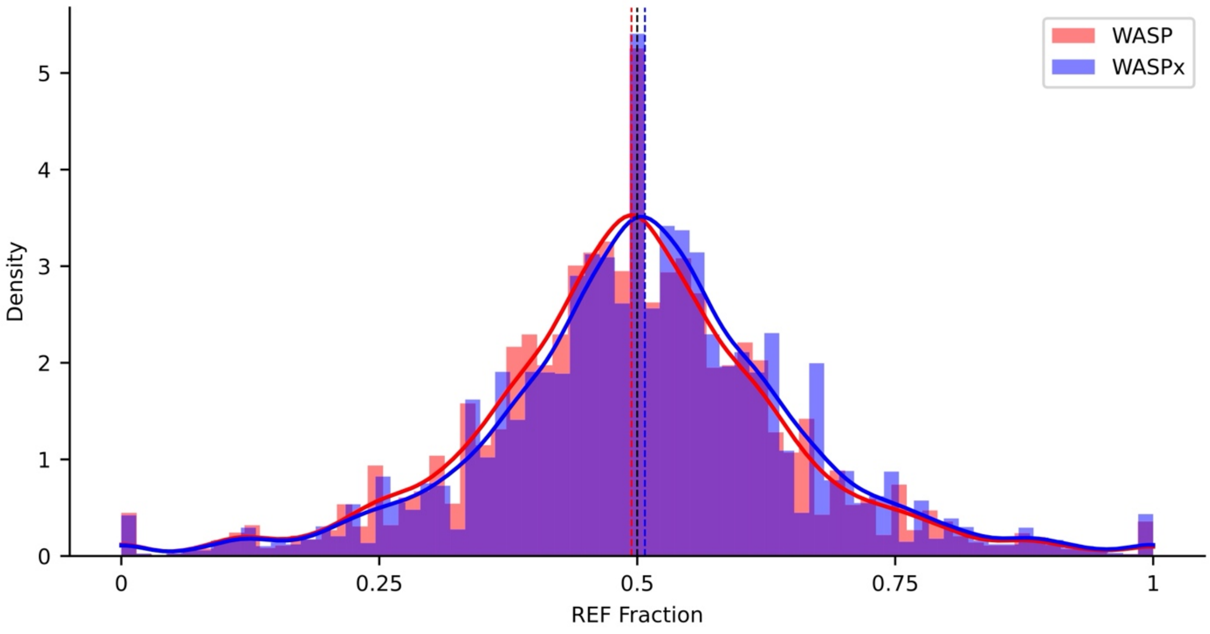


**Fig. S5. Effect of WASP filtering on fraction of reference genotype.** The histogram shows the distribution of reference allele fractions for variants within genes of coverage ≥20, with allelic coverage ≥8, from a single individual. Results are shown with and without usage of WASP filtering, denoted as WASP and WASPx, respectively. The blue dashed line represents the mean reference allele fraction without WASP filtering (0.507), while the red dashed line indicates the mean reference allele fraction with WASP filtering (0.494). The black dashed line represents the expected reference allele fraction under no allelic bias (0.5).

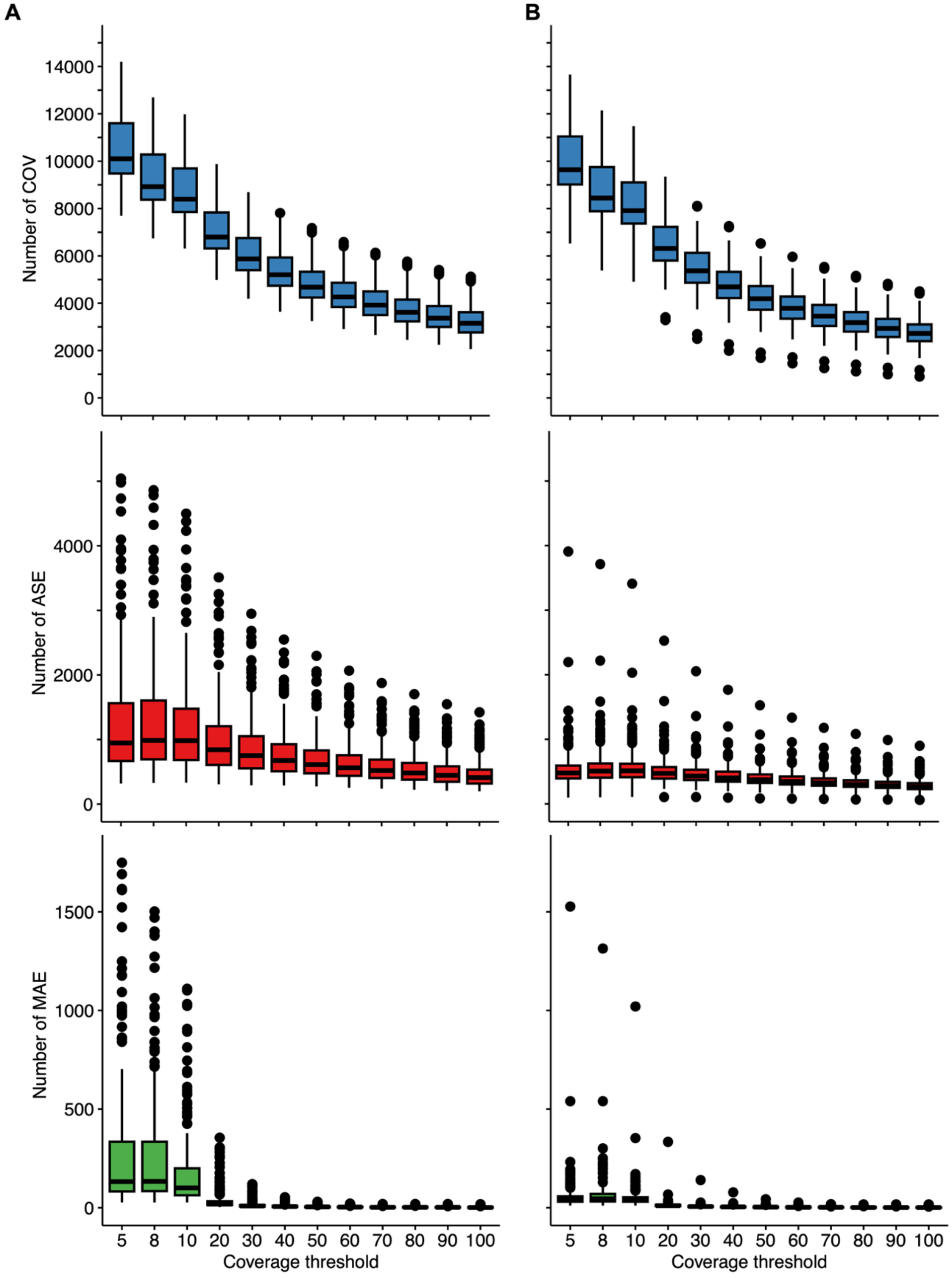


**Fig. S6. Number of COV, ASE, and MAE genes across different haplotypic count coverage thresholds. (A)** Boxplots show the number of genes per individual, with COV in blue, ASE in red, and MAE in green, for NEPTUNE GLOM. The x-axis is the minimum coverage threshold of total haplotypic count per gene from phASER and the y-axis is the number of genes from one individual. **(B)** The number of genes per individual for NEPTUNE TUBE.

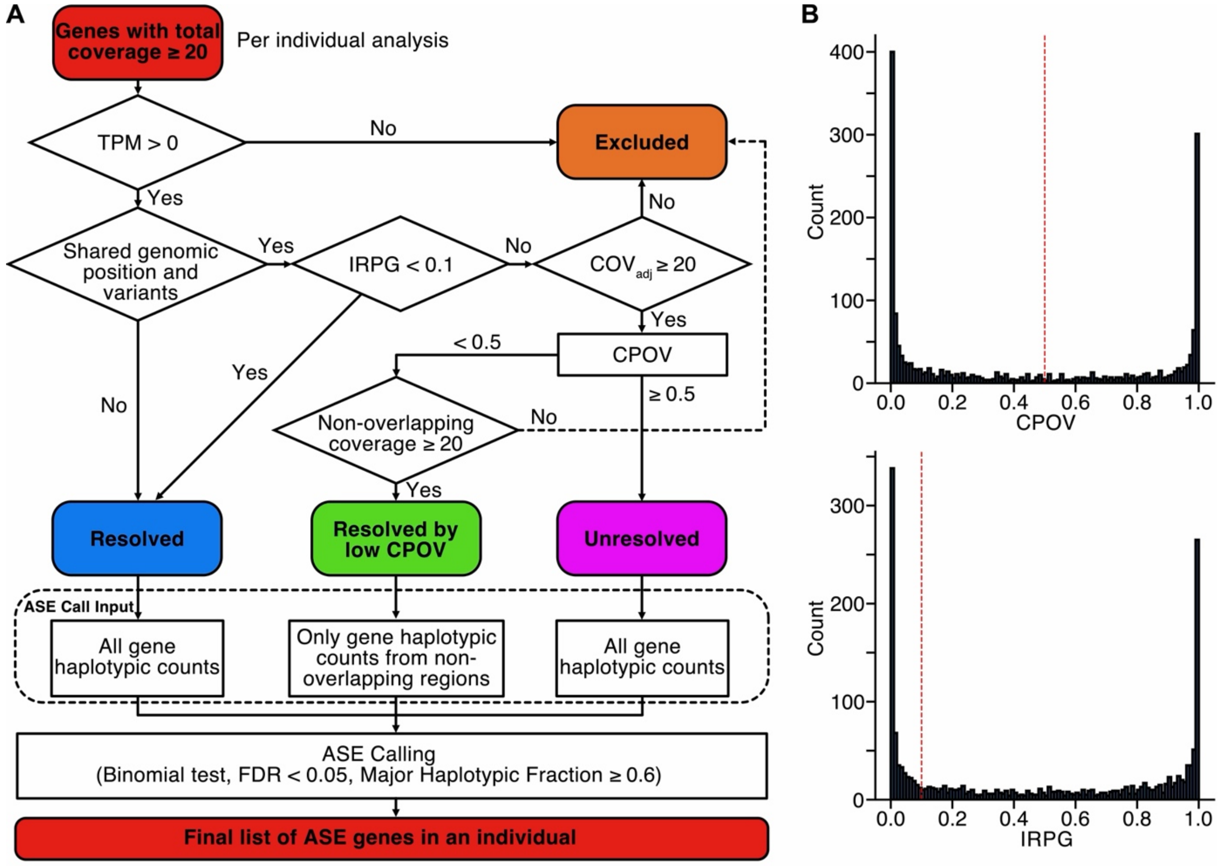


**Fig. S7. Overview of TOGA pipeline. (A)** Schematic workflow of TOGA. The input is COV genes. Each COV gene is classified as Resolved, Resolved by CPOV, Unresolved, or Excluded. COV that are not categorized as Excluded are used as inputs for ASE calling, outputting the final list of ASE genes per individual. **(B)** Distribution of CPOV and IRPG for overlapping genes in one sample. Red dashed lines indicate the threshold applied to resolve overlapping genes. CPOV=Contribution Proportion of Overlapping Variants; IRPG=Incorrectly assigned Read Proportion in Gene.

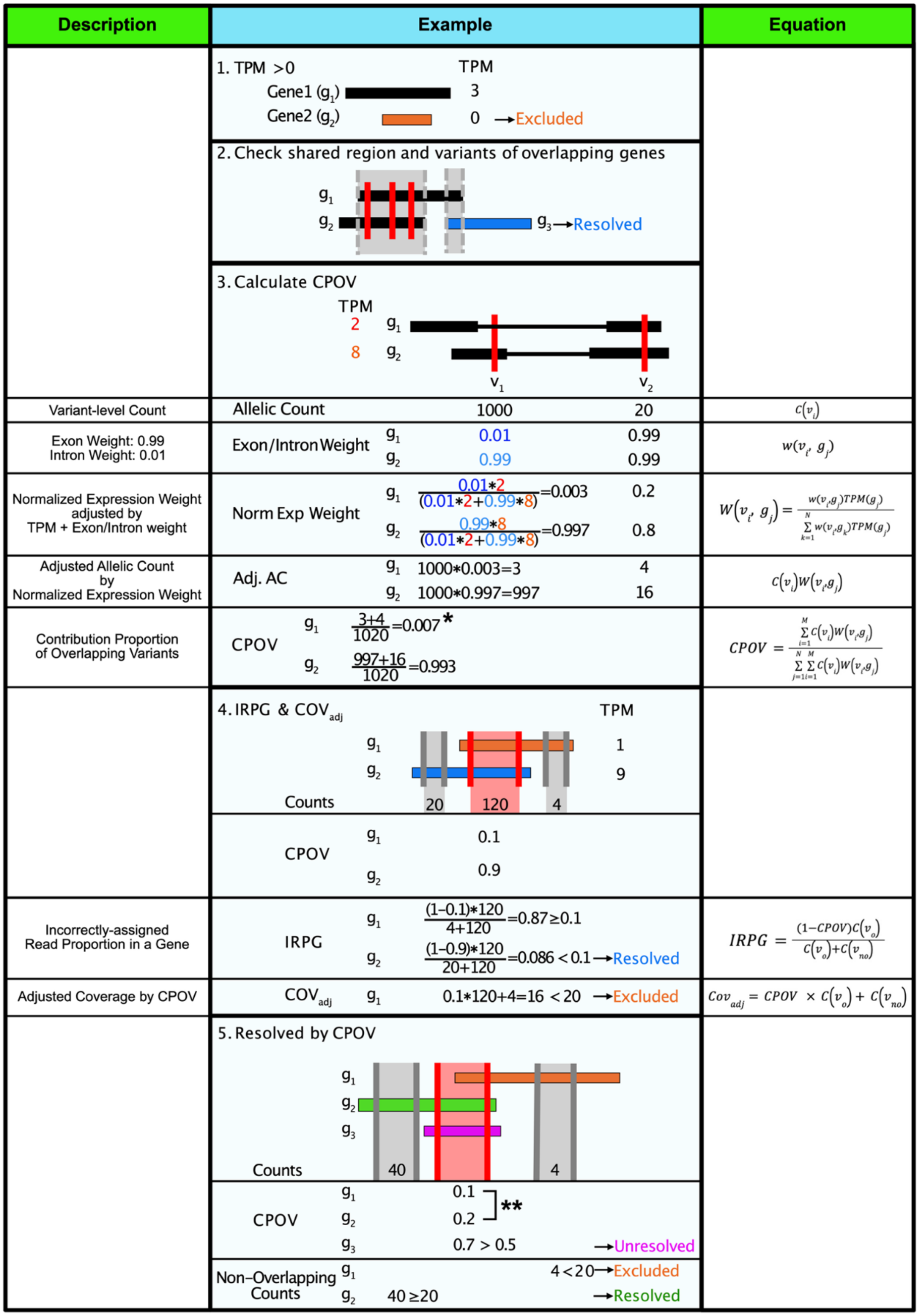


**Fig. S8. Schematic example of TOGA.** Schematic illustration of the TOGA workflow. The left panel describes key terms used in TOGA, while the right panel presents the corresponding equations for these terms. The diagram follows the sequential workflow in **Fig. S7 (A)** from top to bottom. * 0.7% of reads aligned to overlapping variants are derived from g1. ** Given the low CPOV, reads from non-overlapping variants (coverage ≥ 20) are sufficient to determine ASE.

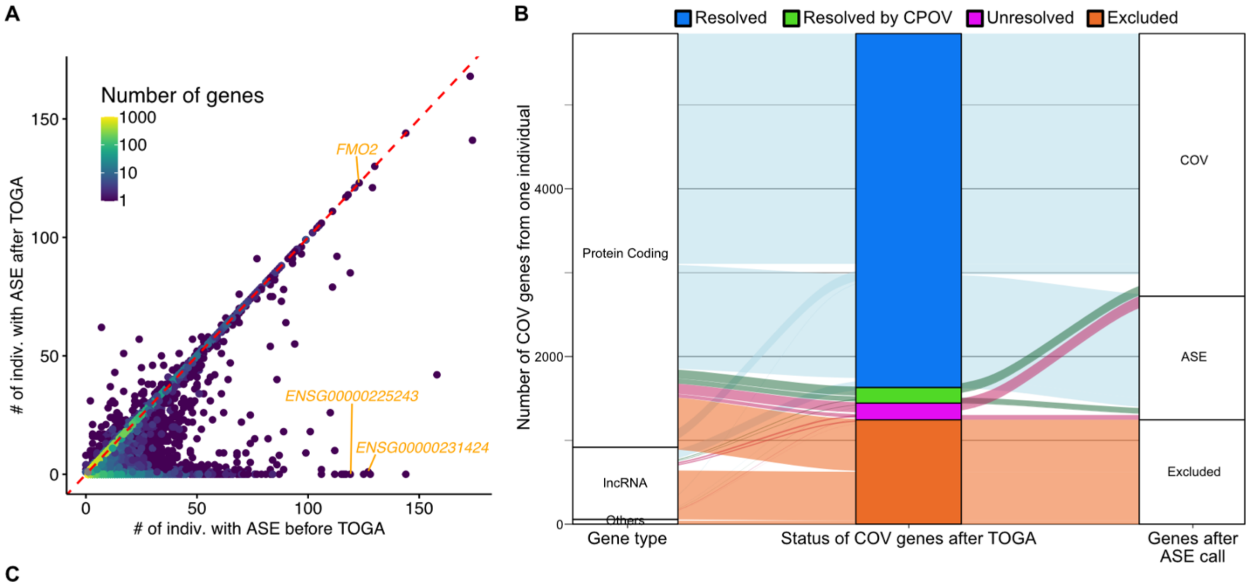


**C**

Before TOGA:

| Rank | Gene | Biotype | # indiv. with ASE (% indiv. with ASE) | # indiv. with COV |
|---|---|---|---|---|
| 1 | *KCNQ1OT1* | lncRNA | 174 (97.2%) | 179 |
| 2 | *SNHG14* | lncRNA | 173 (88.7%) | 195 |
| 3 | *KCNQ1* | protein-coding | 158 (73.5%) | 215 |
| 4 | *ZNF331* | protein-coding | 144 (97.3%) | 148 |
| 5 | *ENSG00000214265* | protein-coding | 144 (98.6%) | 146 |
| 6 | *PLCG2* | protein-coding | 130 (63.4%) | 205 |
| 7 | *SNRPN* | protein-coding | 129 (100%) | 129 |
| 8 | *ENSG00000289733* | lncRNA | 128 (63.7%) | 201 |
| 9 | *ENSG00000231424* | lncRNA | 127 (75.1%) | 169 |
| 10 | *SNURF* | protein-coding | 125 (100%) | 125 |
| 11 | *FMO2* | protein-coding | 123 (78.3%) | 157 |
| 12 | *XYLT2* | protein-coding | 121 (67.6%) | 179 |
| 13 | *LYZ* | protein-coding | 121 (78.1%) | 155 |
| 14 | *RAET1E* | protein-coding | 119 (67.6%) | 176 |
| 15 | *ENSG00000225243* | lncRNA | 119 (79.9%) | 149 |

Overlapping

TOGA

After TOGA:

| Rank | Gene | Biotype | # indiv. with ASE (% indiv. with ASE) | # indiv. with COV |
|---|---|---|---|---|
| 1 | *SNHG14* | lncRNA | 168 (93.9%) | 179 |
| 2 | *ZNF331* | protein-coding | 144 (97.3%) | 148 |
| 3 | *KCNQ1OT1* | lncRNA | 141 (97.9%) | 144 |
| 4 | *PLCG2* | protein-coding | 130 (63.4%) | 205 |
| 5 | *FMO2* | protein-coding | 123 (78.3%) | 157 |
| 6 | *XYLT2* | protein-coding | 121 (67.6%) | 179 |
| 7 | *LYZ* | protein-coding | 121 (79.1%) | 153 |
| 8 | *SNRPN* | protein-coding | 121 (100%) | 121 |
| 42 | *RAET1E* | protein-coding | 85 (71.4%) | 119 |
| 436 | *KCNQ1* | protein-coding | 42 (45.7%) | 92 |
| - | *SNURF* | protein-coding | 0 (0%) | 0 |
| - | *ENSG00000214265* | protein-coding | 0 (0%) | 0 |
| - | *ENSG00000289733* | lncRNA | 0 (0%) | 0 |
| - | *ENSG00000231424* | lncRNA | 0 (0%) | 0 |
| - | *ENSG00000225243* | lncRNA | 0 (0%) | 0 |

**Fig. S9. Effect of TOGA on ASE detection. (A)** Number of individuals with ASE for each gene in NEPTUNE GLOM (n=222) before (X-axis) and after (Y-axis) TOGA. Color heatmap represents point density. As an example, three overlapping genes (FMO2 and lncRNAs ENSG00000231424 and ENSG00000225243, shown in yellow) were initially identified as top ASE genes. However, after TOGA analysis, only FMO2 exhibited true ASE. **(B)** Alluvial plot illustrating the change in ASE gene classification after TOGA. The left panel shows the number of COV genes per individual and their gene types. The middle panel displays the TOGA-classification status. The right panel shows the ASE classification after ASE calling. **(C)** Top 15 most frequently observed GLOM ASE genes before (left) and after (right) TOGA. Matching colors with connected dots indicate genes sharing the same genomic region. Genes within the bold box retained their ASE status, while those outside the box were identified as false-positive ASE genes.

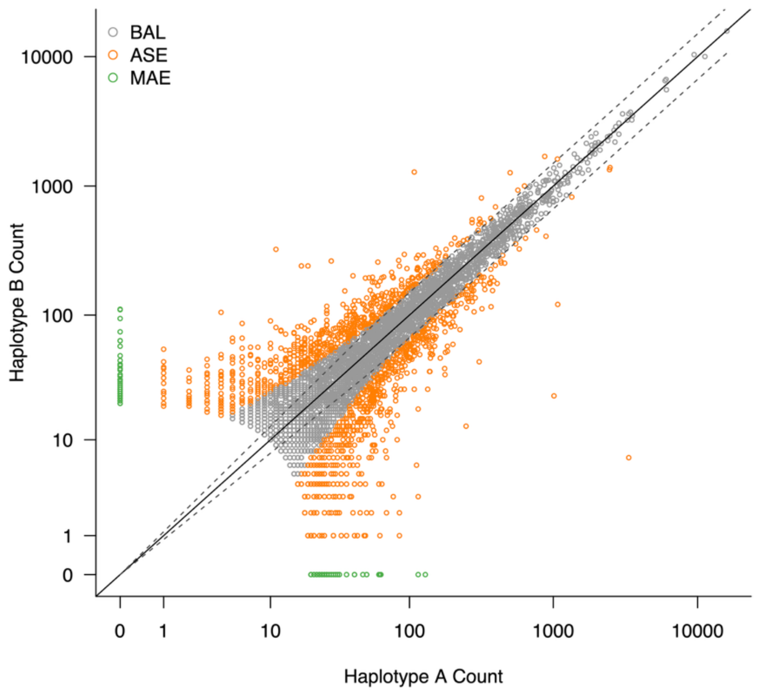


**Fig. S10. Haplotypic counts of genes from a single individual, with allelic imbalance determined by a binomial test and MHF.** Each dot indicates a COV gene from a single individual with the x-axis of counts from one haplotype and the y-axis of counts from the other, plotted on logarithmic scales. Colors indicate following gene categories: BAL, genes with no significant allelic imbalance; ASE, genes with allelic imbalance defined by a binomial test $P_{adj} < 0.05$ and MHF $\geq 0.6$, where one haplotype has $\geq$1.5 fold higher expression than the other; and MAE, ASE genes showing monoallelic expression with zero counts for one haplotype. The dotted grey line indicates the MHF $\geq 0.6$ threshold.

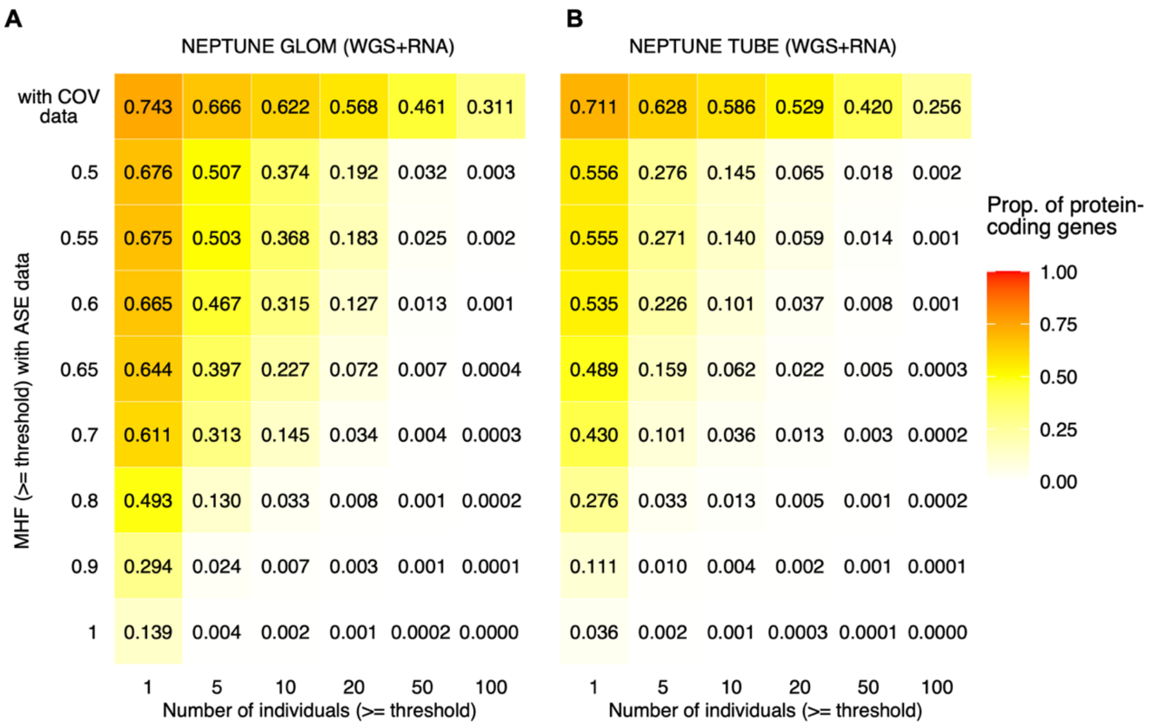


**Fig. S11. Proportion of protein-coding ASE genes across different thresholds of major haplotypic fraction (MHF) and minimum number of individuals. (A)** Heatmap showing the proportion of protein-coding ASE genes in NEPTUNE GLOM across varying MHF thresholds (y-axis) and minimum numbers of individuals in which a gene is observed (x-axis). The denominator is fixed to all autosomal protein-coding genes annotated in GENCODE v45 (n = 19,114). Heatmap color reflects the proportion of protein-coding genes meeting each criterion, and the top row represents the proportion of protein-coding COV. **(B)** The same analysis performed for NEPTUNE TUBE.

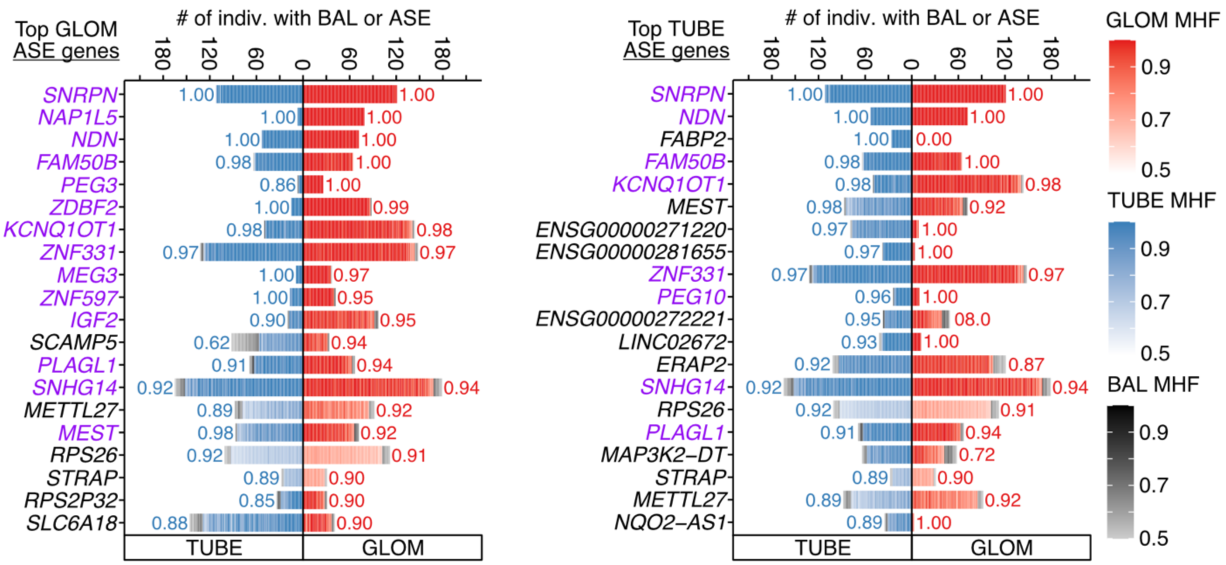


**Fig. S12. The top 20 ASE genes ranked by the proportion of individuals with ASE.** Same as main **Fig 1F**, but applied to the top 20 ASE genes in NEPTUNE GLOM (left) and TUBE (right), ordered by the proportion of individuals exhibiting ASE among those with COV for each gene (y-axis).

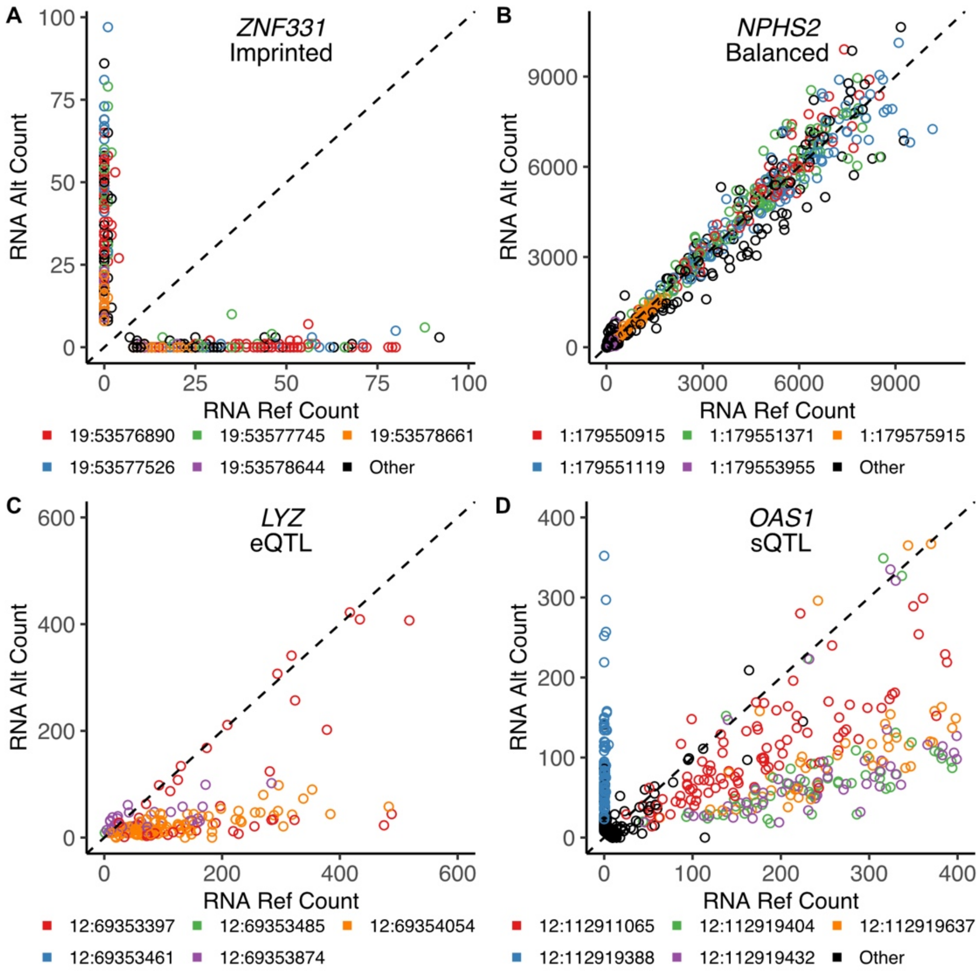


**Fig. S13. Allelic expression in example genes by ASE mechanism.** RNA Ref/Alt Count for all variants in (**A**) *ZNF331*, (**B**) *NPHS2*, (**C**) *LYZ*, and (**D**) *OAS1*, with RNA coverage ≥ 8 in NEPTUNE GLOM samples. For each gene, each dot is a variant-individual and the five most common variants by number of individuals are colored. *ZNF331* is a known imprinted gene, and *NPHS2* was identified as a biallelic/balanced expression gene. In (C), the variants 12:69353397 (red) and 12:69354054 (orange) are significant eQTLs whose reference allele significantly increases expression of *LYZ*. There are no sQTLs associated with *LYZ*. In (D), the variant 12:112919388 (blue) is a significant sQTL, with the alternative allele promoting retention of an intron that includes the variant itself, leading to monoallelic expression of the alternative allele. There are no eQTLs associated with *OAS1* and the axes are limited to 0-400.

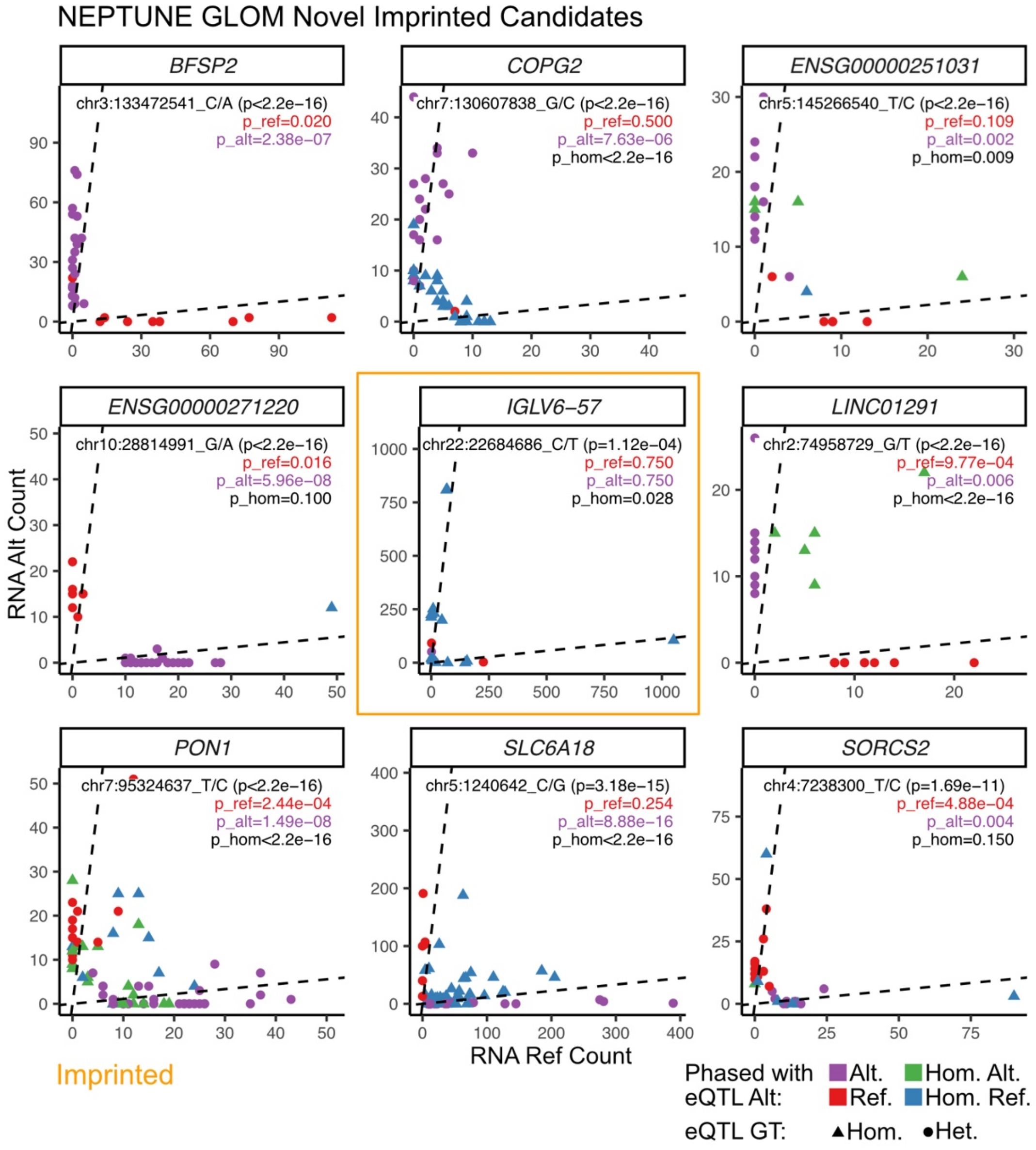


**Fig. S14. NEPTUNE GLOM RNA Ref/Alt Count for all variants in novel imprinted candidates, determined by running the Baran pipeline on NEPTUNE GLOM phASER data.** Each gene has an eQTL (sig. or non-sig.) denoted at the top with its eQTL p-value. Each dot is a variant-individual pair representing a heterozygous coding variant within the gene open reading frame. Coding variants (dots) are colored by the allele in phase with the eQTL alternative allele, e.g., purple if alternative coding variant is phased with alternative allele of the eQTL, green if the individual is homozygous alternative allele at the eQTL. The shape is determined by the genotype of the individual for the eQTL. Each panel shows the p-value for the three binomial tests described in **Methods**, with two-of-three significant binomial tests required to label a gene ASE mechanism as eQTL. Genes determined as imprinted have an orange border, all others have an eQTL ASE mechanism.

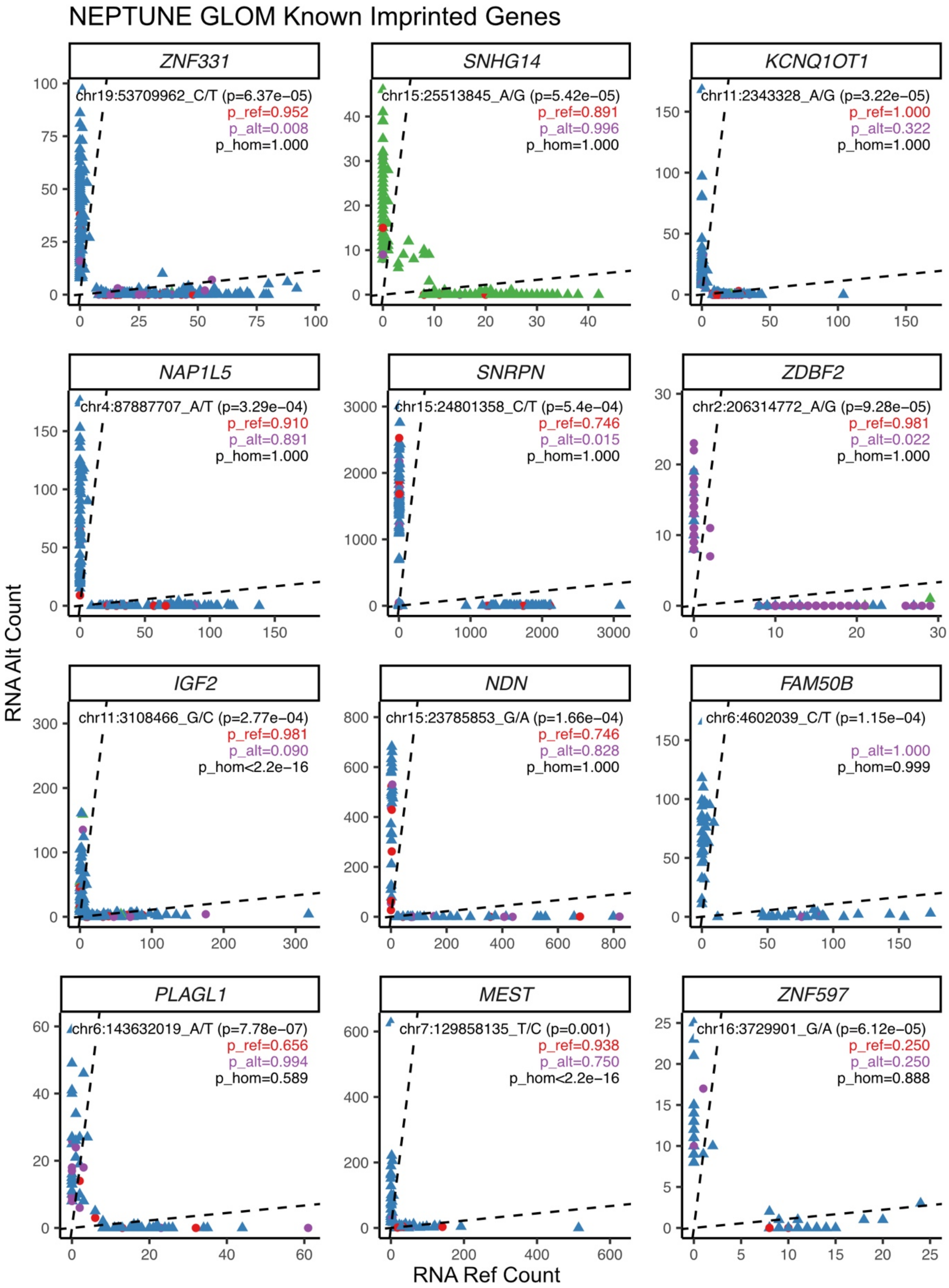
NEPTUNE GLOM Known Imprinted Genes
ZNF331
chr19:53709962_C/T (p=6.37e−05)
p_ref=0.952
p_alt=0.008
p_hom=1.000
SNHG14
chr15:25513845_A/G (p=5.42e−05)
p_ref=0.891
p_alt=0.996
p_hom=1.000
KCNQ1OT1
chr11:2343328_A/G (p=3.22e−05)
p_ref=1.000
p_alt=0.322
p_hom=1.000
NAP1L5
chr4:87887707_A/T (p=3.29e−04)
p_ref=0.910
p_alt=0.891
p_hom=1.000
SNRPN
chr15:24801358_C/T (p=5.4e−04)
p_ref=0.746
p_alt=0.015
p_hom=1.000
ZDBF2
chr2:206314772_A/G (p=9.28e−05)
p_ref=0.981
p_alt=0.022
p_hom=1.000
IGF2
chr11:3108466_G/C (p=2.77e−04)
p_ref=0.981
p_alt=0.090
p_hom<2.2e−16
NDN
chr15:23785853_G/A (p=1.66e−04)
p_ref=0.746
p_alt=0.828
p_hom=1.000
FAM50B
chr6:4602039_C/T (p=1.15e−04)
p_alt=1.000
p_hom=0.999
PLAGL1
chr6:143632019_A/T (p=7.78e−07)
p_ref=0.656
p_alt=0.994
p_hom=0.589
MEST
chr7:129858135_T/C (p=0.001)
p_ref=0.938
p_alt=0.750
p_hom<2.2e−16
ZNF597
chr16:3729901_G/A (p=6.12e−05)
p_ref=0.250
p_alt=0.250
p_hom=0.888
RNA Alt Count
RNA Ref Count

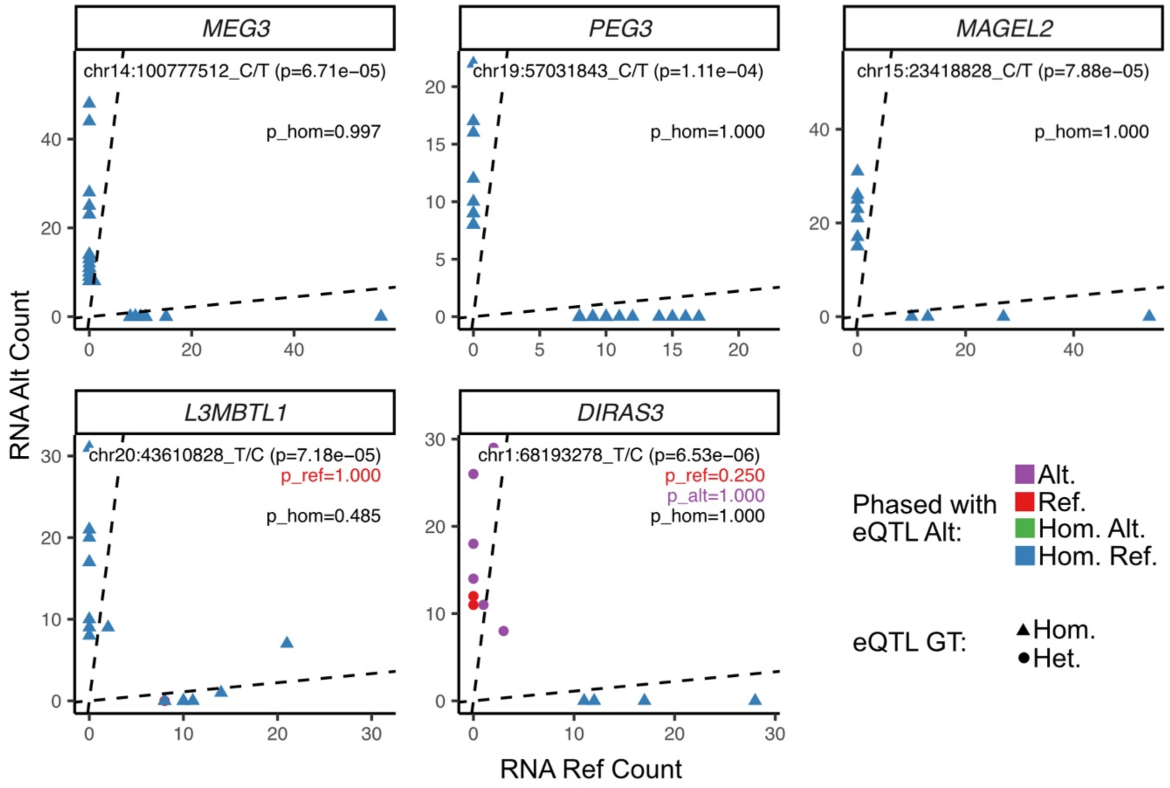


**Fig. S15. NEPTUNE GLOM RNA Ref/Alt Count for all variants in known imprinted genes, determined by running the Baran pipeline on NEPTUNE GLOM phASER data.** Same as **fig S14** but applied to 17 known imprinted genes in GLOM. All genes were determined to be imprinted.

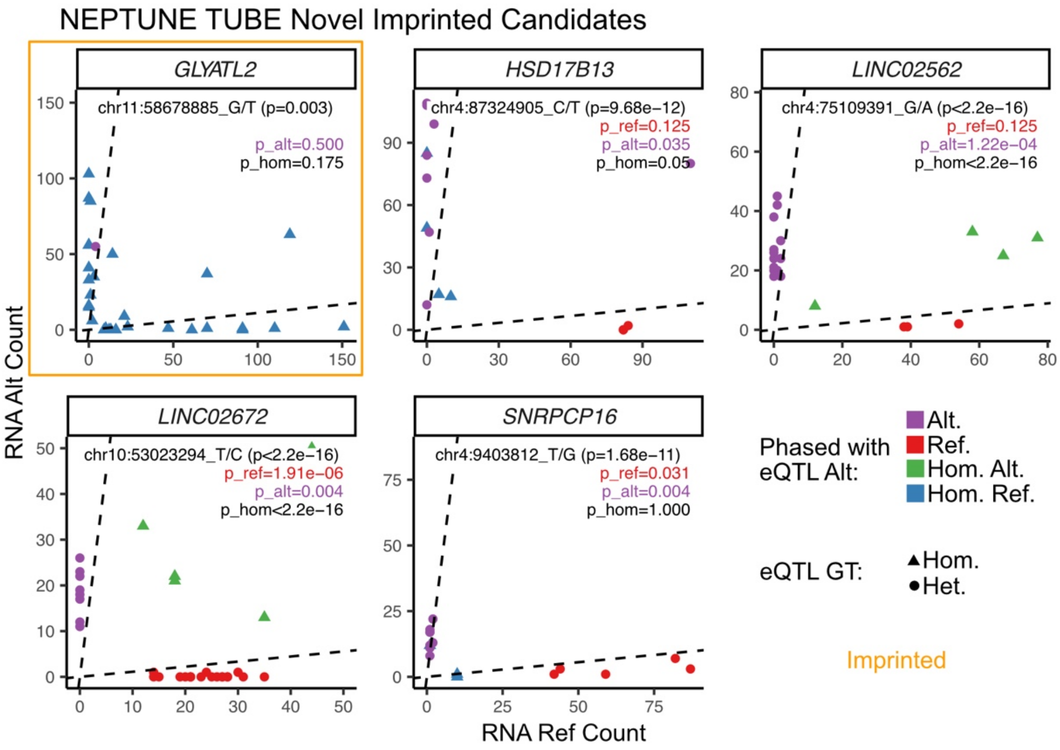


**Fig. S16. NEPTUNE TUBE RNA Ref/Alt Count for all variants in novel imprinted candidates, determined by running the Baran pipeline on NEPTUNE TUBE phASER data.** Same as **fig S14** but applied to 5 novel imprinted candidates in TUBE.

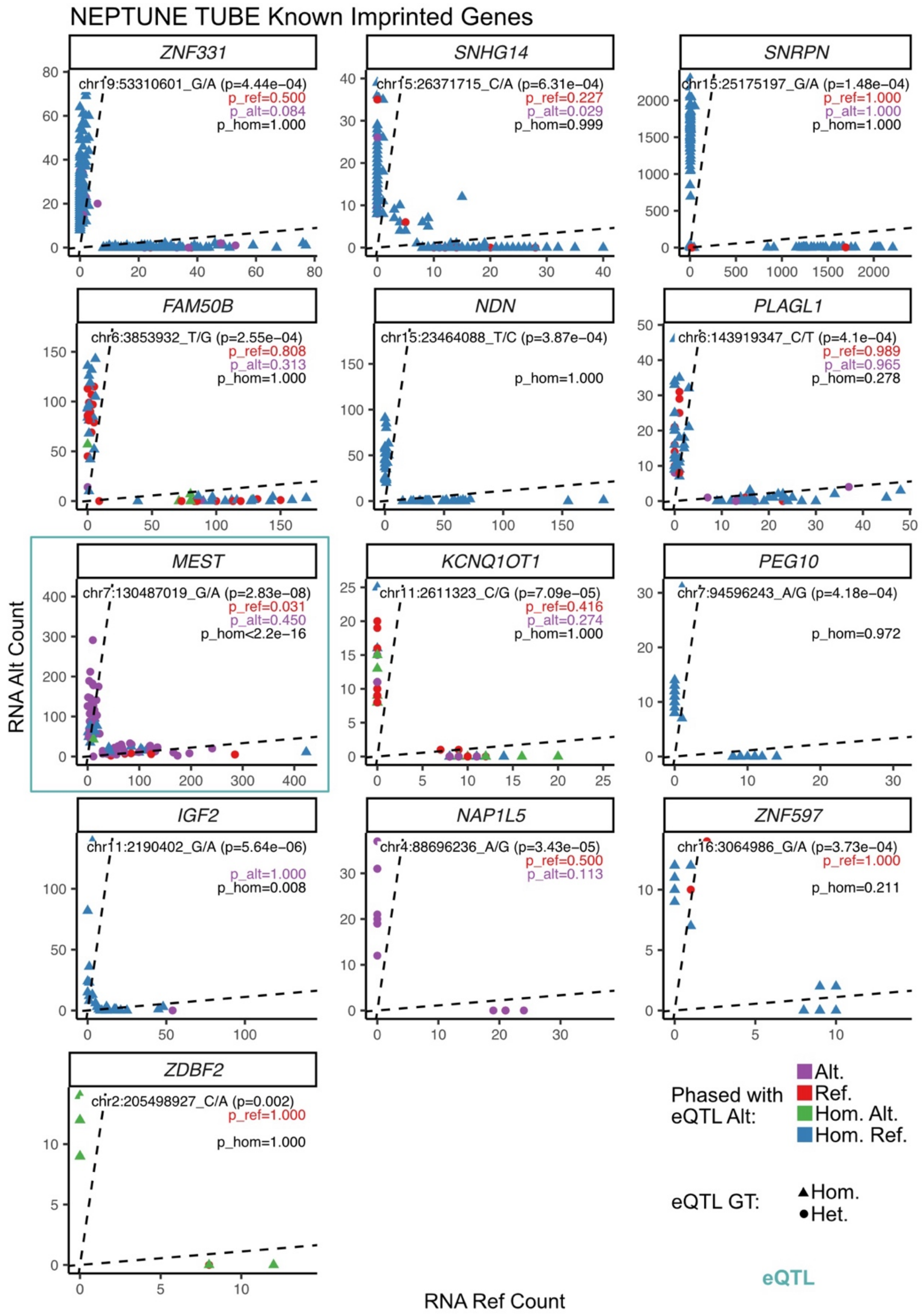


**Fig. S17. NEPTUNE TUBE RNA Ref/Alt Count for all variants in known imprinted genes, determined by running the Baran pipeline on NEPTUNE TUBE phASER data.** Same as **fig S14** but applied to 13 known imprinted genes in TUBE. Genes determined to have an eQTL mechanism have a teal border, all others were determined as imprinted.

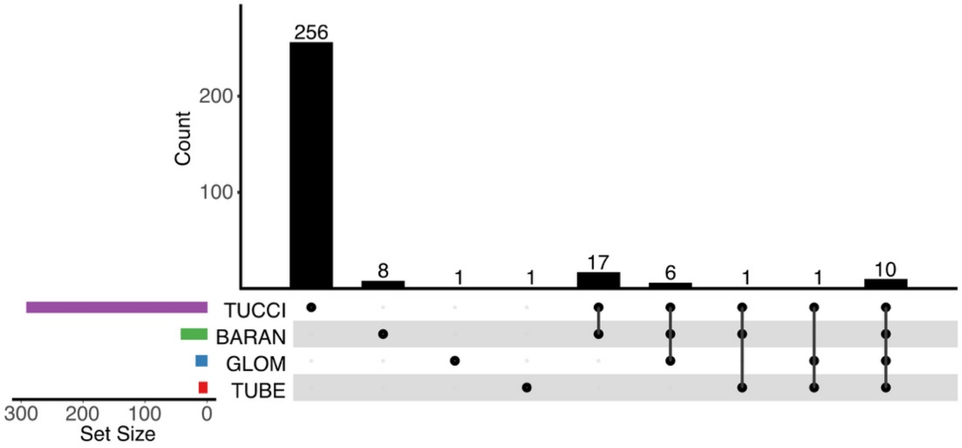


**Fig. S18. UpSet plot shows the number of imprinted genes within the intersection of different imprinted gene sets.** Tucci et al and Baran et al were combined to create our "known imprinted genes" list. GLOM and TUBE are the imprinted gene lists created by running the Baran pipeline on NEPTUNE data. Imprinted Gene Sets are ordered by number of genes: Tucci et al (n = 291), Baran et al (n = 42), NEPTUNE GLOM (n = 18), and NEPTUNE TUBE (n = 13).

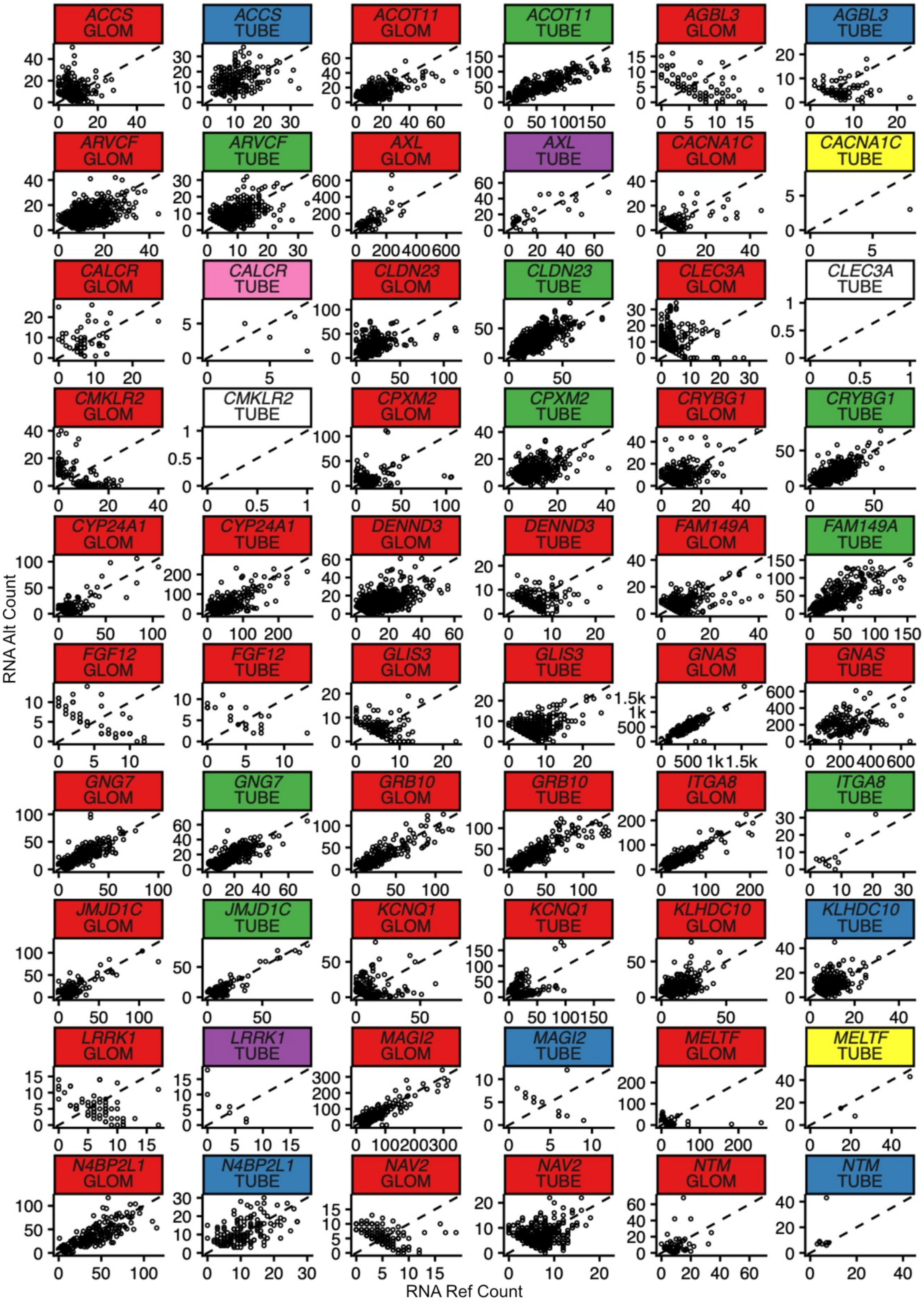
ACCS GLOM
ACCS TUBE
ACOT11 GLOM
ACOT11 TUBE
AGBL3 GLOM
AGBL3 TUBE
ARVCF GLOM
ARVCF TUBE
AXL GLOM
AXL TUBE
CACNA1C GLOM
CACNA1C TUBE
CALCR GLOM
CALCR TUBE
CLDN23 GLOM
CLDN23 TUBE
CLEC3A GLOM
CLEC3A TUBE
CMKLR2 GLOM
CMKLR2 TUBE
CPXM2 GLOM
CPXM2 TUBE
CRYBG1 GLOM
CRYBG1 TUBE
CYP24A1 GLOM
CYP24A1 TUBE
DENND3 GLOM
DENND3 TUBE
FAM149A GLOM
FAM149A TUBE
FGF12 GLOM
FGF12 TUBE
GLIS3 GLOM
GLIS3 TUBE
GNAS GLOM
GNAS TUBE
GNG7 GLOM
GNG7 TUBE
GRB10 GLOM
GRB10 TUBE
ITGA8 GLOM
ITGA8 TUBE
JMJD1C GLOM
JMJD1C TUBE
KCNQ1 GLOM
KCNQ1 TUBE
KLHDC10 GLOM
KLHDC10 TUBE
LRRK1 GLOM
LRRK1 TUBE
MAGI2 GLOM
MAGI2 TUBE
MELTF GLOM
MELTF TUBE
N4BP2L1 GLOM
N4BP2L1 TUBE
NAV2 GLOM
NAV2 TUBE
NTM GLOM
NTM TUBE
RNA Alt Count
RNA Ref Count

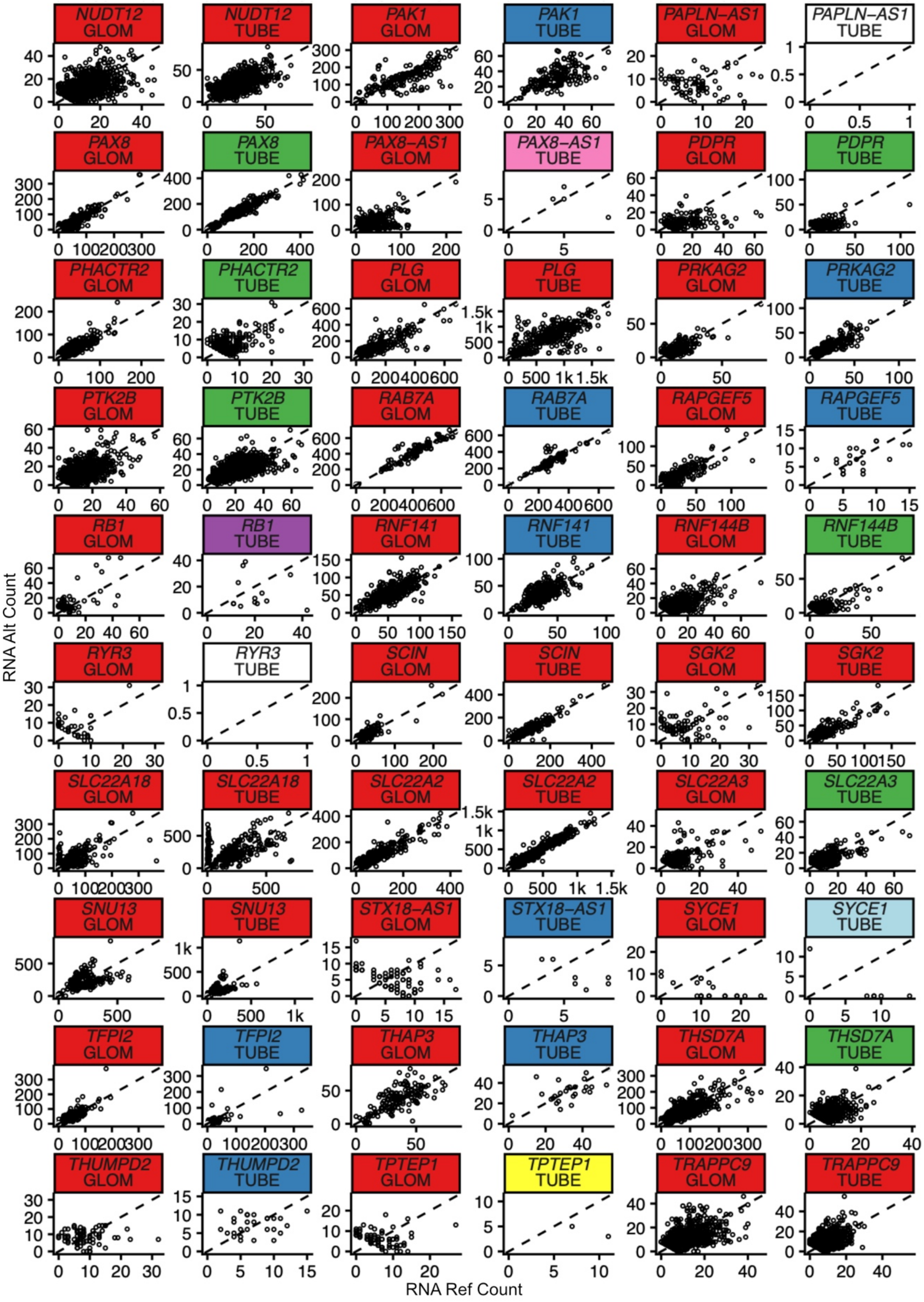
NUDT12 GLOM
NUDT12 TUBE
PAK1 GLOM
PAK1 TUBE
PAPLN-AS1 GLOM
PAPLN-AS1 TUBE
PAX8 GLOM
PAX8 TUBE
PAX8-AS1 GLOM
PAX8-AS1 TUBE
PDPR GLOM
PDPR TUBE
PHACTR2 GLOM
PHACTR2 TUBE
PLG GLOM
PLG TUBE
PRKAG2 GLOM
PRKAG2 TUBE
PTK2B GLOM
PTK2B TUBE
RAB7A GLOM
RAB7A TUBE
RAPGEF5 GLOM
RAPGEF5 TUBE
RB1 GLOM
RB1 TUBE
RNF141 GLOM
RNF141 TUBE
RNF144B GLOM
RNF144B TUBE
RYR3 GLOM
RYR3 TUBE
SCIN GLOM
SCIN TUBE
SGK2 GLOM
SGK2 TUBE
SLC22A18 GLOM
SLC22A18 TUBE
SLC22A2 GLOM
SLC22A2 TUBE
SLC22A3 GLOM
SLC22A3 TUBE
SNU13 GLOM
SNU13 TUBE
STX18-AS1 GLOM
STX18-AS1 TUBE
SYCE1 GLOM
SYCE1 TUBE
TFPI2 GLOM
TFPI2 TUBE
THAP3 GLOM
THAP3 TUBE
THSD7A GLOM
THSD7A TUBE
THUMPD2 GLOM
THUMPD2 TUBE
TPTEP1 GLOM
TPTEP1 TUBE
TRAPPC9 GLOM
TRAPPC9 TUBE
RNA Alt Count
RNA Ref Count

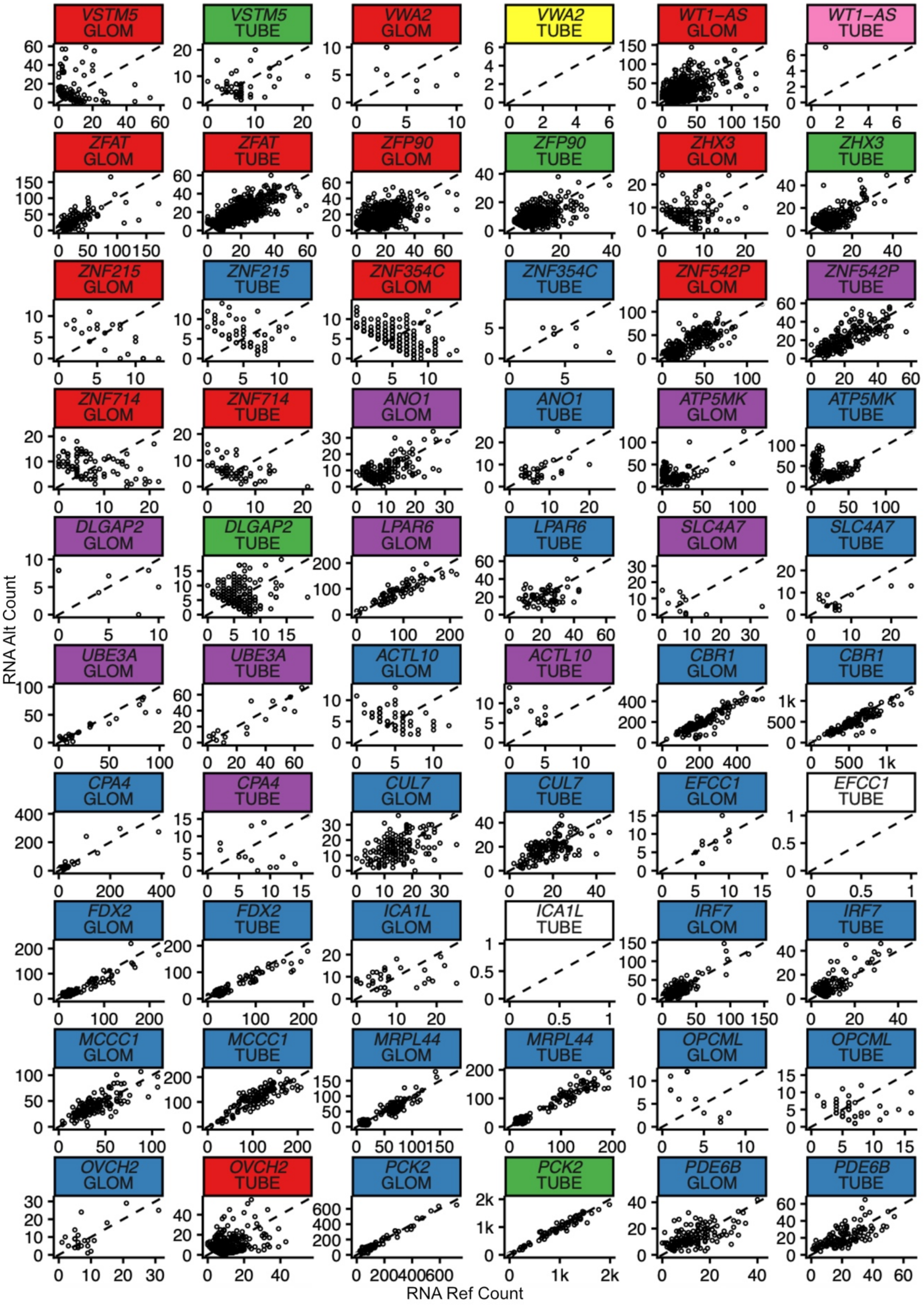
VSTM5 GLOM
VSTM5 TUBE
VWA2 GLOM
VWA2 TUBE
WT1-AS GLOM
WT1-AS TUBE
ZFAT GLOM
ZFAT TUBE
ZFP90 GLOM
ZFP90 TUBE
ZHX3 GLOM
ZHX3 TUBE
ZNF215 GLOM
ZNF215 TUBE
ZNF354C GLOM
ZNF354C TUBE
ZNF542P GLOM
ZNF542P TUBE
ZNF714 GLOM
ZNF714 TUBE
ANO1 GLOM
ANO1 TUBE
ATP5MK GLOM
ATP5MK TUBE
DLGAP2 GLOM
DLGAP2 TUBE
LPAR6 GLOM
LPAR6 TUBE
SLC4A7 GLOM
SLC4A7 TUBE
UBE3A GLOM
UBE3A TUBE
ACTL10 GLOM
ACTL10 TUBE
CBR1 GLOM
CBR1 TUBE
CPA4 GLOM
CPA4 TUBE
CUL7 GLOM
CUL7 TUBE
EFCC1 GLOM
EFCC1 TUBE
FDX2 GLOM
FDX2 TUBE
ICA1L GLOM
ICA1L TUBE
IRF7 GLOM
IRF7 TUBE
MCCC1 GLOM
MCCC1 TUBE
MRPL44 GLOM
MRPL44 TUBE
OPCML GLOM
OPCML TUBE
OVCH2 GLOM
OVCH2 TUBE
PCK2 GLOM
PCK2 TUBE
PDE6B GLOM
PDE6B TUBE
RNA Alt Count
RNA Ref Count

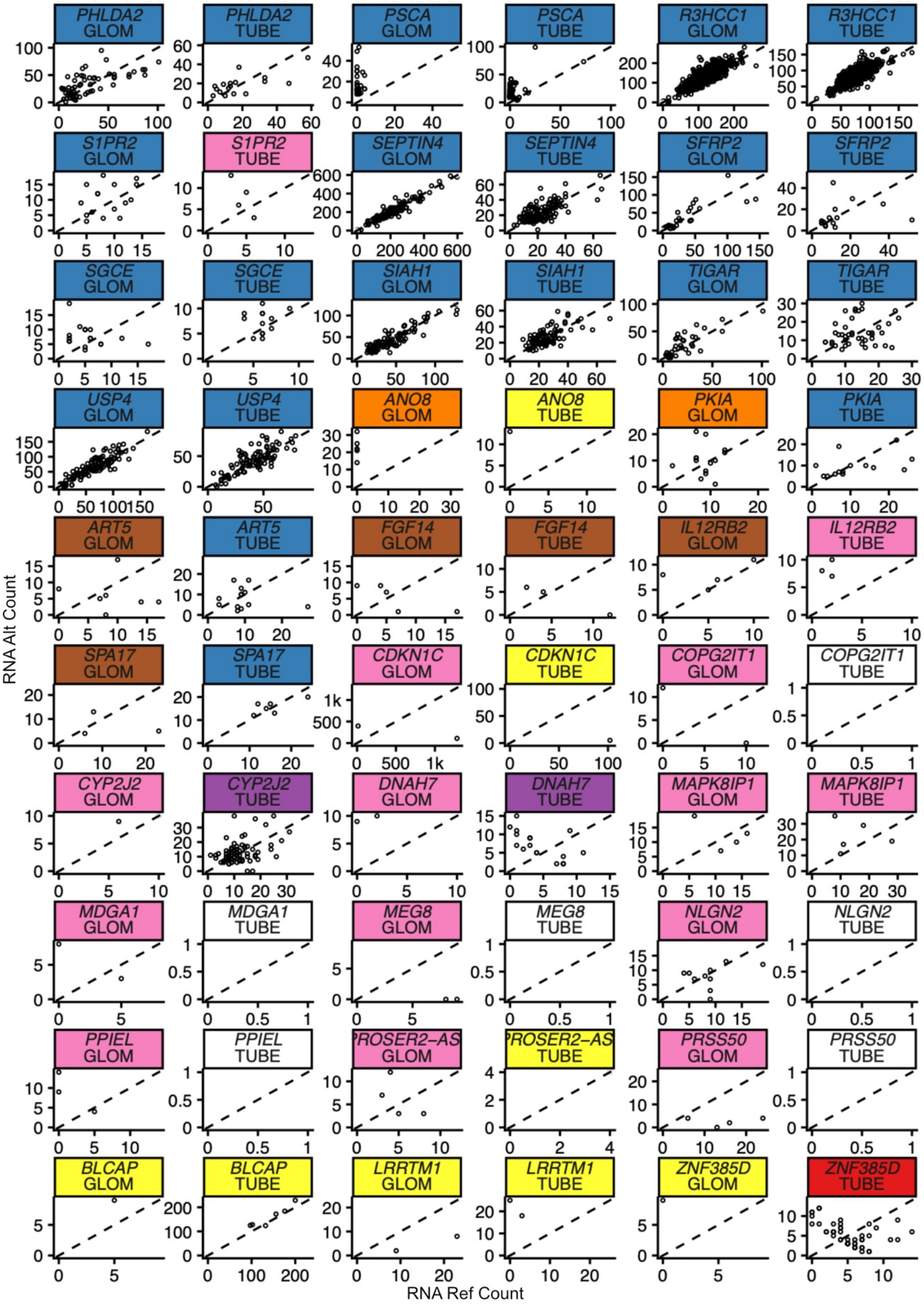
PHLDA2 GLOM
PHLDA2 TUBE
PSCA GLOM
PSCA TUBE
R3HCC1 GLOM
R3HCC1 TUBE
S1PR2 GLOM
S1PR2 TUBE
SEPTIN4 GLOM
SEPTIN4 TUBE
SFRP2 GLOM
SFRP2 TUBE
SGCE GLOM
SGCE TUBE
SIAH1 GLOM
SIAH1 TUBE
TIGAR GLOM
TIGAR TUBE
USP4 GLOM
USP4 TUBE
ANO8 GLOM
ANO8 TUBE
PKIA GLOM
PKIA TUBE
ART5 GLOM
ART5 TUBE
FGF14 GLOM
FGF14 TUBE
IL12RB2 GLOM
IL12RB2 TUBE
SPA17 GLOM
SPA17 TUBE
CDKN1C GLOM
CDKN1C TUBE
COPG2IT1 GLOM
COPG2IT1 TUBE
CYP2J2 GLOM
CYP2J2 TUBE
DNAH7 GLOM
DNAH7 TUBE
MAPK8IP1 GLOM
MAPK8IP1 TUBE
MDGA1 GLOM
MDGA1 TUBE
MEG8 GLOM
MEG8 TUBE
NLGN2 GLOM
NLGN2 TUBE
PPIEL GLOM
PPIEL TUBE
PROSER2-AS GLOM
PROSER2-AS TUBE
PRSS50 GLOM
PRSS50 TUBE
BLCAP GLOM
BLCAP TUBE
LRRTM1 GLOM
LRRTM1 TUBE
ZNF385D GLOM
ZNF385D TUBE
RNA Alt Count
RNA Ref Count

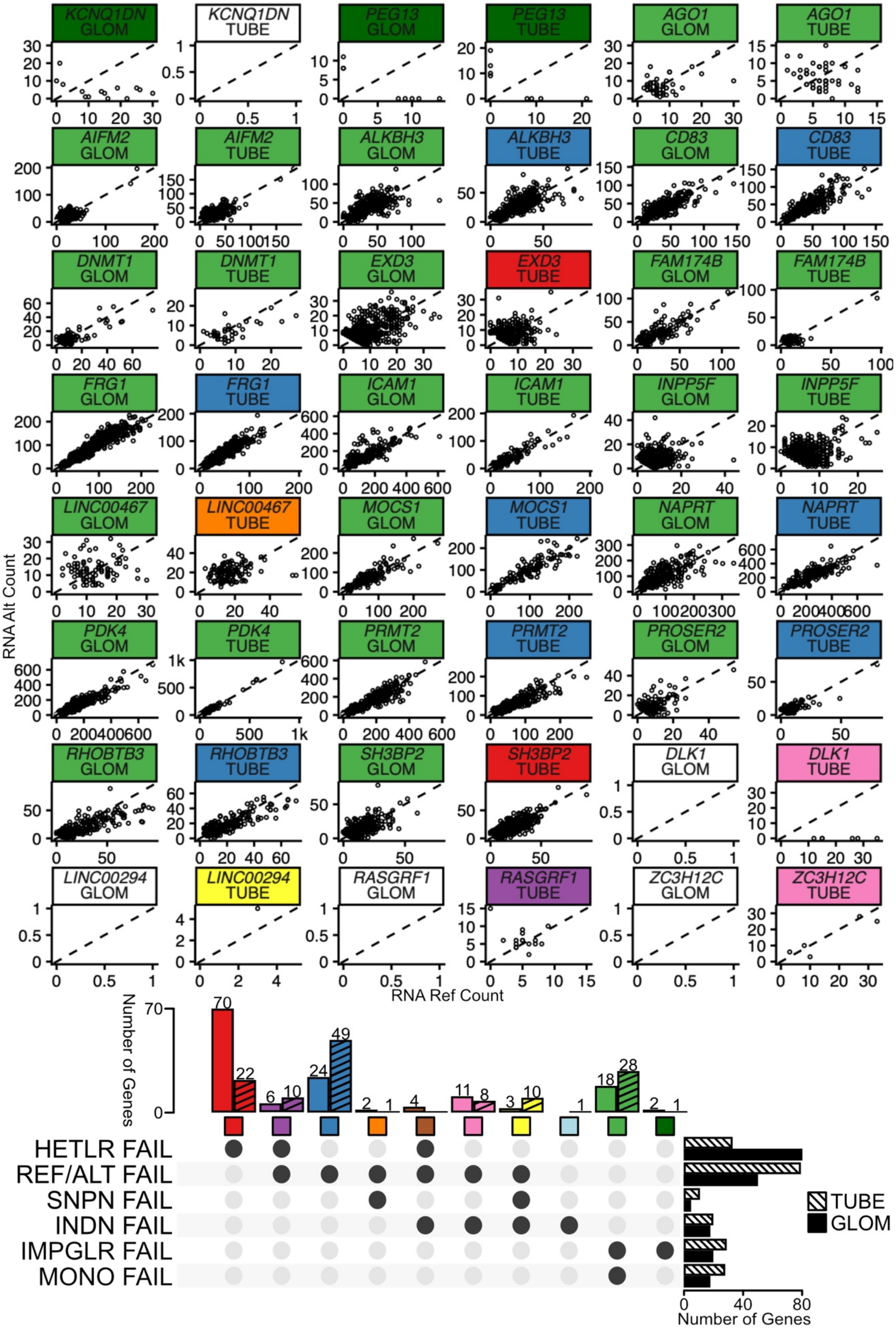

KCNQ1DN GLOM
KCNQ1DN TUBE
PEG13 GLOM
PEG13 TUBE
AGO1 GLOM
AGO1 TUBE
AIFM2 GLOM
AIFM2 TUBE
ALKBH3 GLOM
ALKBH3 TUBE
CD83 GLOM
CD83 TUBE
DNMT1 GLOM
DNMT1 TUBE
EXD3 GLOM
EXD3 TUBE
FAM174B GLOM
FAM174B TUBE
FRG1 GLOM
FRG1 TUBE
ICAM1 GLOM
ICAM1 TUBE
INPP5F GLOM
INPP5F TUBE
LINC00467 GLOM
LINC00467 TUBE
MOCS1 GLOM
MOCS1 TUBE
NAPRT GLOM
NAPRT TUBE
PDK4 GLOM
PDK4 TUBE
PRMT2 GLOM
PRMT2 TUBE
PROSER2 GLOM
PROSER2 TUBE
RHOBTB3 GLOM
RHOBTB3 TUBE
SH3BP2 GLOM
SH3BP2 TUBE
DLK1 GLOM
DLK1 TUBE
LINC00294 GLOM
LINC00294 TUBE
RASGRF1 GLOM
RASGRF1 TUBE
ZC3H12C GLOM
ZC3H12C TUBE
RNA Alt Count
RNA Ref Count
Number of Genes
70
22
6
10
24
49
2
1
4
11
8
3
10
1
18
28
2
1
HETLR FAIL
REF/ALT FAIL
SNPN FAIL
INDN FAIL
IMPGLR FAIL
MONO FAIL
TUBE
GLOM
0
40
80
Number of Genes

**Fig. S19. NEPTUNE GLOM and TUBE RNA Ref/Alt Count for all variants in known imprinted genes that were not found as imprinted by running the Baran pipeline on NEPTUNE data.** Each dot represents a variant-individual with RNA coverage ≥ 8 and HWE p-value ≥ 0.001. Panels are stratified by gene and NEPTUNE tissue (GLOM/TUBE), then colored by the reason the gene was not labelled as imprinted, as shown in the UpSet plot key at the bottom. Genes that lacked sufficient data to consider imprinted status had hetlr ≥ 1 (HETLR FAIL), no variant with monoallelic reference expression and monoallelic alternative expression in different individuals (REF/ALT FAIL), < 2 variants (SNPN FAIL), < 5 individuals (INDN FAIL). "Considered genes" that lacked evidence of imprinting had impglr ≤ 40 (IMPGLR FAIL) or < 80% of variant-individuals within the gene were expressed monoallelically (MONO FAIL). Panel labels with no color represent genes with no data in that tissue.

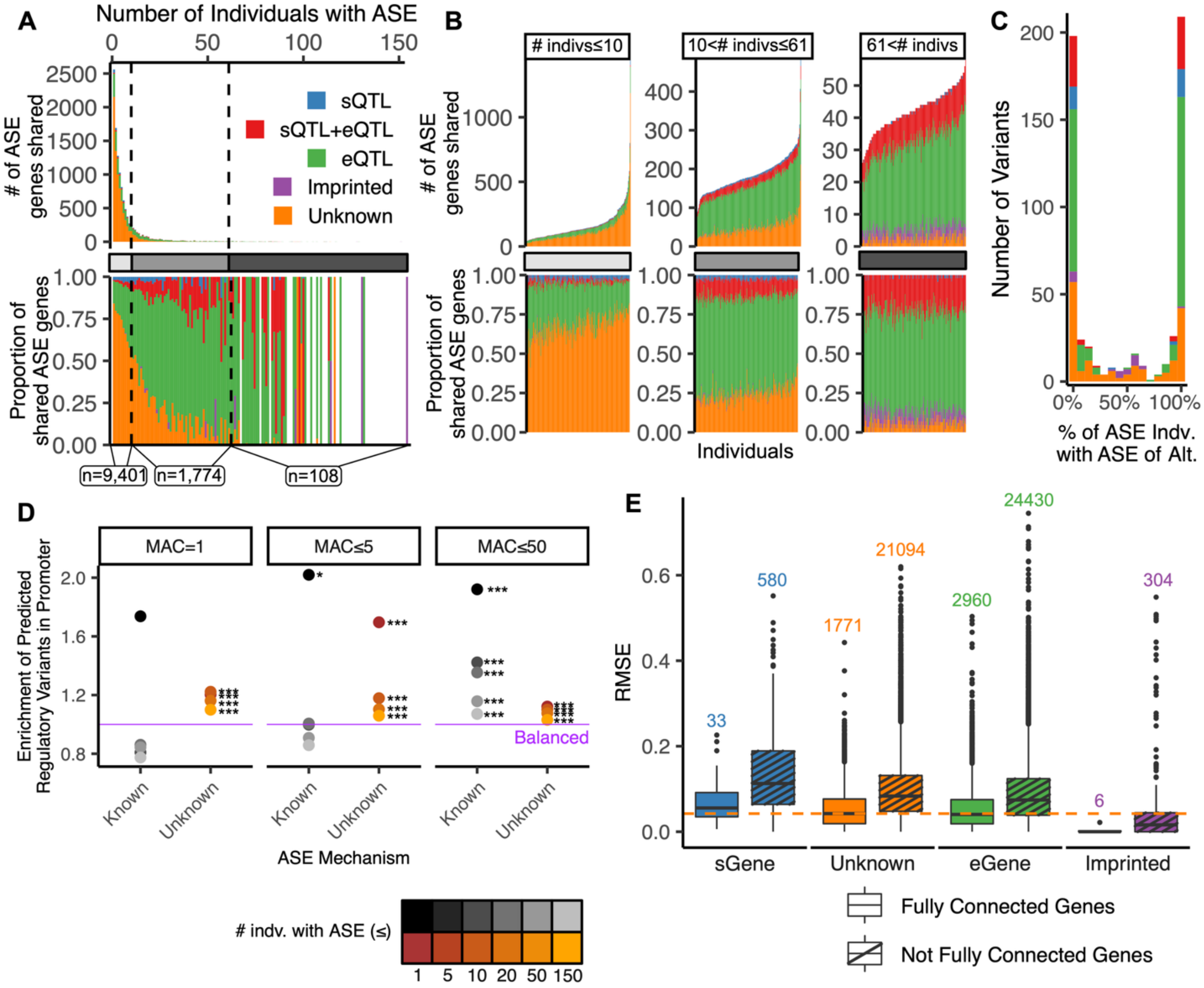


**Fig. S20. Molecular mechanisms of ASE in NEPTUNE TUBE** Same as **(A) Fig 2B, (B) Fig 2C**, (**C**) **Fig 2D**, **(D) Fig2F**, and **(E) Fig 3B,** but applied to NEPTUNE TUBE. In (E), pairwise Wilcoxon test showed that all groups had significant mean differences ($P < 0.001$) except Unknown and eGene ($P = 0.34$).

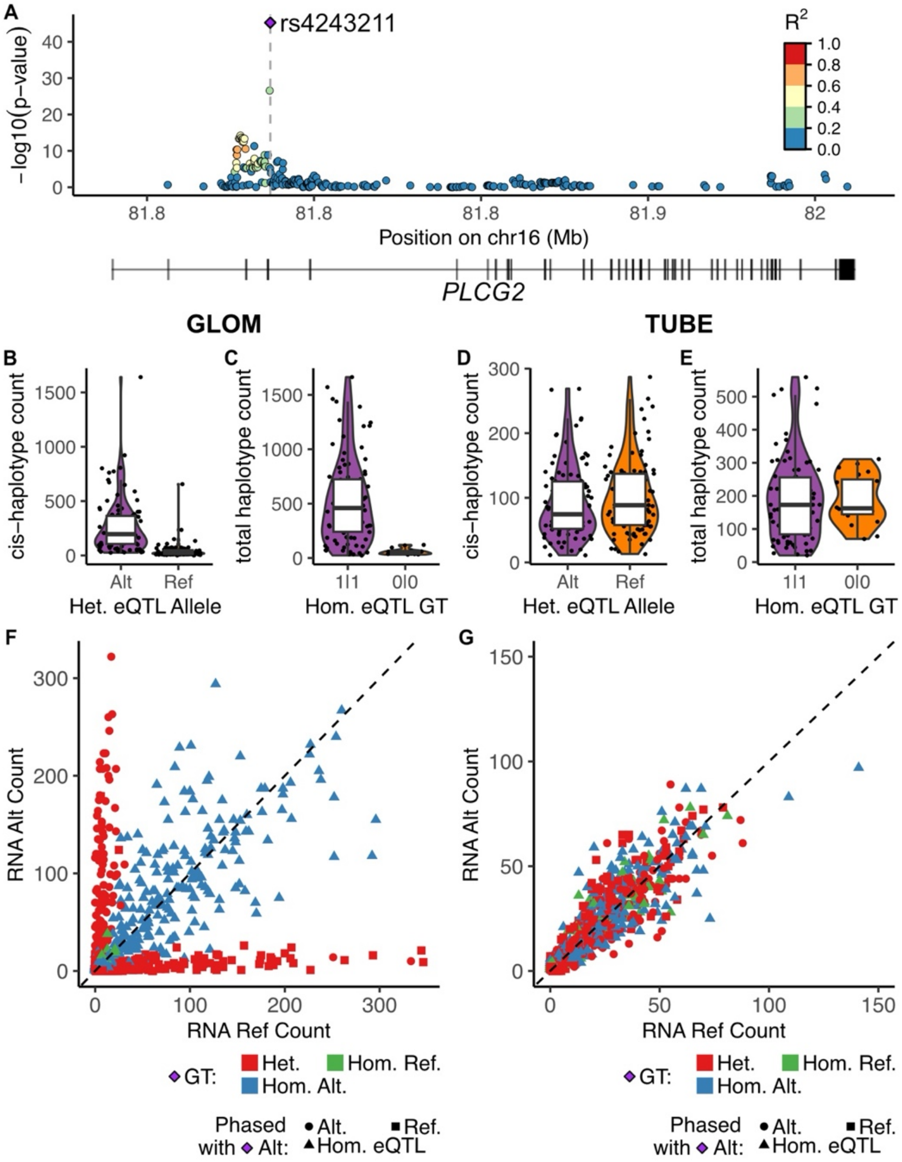


**Fig. S21. A GLOM-specific eQTL drives ASE of *PLCG2*. (A)** eQTL p-values of *PLCG2* variants in GLOM samples. Variants are colored by $R^2$ with rs4243211 (purple), the most significant eQTL in GLOM. *PLCG2* gene annotation is shown below the plot. **(B)** Haplotypic count of the PLCG2 haplotype in cis with the alternative (Alt) or reference (Ref) allele of the rs4243211 eQTL among all individuals heterozygous for rs4243211. Each dot is a GLOM sample. **(C)** Total haplotypic count (haplotype A count + haplotype B count) for *PLCG2* from phASER Gene AE among individuals homozygous for the rs4243211 eQTL. Each dot is a GLOM sample, stratified by rs4243211 genotype. **(D)** Same as (B), using TUBE samples. **(E)** Same as (C), using TUBE samples. **(F)** RNA Allelic Counts in all GLOM samples for all variants in *PLCG2*. Each dot is a variant in an individual. Variants are colored by the genotype of rs4243211 in the individual, the most significant eVariant in *PLCG2* in GLOM (see A). The shape of the variant point is determined by the allele (reference or alternative) that was phased on the same haplotype as the alternative allele of rs4243211, the allele that increases *PLCG2* expression. **(G)** Same as (F), using TUBE samples.

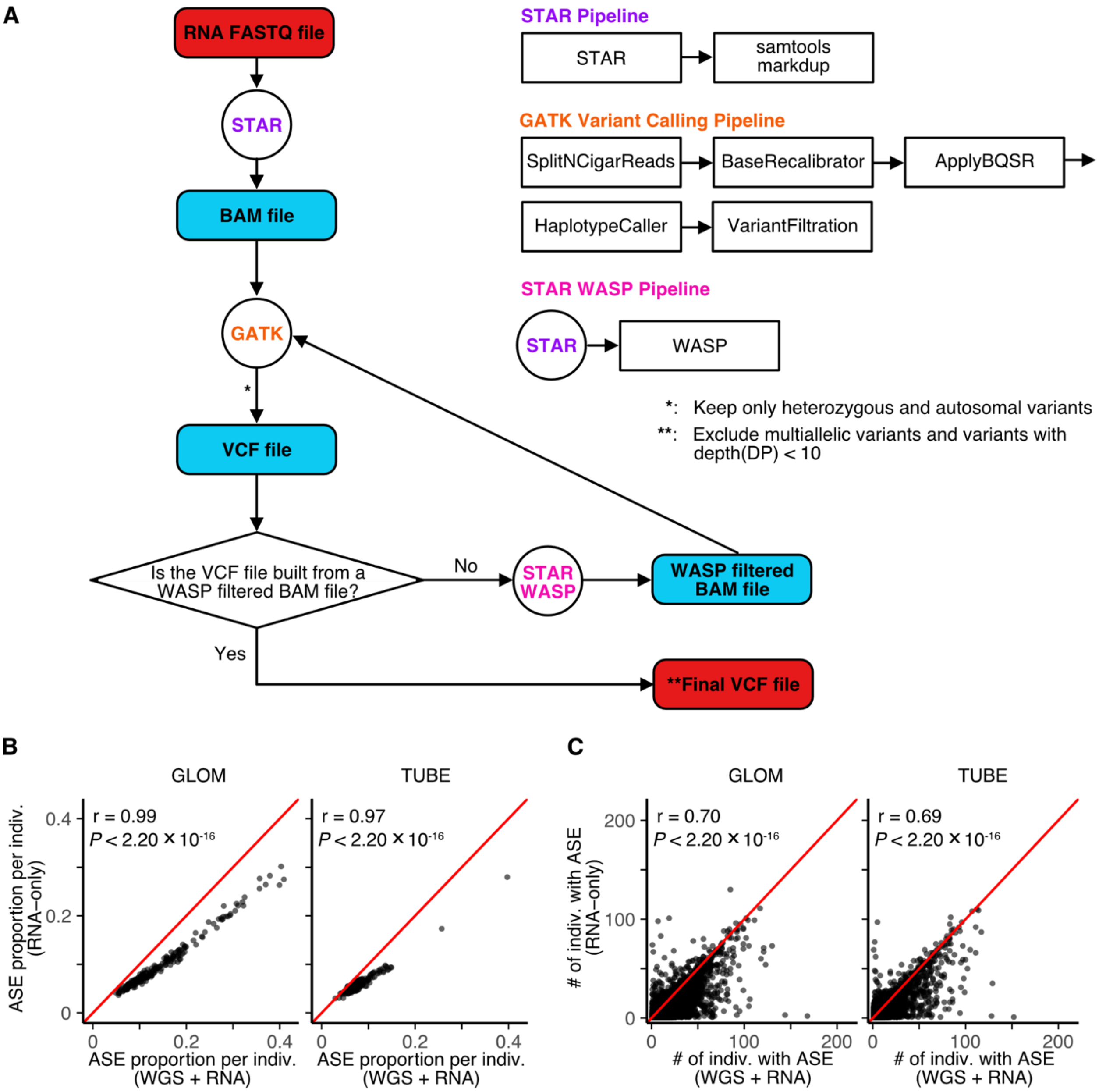


**Fig. S22. ASE pipeline using RNA-only data. (A)** Schematic workflow for variant calling from RNA-seq FASTQ data. **(B)** Comparison of ASE proportion per individual between the WGS+RNA and RNA-only methods. Each dot represents an individual with ASE proportions. Pearson correlation analysis shows a strong correlation in ASE proportion per individual between the two approaches. The red line represents the identity line (y = x). **(C)** Comparison of the number of individuals in which each gene was identified as ASE using WGS+RNA versus RNA-only. Each dot represents a gene identified as ASE in at least one individual by both methods. Spearman correlation analysis reveals a high concordance in frequency of ASE genes across individuals between the two methods.

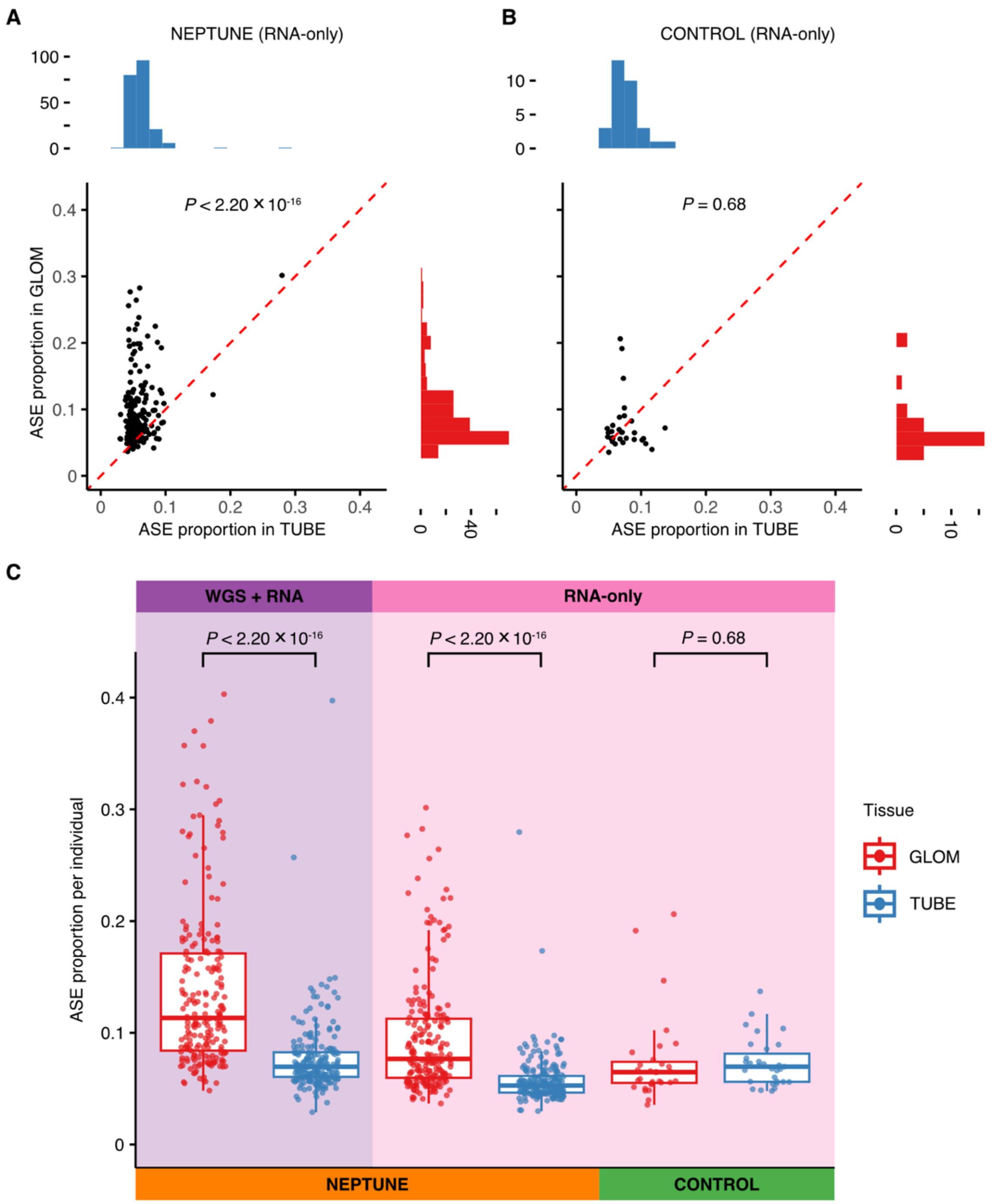


**Fig. S23. ASE proportion per individual in NEPTUNE and control.** (**A**) ASE proportion per individual in paired NEPTUNE samples (n = 206), in the RNA-only ASE analysis pipeline (using RNA-seq data alone). Each dot represents an individual, with ASE proportion in TUBE (x-axis) and GLOM (y-axis). Marginal histograms show ASE proportion distributions in GLOM (red) and TUBE (blue). (**B**) ASE proportion per individual in paired CONTROL samples (n = 31), in the RNA-only pipeline. (**C**) ASE proportion per individual across cohorts. The top panel indicates the method used: purple for WGS+RNA pipeline and pink for RNA-only pipeline. The bottom panel shows the cohort: paired NEPTUNE samples (n = 206) and paired control samples (n = 31). Statistical significance was assessed using the Wilcoxon signed-rank test.

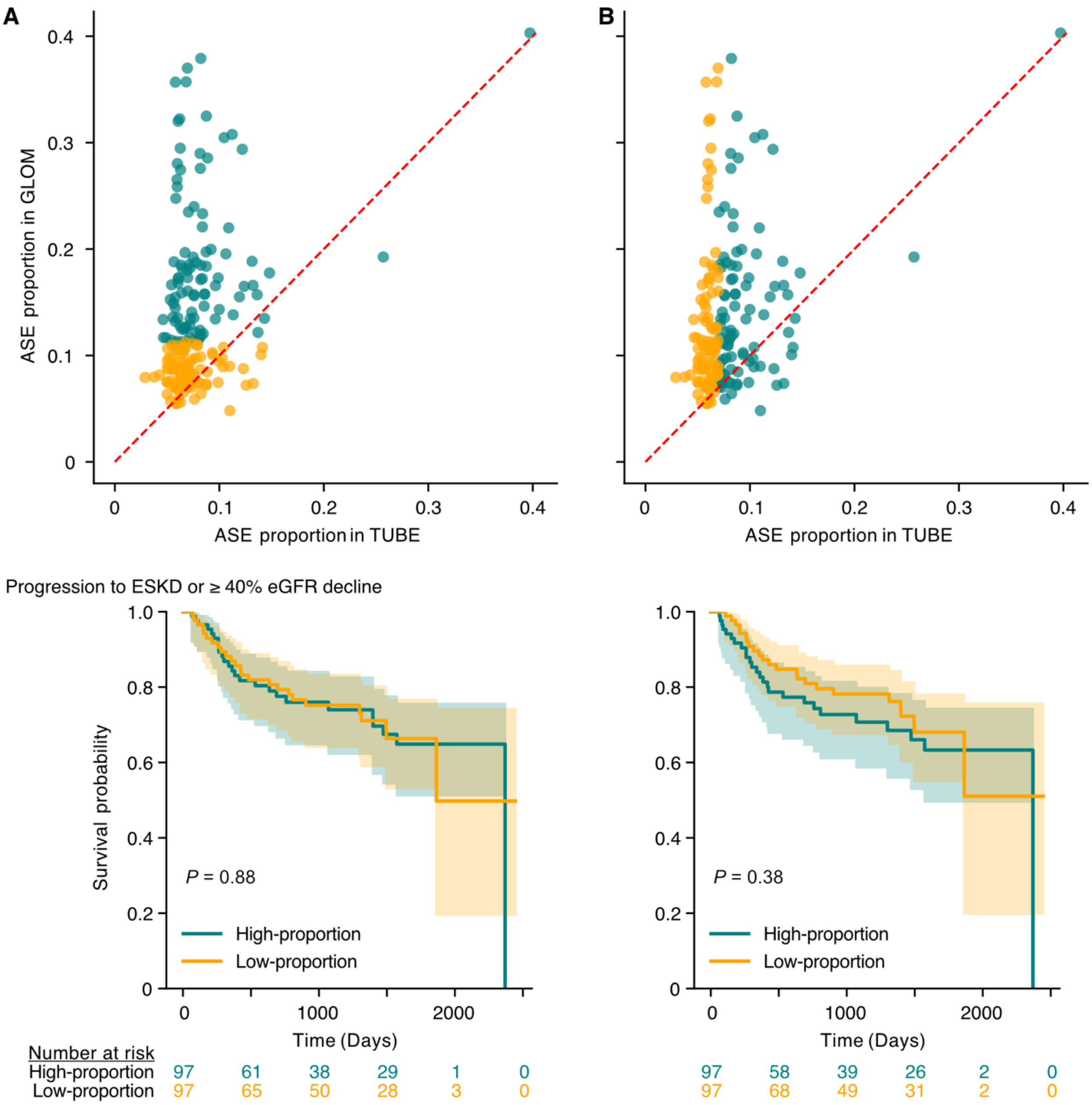


**Fig. S24. Kaplan–Meier survival analysis of NEPTUNE samples stratified by ASE proportion.** (**A**) The top scatterplot shows the samples with GLOM and TUBE ASE proportion, with samples divided into two equal groups based on GLOM ASE proportion. The ASE proportion of the high ASE group ranged from 0.403 to 0.113, and the low ASE group ranged from 0.112 to 0.048. The bottom plot shows Kaplan-Meier survival analysis representing the progression to the ESKD and/or ≥ 40% eGFR decline. (**B**) The same analysis is shown for TUBE. Samples were also equally divided into two groups based on TUBE ASE proportion. The ASE proportion of high ASE group ranged from 0.397 to 0.070, and the low ASE group ranged from 0.070 to 0.029.

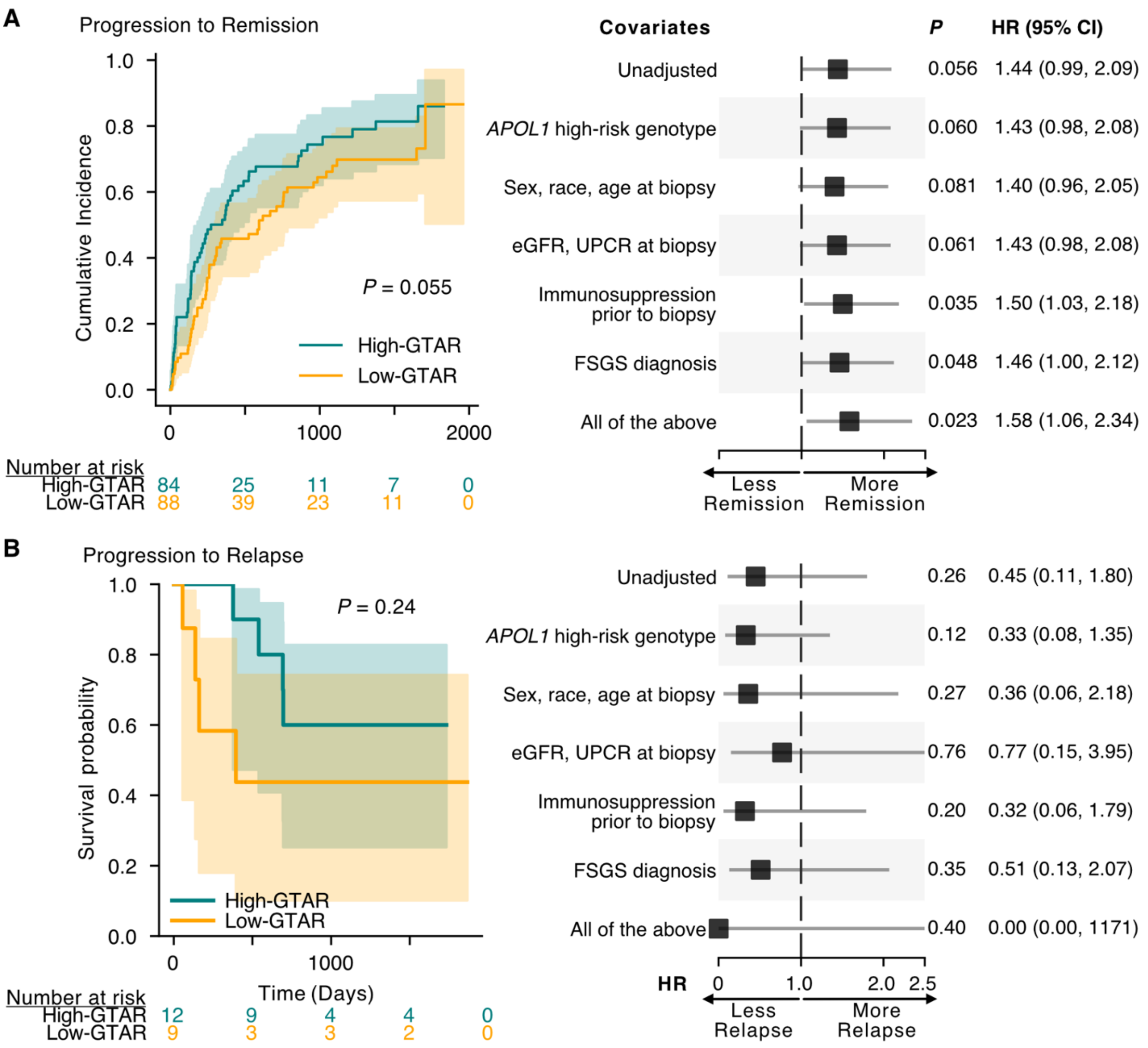


**Fig. S25. Survival analysis using clinical outcomes of remission and relapse.** (**A**) Remission-based analysis: Kaplan–Meier survival curves (left) and Cox proportional hazards models adjusting for different potential confounders (right). Here, the event of interest is remission, which represents a favorable clinical outcome. (**B**) Relapse-based analysis: Kaplan–Meier survival curves (left) and Cox proportional hazards models adjusting for different potential confounders (right). Here, the event of interest is relapse, which represents an unfavorable clinical outcome.

**Table S1.** Demographic and clinical characteristics of the NEPTUNE and control cohorts.

**Table S2.** Summary result of TOGA.

**Table S3.** Summary statistics of heterozygous variants in NEPTUNE.

**Table S4.** Summary statistics of ASE analysis by cohort and method.

**Table S5.** GLOM and TUBE ASE genes observed across all individuals with the number of samples.

**Table S6.** List of 299 known imprinted genes curated through literature review.

**Table S7.** Cell proportion and normalized expression of GLOM/TUBE ASE genes using KPMP single cell RNA-seq data. The grouping of original KPMP cell types is also included.

**Table S8.** Statistical comparisons between GLOM and TUBE ASE proportion and clinical phenotypes. Results are presented for paired NEPTUNE samples (first tab), paired NEPTUNE samples used in survival analysis (second tab), and CONTROL samples (third tab).

**Table S9.** Differentially expressed genes identified between high and low GTAR groups based on RNA expression in GLOM and TUBE.

**Table S10**. Detailed characterization of 181 DEGs.

**Table S11.** Significantly enriched pathways and gene ontology (GO) terms comparing high and low GTAR groups. Both gene set enrichment analysis and over-representation analysis of significantly upregulated and downregulated genes from DESeq2 are included.

### Members of the Nephrotic Syndrome Study Network (NEPTUNE)

**NEPTUNE Collaborating Sites**

*Atrium Health Levine Children's Hospital, Charlotte, SC*: Susan Massengill[*], Layla Lo[#]
*Cleveland Clinic, Cleveland, OH*: Katherine Dell[*], John O'Toole[*], John Sedor[**], Victoria Grange[#]
*Children's Hospital, Denver, CO:* Bradley Dixon[*], Nathan Rogers[#]
*Children's Hospital, Los Angeles, CA*: Rachel Lestz[*], Natalie Esquivias[#]
*Children's Mercy Hospital, Kansas City, MO*: Tarak Srivastava[*], Kelsey Markus[#]
*Cohen Children's Hospital, New Hyde Park, NY*: Christine Sethna[*], Suzanne Vento[#]
*Columbia University, New York, NY:* Pietro Canetta[*]
*Duke University Medical Center, Durham, NC:* Opeyemi Olabisi[*], Rasheed Gbadegesin[**], Kimberly Cicio[#]
*Emory University, Atlanta, GA:* Laurence Greenbaum[*], Chia-shi Wang[*], Chris Fan[#]
*The Lundquist Institute, Torrance, CA:* Sharon Adler[*], Janine LaPage[#]
*John H Stroger Cook County Hospital, Chicago, IL:* Amatur Amarah[*]
*Johns Hopkins Medicine, Baltimore, MD:* Meredith Atkinson[*], Ryan Hutson[#]
*Mayo Clinic, Rochester, MN:* John Lieske, Marie Hogan, Fernando Fervenza
*Medical University of South Carolina, Charleston, SC:* David Selewski[*], Cheryl Alston[#]
*Montefiore Medical Center, Bronx, NY:* Kim Reidy[*], Michael Ross[*], Frederick Kaskel[**], Patricia Flynn[#]
*New York University Medical Center, New York, NY:* Laura Malaga-Dieguez[*], Olga Zhdanova[**], Laura Jane Pehrson[#], Melanie Miranda[#]
*The Ohio State University College of Medicine, Columbus, OH*: Salem Almaani[*], Laci Roberts[#]
*Riley Children's Hospital of Indiana University, Indianapolis, IN:* Myda Khalid[*], Veronica Servin[#]
*Stanford University, Stanford, CA:* Richard Lafayette[*], Elizabeth Chen[#]
*Temple University, Philadelphia, PA:* Iris Lee[**]
*Texas Children's Hospital at Baylor College of Medicine, Houston, TX*: Shweta Shah[*], Thinh Phan[#]
*University Health Network Toronto:* Heather Reich[*], Michelle Hladunewich[**], Paul Ling[#], Martin Romano[#]
*University of California at San Diego, San Diego, CA:* Ambarish Athavale[*], Caitlin Carter[*], Kristin Zeeb[#]
*University of California at San Francisco, San Francisco, CA*: Paul Brakeman[*], Daniel Schrader
*University of Colorado Anschutz Medical Campus, Aurora, CO*: James Dylewski[*] Nathan Rogers[#]
*University of Kansas Medical Center, Kansas City, KS*: Ellen McCarthy[*], Catherine Creed[#]
*University of Miami, Miami, FL:* Alessia Fornoni[*], Miguel Bandes[#]
*University of Michigan, Ann Arbor, MI:* Matthias Kretzler[*], Laura Mariani[*], Zubin Modi[*], Amanda Williams[#], Roxy Ni[#]
*University of Minnesota, Minneapolis, MN:* Patrick Nachman[*], Michelle Rheault[*], Ariel Langenberger[#], Brady Wallner[#]
*University of North Carolina, Chapel Hill, NC:* Vimal Derebail[*], Keisha Gibson[*], Anne Froment[#], Sharia Warren[#]
*University of Pennsylvania, Philadelphia, PA:* Lawrence Holzman[*], Kevin Meyers[**], Krishna Kallem[#], Arielle Swenson[#]

*University of Texas San Antonio, San Antonio, TX*: Samin Sharma**
*University of Texas Southwestern, Dallas, TX:* Elizabeth Roehm*, Kamalanathan Sambandam**, Elizabeth Brown**
*University of Washington, Seattle, WA:* Ashley Jefferson*, Sangeeta Hingorani**, Katherine Tuttle**§, Linda Manahan #, Emily Pao#, Kelli Kuykendall§
*Wake Forest University Baptist Health, Winston-Salem, NC:* Jen Jar Lin**
*Washington University in St. Louis, St. Louis, MO*: Brian Stotter*, Joseph Dumayas#

**Data Analysis and Coordinating Center:** *University of Michigan:* Matthias Kretzler*, Brenda Gillespie**, Laura Mariani**, Zubin Modi**, Eloise Salmon**, Howard Trachtman**, Hailey Desmond, Sean Eddy, Damian Fermin, Wenjun Ju, Maria Larkina, Chrysta Lienczewski, Rebecca Scherr, Jonathan Troost, Amanda Williams, Yan Zhai; *Cleveland Clinic:* Crystal Gadegbeku**, John Sedor**, *Duke University:* Laura Barisoni**; *Harvard University:* Matthew G Sampson**; *Northwestern University:* Abigail Smith**; *University of Pennsylvania:* Lawrence Holzman**, Jarcy Zee**

**Digital Pathology Committee:** Carmen Avila-Casado *(University Health Network)*, Serena Bagnasco *(Johns Hopkins University)*, Lihong Bu *(Mayo Clinic)*, Shelley Caltharp *(Emory University)*, Clarissa Cassol *(Arkana)*, Dawit Demeke *(University of Michigan)*, Brenda Gillespie *(University of Michigan)*, Jared Hassler *(Temple University)*, Leal Herlitz *(Cleveland Clinic)*, Stephen Hewitt *(National Cancer Institute)*, Jeff Hodgin *(University of Michigan)*, Danni Holanda *(Arkana)*, Neeraja Kambham *(Stanford University)*, Kevin Lemley, Laura Mariani *(University of Michigan)*, Nidia Messias *(Washington University)*, Alexei Mikhailov *(Wake Forest)*, Vanessa Moreno *(University of North Carolina)*, Behzad Najafian *(University of Washington)*, Matthew Palmer *(University of Pennsylvania)*, Avi Rosenberg *(Johns Hopkins University)*, Virginie Royal *(University of Montreal)*, Miroslav Sekulik *(Columbia University)*, Barry Stokes *(Columbia University)*, David Thomas *(Duke University)*, Ming Wu *(University of New York)*, Michifumi Yamashita *(Cedar Sinai)*, Hong Yin *(Emory University)*, Jarcy Zee *(University of Pennsylvania)*, Yiqin Zuo *(University of Miami)*. Co-Chairs: Laura Barisoni *(Duke University)*, Cynthia Nast *(Cedar Sinai)*.
